# Foxg1 regulates translation of neocortical neuronal genes, including the main NMDA receptor subunit gene, *Grin1*

**DOI:** 10.1101/2022.10.05.510986

**Authors:** Osvaldo Artimagnella, Mauro Esposito, Elena Sabina Maftei, Remo Sanges, Antonello Mallamaci

**Affiliations:** Laboratory of Cerebral Cortex Development, SISSA, Trieste, Italy; Laboratory of Computational Genomics, SISSA, Trieste, Italy

**Author notes:** Corresponding Author: Antonello Mallamaci Lab of Cerebral Cortex Development SISSA via Bonomea 265 - 34136 Trieste - Italy.

**Keywords:** Foxg1, translation, Grin1, NMDAR, neuronal activity

## Abstract

Mainly known as a transcription factor patterning the rostral brain and governing its histogenesis, Foxg1 has been also detected outside the nucleus, however biological meaning of that has been only partially clarified. Here, moving from Foxg1 expression in cytoplasm of neocortical neurons, we investigated its implication in translational control. We documented an impact of Foxg1 on ribosomal recruitment of *Grin1*-mRNA, encoding for the main subunit of NMDA receptor. Next, we showed that Foxg1 increases Grin1 protein level by enhancing translation of its mRNA, while not increasing its stability. Such enhancement was associated to augmented translational initiation and, possibly, polypeptide elongation. Molecular mechanisms at the basis of this activity included Foxg1 interaction with Eif4e and Eef1d as well as with *Grin1*-mRNA. Besides, we found that, within murine neocortical cultures, Grin1 *de novo* synthesis undergoes a prominent and reversible, homeostatic regulation and Foxg1 is instrumental to that. Finally, through TRAP-seq, we discovered that Foxg1 is implicated in the translation of hundreds of neuronal genes at the level of ribosome engagement and progression. All that points to Foxg1 as a key effector, crucial to multi-scale temporal tuning of neocortical pyramid activity, an issue with profound physiological and neuropathological implications.

## INTRODUCTION

Foxg1 is an ancient transcription factor mastering a number of developmental processes that take place in the rostral brain. These include early activation of pan-telencephalic (Hanashima et al., 2007), subpallial (Manuel et al., 2010), and paleo-neo-pallial (Muzio and Mallamaci, 2005) programs, promotion of neural precursors selfrenewal (Martynoga et al., 2005), balance between neuronogenesis and gliogenesis (Brancaccio et al., 2010; Falcone et al., 2019; Frisari et al., 2022; Santo et al., 2022), and laminar specification of neocortical neurons (Hanashima et al., 2004; Miyoshi and Fishell, 2012; Toma et al., 2014). Later, it promotes morphological maturation of glutamatergic (Brancaccio et al., 2010; Chiola et al., 2019; Yu et al., 2019) and gabaergic (Zhu et al., 2019) telencephalic neurons. Moreover, Foxg1 sustains activity and excitability of these neurons (Frisari et al., 2022; Tigani et al., 2020; Zhu et al., 2019), via an articulated impact on transcription of specific gene-sets (Artimagnella and Mallamaci, 2019; Tigani et al., 2020). Besides, it is in turn transiently promoted by neuronal activity (Fimiani et al., 2016; Tigani et al., 2020). Finally, Foxg1 promotes hippocampal plasticity, by enhancing NMDA receptor-mediated currents (Yu et al., 2019). As a result of such a pleiotropic impact on brain development and neuronal function, *Foxg1* mutations result in complex, cognitive and behaviourial, phenotypes, in both mutant mouse models and human patients. In the mouse, loss of *Foxg1* leads to defective social interaction and impaired spatial learning and memory (Shen et al., 2006; Yu et al., 2019). Moreover, a co-misregulation of *Foxg1* in postnatal, excitatory as well as inhibitory neurons, is necessary and sufficient to evoke the emergence of ASD-like phenotypes (Miyoshi et al., 2021). In humans, >120 distinct *FOXG1* mutations result into a complex series of neuropathologies, collectively referred to as *FOXG1* syndrome, including brain dismorphologies, epilepsy and ASD-like symptoms (Wong et al., 2019). Moreover, a specific *FOXG1* upregulation has been detected in brain organoids originating from ASD-patient iPSCs (Mariani et al., 2015).

Albeit mainly known as a transcription factor (Kumamoto and Hanashima, 2017), Foxg1 was also previously reported to be in the *cytoplasm* of olfactory placode and early born neocortical neurons (Regad et al., 2007), as well as in the cytoplasm and mitochondria of a hippocampal neuronal line and whole brain homogenates (Pancrazi et al., 2015). Next, three high throughoutput screenings, in HEK293T, yeast and N2A cells (Li et al., 2015; Stelzl et al., 2005; Weise et al., 2019) showed that FOXG1 may interact with a number of factors implicated in post-transcriptional gene regulation, including translation. In addition, we noticed that Foxg1 harbors a YATHHLT motif (at 366-372 position), conserved among vertebrates (not shown) and reminiscent of the eIF4E-binding motif detectable in eIF4-BP, eIF4G and other effectors (Nédélec et al., 2004). These observations suggest a possible involvement of Foxg1 in control of mRNA translation.

To address this issue, we interrogated primary neocortical cultures via a variety of complementary experimental approaches. We found that - in neuronal soma as well as in neurites - Foxg1 promotes translation of *Grin1*, encoding for the main subunit of the NMDA receptor and playing a pivotal role in neuronal plasticity, and this requires Foxg1 interaction with Eif4e. We also demonstrated that Foxg1 promotes fast homeostatic tuning of Grin1, an issue of potential relevance to etiopathogenesis of *FOXG1*-linked neurological disorders. Finally, we got evidence that Foxg1 modulates ribosomal recruitment of dozens of other mRNAs encoding for effectors of neuronal activity, and it also affects ribosome progression. In this way, beyond its “slow” and cell-wide transcriptional impact on gene expression, Foxg1 might exert a far more spatio-temporally articulated control on neuronal functions.

## RESULTS

### Foxg1 promotes *Grin1*-mRNA translation in neocortical neurons

To corroborate previous reports of non-nuclear Foxg1 localization, we scored neurons from DIV8 cultures of murine E16.5 neocortical precursors (**Figure 1A-F**) and DIV7 cultures of rat P2 hippocampal precursors (**Figure 1G,H**) for distribution of extra-nuclear αFoxg1 immunoreactivity. As expected, in the former case, Foxg1 was specifically detectable in neuronal soma, as well as in punctate-Psd95^+^ dendrites (**Figure 1C,D**) and Smi312^+^ axons (**Figure 1E,F**). In the latter, although mainly associated to mitochondria, non-nuclear Foxg1 signal was not restricted to them (**Figure 1H**).

**Figure 1.**
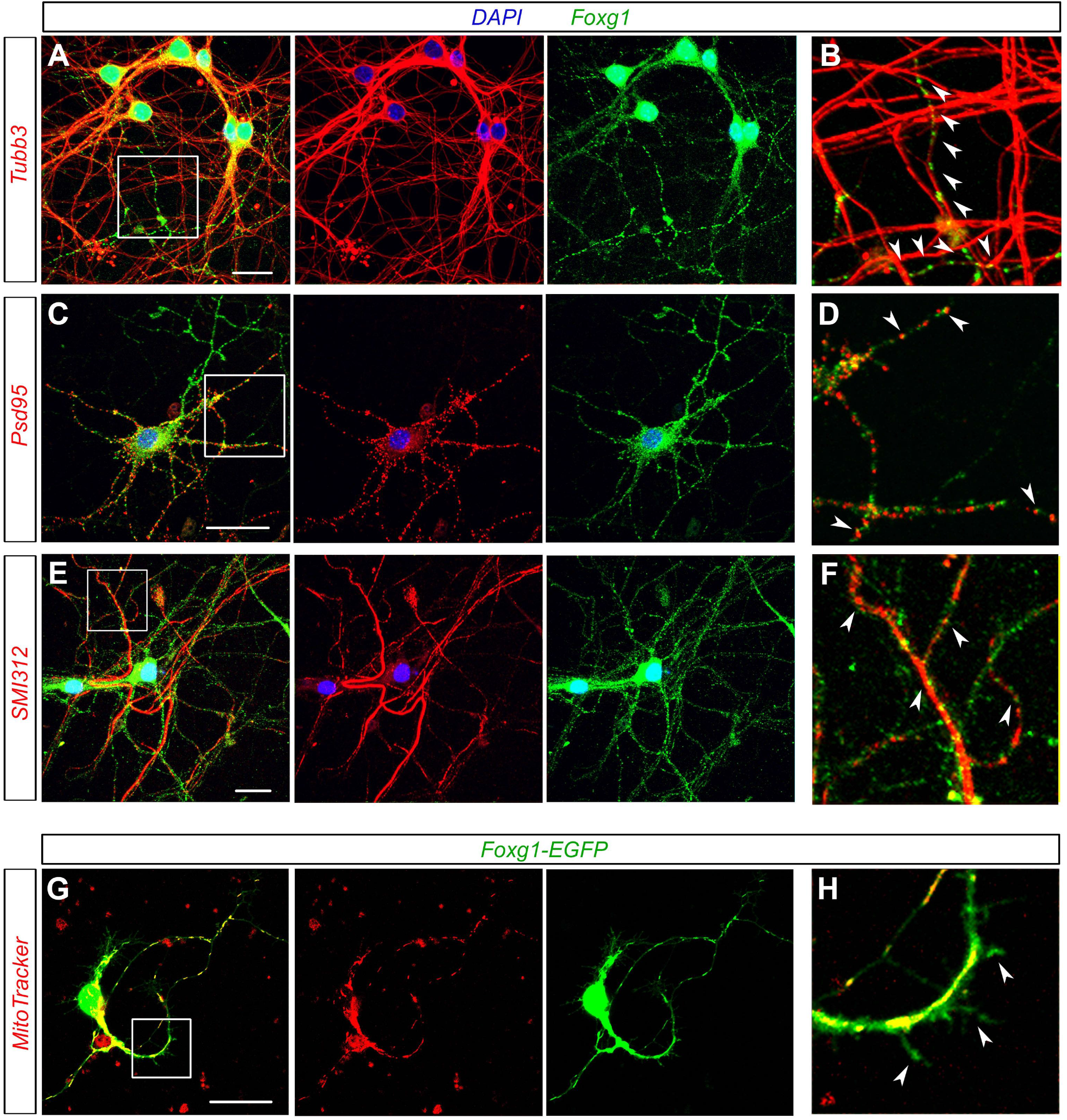
Neuronal, subcellular Foxg1 localization. **(A-F)** Preparations obtained by dissociation of E16.5 murine neocortices were cultured up to day in vitro 8 (DIV8), under a pro-differentiative medium supplemented with AraC. They were co-immunoprofiled for Foxg1 and, alternatively, Tubb3 (A, B), Psd95 (C,D), and SMI312 antigen (E,F), by confocal microscopy. Arrowheads point to Foxg1 immunoreactive grains adjacent to Tubb3-positive bundles (B), Psd95-positive spots (D), and SMI312-positive bundles (F). **(G, H)** Preparations obtained by dissociation of P2 rat hippocampi were cultured up to DIV7, under a pro-differentiative medium. Such cultures were transduced at DIV2 by lentiviral vectors driving p(Pgk1)/TetON-controlled Foxg1-EGFP chimera expression, preterminally labelled by the red MitoTracker dye for 30 min, and finally profiled by live-confocal microscopy. Arrowheads in (H) point to cytoplasmic Foxg1-EGFP signal which does not co-localize with mitochondria. High power magnifications of (A,C,E,G) panel insets are in (B,D,F,H), respectively. Scalebars, 20 µm.

Next, to preliminary explore Foxg1 implication in translation, we selected a small sample of genes undergoing translational regulation and/or being implicated in fine tuning of neuronal activity (*Grid1*, *Grin1, Slc17a6*, *Gria1*, *Gabra1*, *Bdnf* - *2c* and *4* isoforms -, *Psd95*, and *Foxg1)* (Baez et al., 2018; Dörrbaum et al., 2020), and we evaluated the impact of *Foxg1* expression level on ribosomal engagement of their mRNAs. For this purpose, we used neocortical neurons obtained from E16.5 *Rpl10a^EGFP-Rpl10a/+^* mouse embryos (Zhou et al., 2013), engineered to conditionally overexpress *Foxg1* (*Foxg1*-OE) or a *PLAP* control (**Figure S1**). Four days after transgenes activation, at DIV8, we analyzed them by Translating Ribosome Affinity Purification (TRAP)-qRTPCR **(Figure 2A, left)**. Specifically, by means of an anti-EGFP antibody, we purified RNA associated to EGFP-tagged-ribosomes (IP component) and supernatant cellular RNA (SN component). Then, we profiled these RNAs by qRT-PCR, paying attention to the above-mentioned candidate genes. Upon normalization against *Rpl10a*-mRNA, IP/SN ratios were averaged and further normalized against *Plap* controls. *Grid1* and *Grin1* IP/SN ratios were upregulated in *Foxg1*-OE samples (+50.7±16.2%, with p<0.01 and n=6,6; and +34.7±7.0%, with p<0.01 and n=6,6, respectively), suggesting that Foxg1 may promote ribosomal engagement of their mRNAs. The same index was not affected in the remaining cases **(Figure 2B)**.

**Figure 2.**
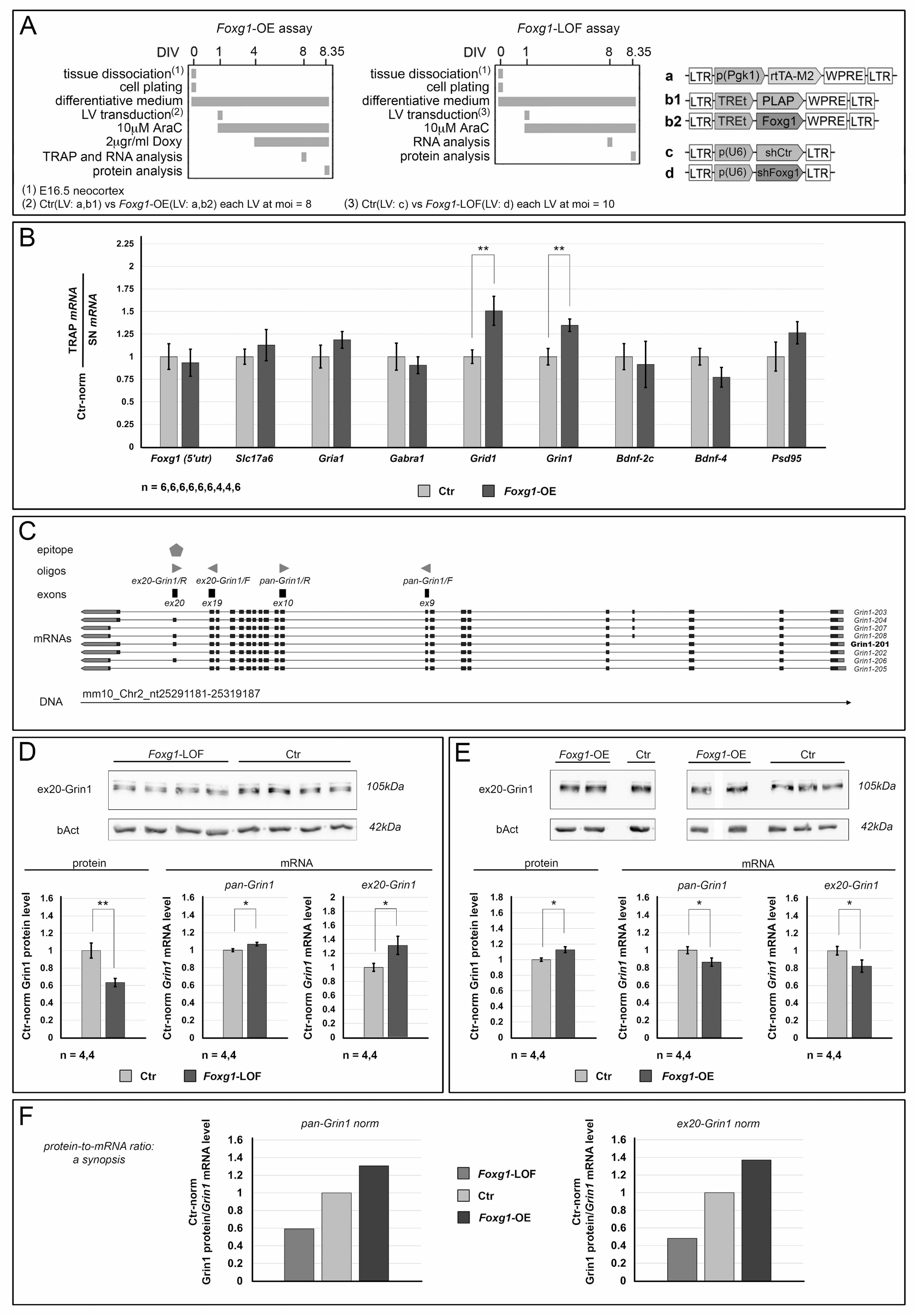
Impact of *Foxg1* manipulation on ribosomal allocation and protein output of selected neuronal transcripts. **(A)** Protocols and lentiviral vectors used to engineer neocortical cultures to conditionally overexpress *Foxg1* (Foxg1-OE, left panel) or reduce its level (*Foxg1*-LOF, right panel). **(B)** Comparative, Translating Ribosome Affinity Purification (TRAP) quantification of ribosome-associated mRNA fraction (TRAP-mRNA) and its supernatant fraction (SN-mRNA), referring to selected neuronal transcripts, in *Foxg1*-OE cultures. mRNA levels measured by qRT-PCR, and double normalized, against *Rpl10a*-mRNA and controls. **(C)** *Grin1* gene locus with the main polypeptide-encoding transcripts originating from it. The top polygon represents the protein epitope recognized by the anti-Grin1 antibody used in Western blot assays. Arrowheads indicate oligos used to quantify *Grin1*-mRNA, distinguishing between ex20-containing Grin1 (ex20-*Grin1*) isoforms and pan-*Grin1* isoforms. **(D,E)** Western blot analysis of Grin1 protein and qRT-PCR quantification of pan-*Grin1* and ex20-*Grin1* mRNA isoforms, upon *Foxg1*-LOF (D) and *Foxg1*-OE (E) manipulations. Protein levels double normalized against bAct and controls, mRNA levels against *Rpl10a*-mRNA and controls. **(F)** Progression of “normalized Grin1-protein” - to - “normalized *Grin1*-mRNA” ratio upon *Foxg1* manipulation, referring to pan-*Grin1* mRNA (left graph) or ex20-*Grin1*-mRNA (right graph). Throughout figure, *n* is the number of biological replicates, i.e. independently cultured and engineered preparations, originating from a common neural cell pool. Statistical evaluation of results was performed by one-way ANOVA, one-tailed and unpaired. * *p*<0.05, ** *p*<0.01. Errors bars indicate s.e.m.

To corroborate this finding and explore its biological meaning, we focused our attention on *Grin1* gene, encoding for the main subunit of NMDA receptor, whose activity is impaired upon conditional *Foxg1* ablation in the murine hippocampus (Yu et al., 2019). For this gene, we evaluated the protein-to-mRNA ratio upon artificial modulation of *Foxg1* expression. Tests were run in cultures of E16.5+DIV8 murine neocortical neurons, engineered to conditionally overexpress *Foxg1* **(Figure S1** and **2A,left**) or reduce its level (*Foxg1*-LOF) **(Figure S1** and **2A, right)**. Grin1 protein was quantified via WB, by a monoclonal antibody recognizing an epitope encoded by *Grin1*-exon 20. *Grin1*-mRNA was measured via qRTPCR, by two oligonucleotide pairs, detecting all *Grin1* isoforms (*pan-Grin1*) or exon20-containing ones (*ex20-Grin1*) **(Figure 2C)**. Normalized against bAct, Grin1 protein was decreased (−36.7±4.7%, with p<0.005 and n=4,4) and increased (+12.8±3.8%, with p<0.02 and n=4,4), following down-**(Figure 2D)** and up-regulation **(Figure 2E)** of *Foxg1*, respectively. Opposite trends were displayed by *pan-Grin1*-mRNA (+6.9±2.0, with p<0.02 and n=4,4, in *Foxg1*-LOF samples; −13.7±4.7%, with p<0.04 and n=4,4, in *Foxg1*-OE ones). Remarkably, such opposite trends were even more pronounced in case of *ex20-Grin1*-mRNA (+31.5±13.0%, with p<0.04 and n=4,4, in *Foxg1*-LOF samples; −17.7±7.0%, with p<0.05 and n=4,4 in *Foxg1*-OE ones) **(Figure 2D,E)**. Finally, to get a comprehensive index of post-transcriptional *Foxg1* impact on Grin1 level, we calculated the “Grin1-protein/*Grin1*-mRNA” ratios peculiar to *Foxg1*-misexpressing cultures and normalized them against their controls. Such ratios ranged from 0.59 (*Foxg1*-LOF) to 1.31 (*Foxg1*-OE), referring to *pan-Grin1*-mRNA, from 0.48 (*Foxg1*-LOF) to 1.37 (*Foxg1*-OE), taking specifically into account *ex20-Grin1*-mRNA **(Figure 2F)**. All that suggests that *Foxg1* exerts a robust positive impact on post-transcriptional Grin1-protein tuning.

Next question was: (1) does *Foxg1* enhance translation of *Grin1*-mRNA and/or (2) does it diminish degradation of Grin1 protein ?

As for (1), we assessed Grin1 translation rates in E16.5+DIV8 neocortical cultures made *Foxg1*-LOF by RNAi (**Figure S1**). To this aim, we terminally pulsed these cultures with puromycin and we measured levels of nascent Grin1 protein, (n)Grin1, via anti-Grin1/anti-puromycin-driven proximity ligation assay (PLA) **(Figure 3A)**. To distinguish among translation of all *Grin1* isoforms (*pan-Grin1*) and exon20-containing ones (*ex20-Grin1*), two anti-Grin1 antibodies were alternatively used in addition to anti-puromycin (**Figure 3A**, a and b). The former, anti-Grin1-NH2-term, recognizes the aminoterminal protein region shared by all isoforms, and the latter, anti-Grin1-COOH-term, a more carboxyterminal ex20-encoded epitope (**Figure 2C**). Next, the analysis was run on whole neurons or it was limited to neurites. In case of whole neurons, two indices of (n)Grin1 levels were evaluated, the cumulative PLA signal per cell and the cumulative PLA signal per spot. In case of neurites, the first parameter was hard to evaluate, and measure was restricted to cumulative PLA signal per spot.

**Figure 3.**
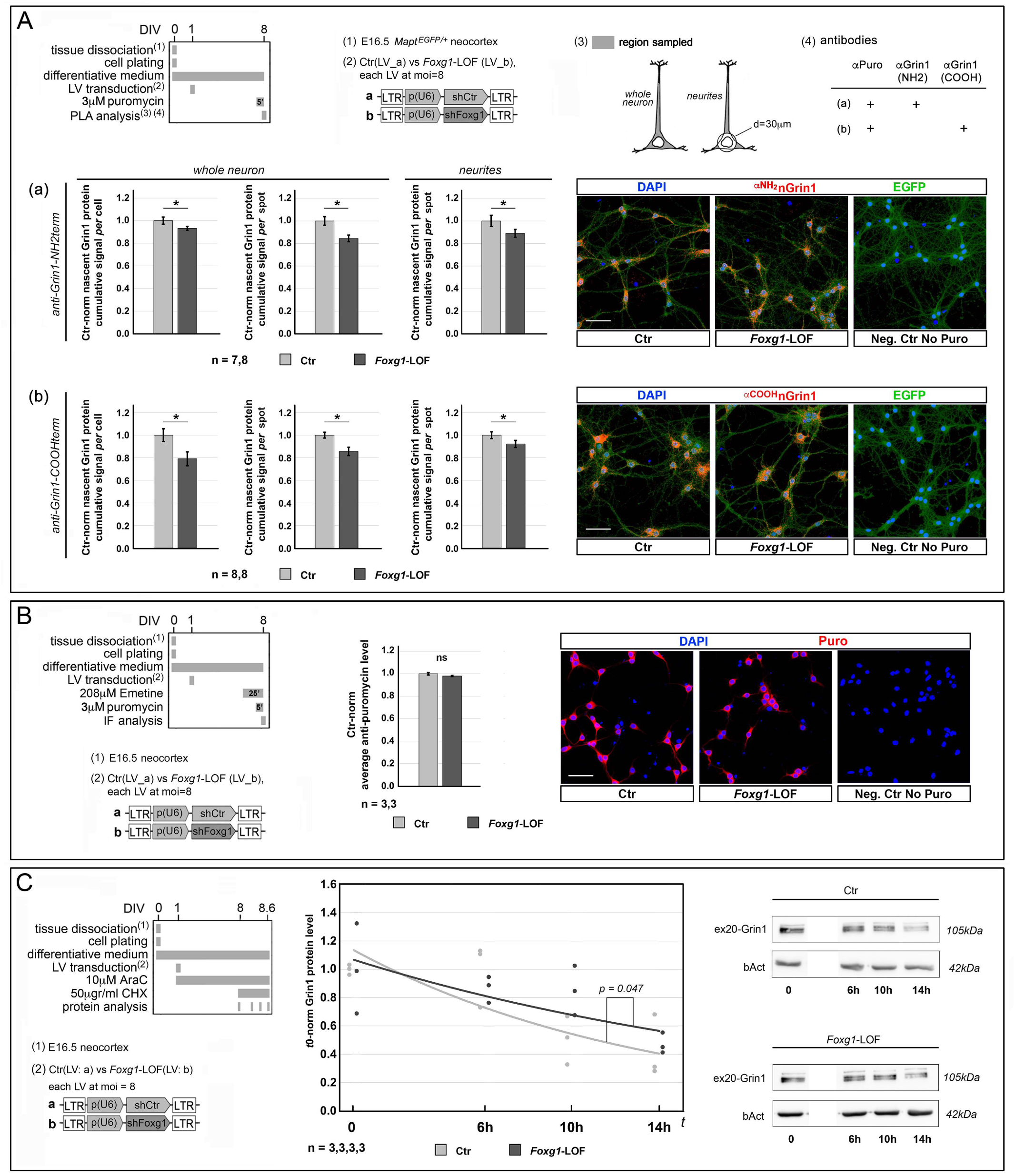
Quantification of nascent Grin1 protein (A) and nascent proteome (B), and evaluation of Grin1-protein degradation rate (C) in *Foxg1*-LOF neurons. (A) To top, protocols (including lentiviruses employed, and operational details of proximity ligation assay (PLA) analysis), to bottom, results. Graphs represent quantitative confocal immunofluorescence (qIF) assesment of nascent Grin1 protein, (n)Grin1, performed upon *Foxg1* down-regulation, terminal (5’) puromycin administration and subsequent anti-Grin1/anti-puromycin-driven PLA. Two anti-Grin1 antibodies were alternatively used, recognizing (a) aminoterminal (anti-Grin1-NH2-term) and (b) carboxyterminal (anti-Grin1-COOH-term) protein regions. Neuron cell silhouettes were identified by direct EGFP fluorescence, driven by the *Mapt^EGFP^*transgene. PLA signal was quantified throughout the whole neuron or restricted to neurites. As indices of (n)Grin1 levels, shown are the average cumulative qIF signals per cell and the average cumulative qIF signals per spot. **(B)** To left, protocols (including lentiviruses employed), to right, results. Graph represents quantitative confocal immunofluorescence (qIF) evaluation of nascent total puromycinilated proteins, performed upon *Foxg1* down-regulation, terminal emetine (25’) and puromycin (5’) supplementation, and final anti-puromycin-driven qIF. In both (A) and (B), results were normalized against controls, error bars indicate s.e.m., and statistical evaluation of results was performed by one-way ANOVA, one-tailed and unpaired (* p<0.05). In both (A) and (B), included are examples of primary data referred to by the corresponding graphs. Scalebars, 50μm. **(C)** To left, protocols and lentiviruses employed for this analysis, to right, results. Graph represents progression of Grin1-protein levels at different t_i_ times, evaluated by western blot, upon *Foxg1* down-regulation and subsequent 50μg/ml cycloheximide (CHX) blockade of translation. For each genotype, results double normalized, against (t_i_)bAct protein levels and (t_0_)average values. Superimposed, exponential trendlines. Statistical evaluation of results performed by ANCOVA test. Included are examples of primary data referred to by the corresponding graphs. Throughout figure, *n* is the number of biological replicates, i.e. independently cultured and engineered preparations, originating from a common neural cell pool.

Compared to controls, whole neuron (n)pan-Grin1 signal was reduced in *Foxg1*-LOF samples, by 6.7±1.8% per cell (*p<*0.039, *n*=7,8), and 15.6±2.9% per spot (*p<*0.003, *n*=7,8). In a similar way, neurite (n)pan-Grin1 signal per spot was decreased by 11.0±3.5% (*p<*0.043, *n*=7,8) **(Figure 3A**,a). As for (n)ex20-Grin1, its signal was also reduced in *Foxg1*-LOF samples, by 20.9±6.1% per cell (*p<*0.013, *n*=8,8), and 14.4±3.6% per spot (*p<*0.003, *n*=8,8). In a similar way, neurite (n)ex20-Grin1 signal per spot was also decreased, by 7.7±3.2% (*p<*0.048, *n*=8,8) **(Figure 3A**,b). In a few words, dampening Foxg1 reduces Grin1 translation, in soma as well as in neurites. Interestingly, this is peculiar to Grin1, and it does not apply to all translatome, as shown by anti-puro immunofluorescence (IF) run on *Foxg1*-LOF neural cultures terminally treated by emetine and puromycin **(Figure 3B)**. Taking into account the 6.9% and 31.5% increases undergone by *pan-Grin1*- and *ex20*-*Grin1*-mRNA, respectively, upon *Foxg1* down-regulation (see **Figure 2D**), these data suggest that *Foxg1* specifically promotes *Grin1*-mRNA translation, with particular emphasis on its ex20 isoform.

As for (2), we evaluated Grin1 degradation rates in similar *Foxg1*-LOF neocortical samples. To this aim, we blocked translation by cycloheximide and monitored time course progression of previously synthesized Grin1 protein over 14 hours **(Figure 3C,left)**. Remarkably, Grin1 degradation rate resulted to be not increased, but - rather - slightly decreased upon *Foxg1* downregulation. Specifically, the Grin1(t_i_)/Grin1(t_0_) ratio equalled e^-0.046*hours^ and e^-0.074*hours^ in *Foxg1*-LOF cultures and controls, respectively (with *p*<0.048, *n*=3,3,3,3) **(Figure 3C,right)**. This result rules out that the *Foxg1*-dependent increase of “Grin1-protein/*Grin1*-mRNA ratio” **(Figure 2F)** may be due to *Foxg1* inhibition of Grin1 protein degradation.

### Foxg1 promotes ex20-Grin1 translation by enhancing ribosome recruitment at *Grin1*-mRNA and accelerating Grin1 polypeptide synthesis

Cumulative translation gain is a function of the rate of ribosomes engagement to mRNA and the speed at which they progress along mRNA-cds. We have shown that Foxg1 promotion of Grin1 translation firstly reflects an improved recruitment of ribosomes to *Grin1*-mRNA (**Figure 2B**). We wondered: is *Foxg1* also able to promote ribosome progression along *Grin1*-cds?

To address this issue, we set a dedicated puro-PLA run-off assay (**Figure S2A-C**), implementing it in neocortical neuronal cultures. Specifically, upon blockade of de novo ribosome recruitment to mRNA cap by harringtonine, ribosomes were allowed to continue ongoing translations for a time presumptively close to that required for full *Grin1*-mRNA translation. At the end of this time, unfinished Grin1 polypeptides were labeled by terminal puromycin supplementation and revealed by anti-Grin1/puro-PLA. The PLA signal was subtracted from its t=0 counterpart (evaluated prior to harringtonine treatment) and the resulting difference, normalized against t=0 value, was employed as an index positively correlated to ribosome progression speed along *Grin1*-cds (**Figure S2; Figure 4A**, top). Such run-off assay was performed on *Foxg1*-LOF samples and their “wild type” controls, driving PLA by anti-Grin1-NH2-term (which recognizes *all* Grin1 polypeptides). As expected, 11 minutes after harringtonine supplementation, (n)pan-Grin1 signal underwent a substantial decline compared to its t=0 value (almost −40%), however no difference was detected between *Foxg1*-LOF samples and controls (**Figure 4A**, bottom). This suggests that no generalized change of Grin1 translation speed occurs upon Foxg1 manipulation.

**Figure 4.**
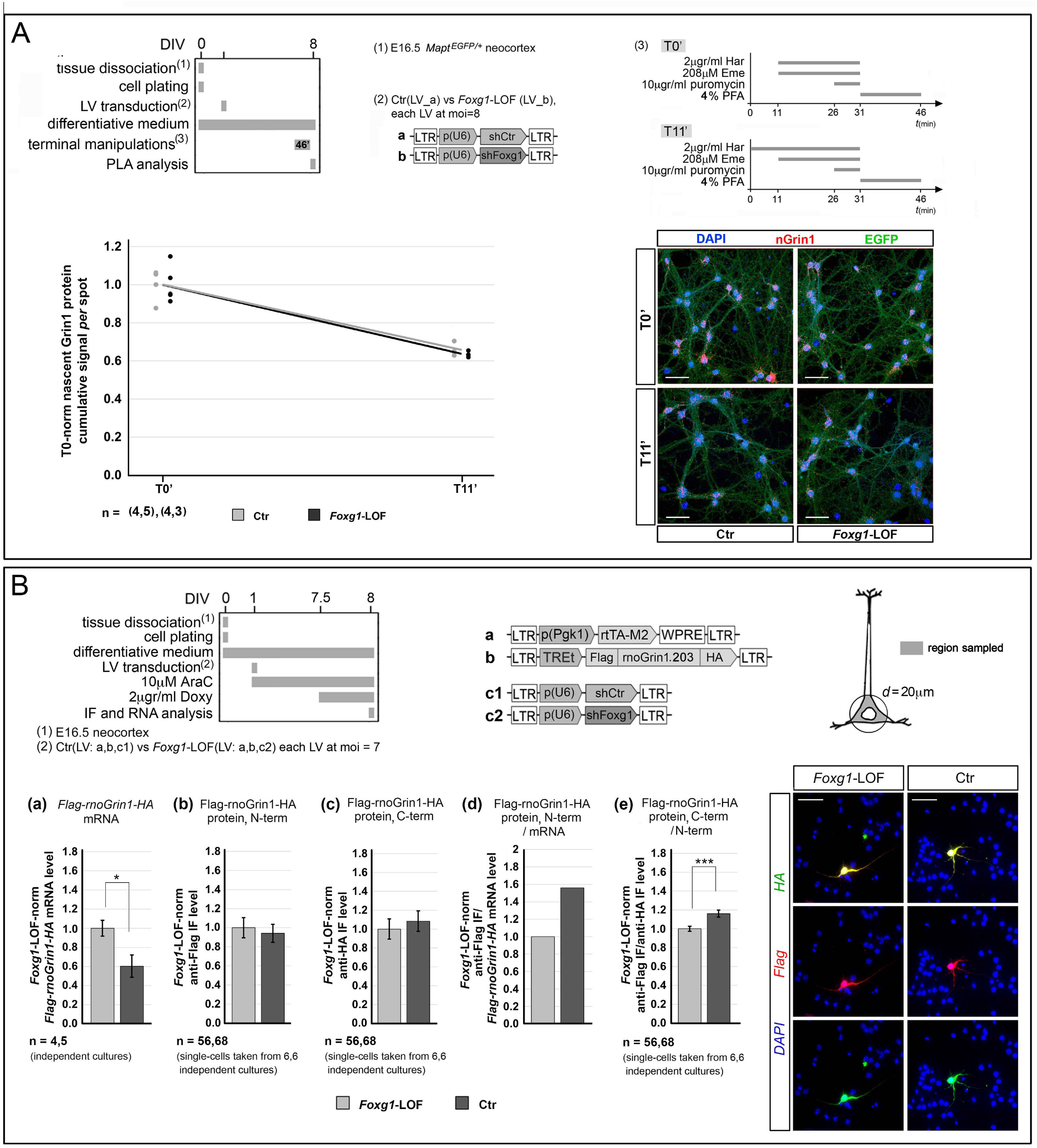
Evaluation of endogenous pan-*Grin1* elongation rate by run-off assay (A) and assesment of ex20-*Grin1* translational initiation and elongation rates by means of a heterologous *Flag-rnoGrin1.203-HA* sensor in *Foxg1*-LOF neurons (B). (A) To top, protocols (including lentiviruses employed, and operational details of the translational run-off assay), to bottom, results. Graph represents progression of nascent Grin1 levels evaluated by anti-Grin1-NH2-term/anti-puromycin-driven PLA, upon *Foxg1* down-regulation, in basal conditions (T0’) and 11 minutes after 2μg/ml harringtonine (har) blockade of translation initiation (T11’). In both cases, ribosome progression was subsequently inhibited by 208μM emetine (eme), and nascent polypeptides were terminally labeled by 10μg/ml puromycin (puro). For each genotype, results normalized, against (T0’) average values. Superimposed, linear trendlines. Statistical evaluation of results performed by ANOVA test (one-tail, unpaired). *n* is the number of biological replicates, i.e. independently cultured and engineered preparations, originating from a common neural cell pool. Included are examples of primary PLA data referred to by the corresponding graphs. Scalebars, 50μm. **(B)** Protocols (*top-left*) and lentiviral vectors (*top-mid*) used to make neocortical cultures *Foxg1*-LOF and profile mRNA and protein outputs of their doxycycline-activatable, *Flag-rnoGrin1.203-HA* transgene. *rnoGrin1.203* is the ortholog of *mmuGrin1.*201 isoform, including exon 20. The silhouette (*top*-*right*) represents an idealized pyramidal cell, where the somatic cytoplasm sampled is highlighted in grey. Graphs (*bottom-left*) showing: (a) qRT-PCR quantification of *Flag-rnoGrin1.203-HA*-mRNA levels in bulk cultures, primarily normalized against *Rpl10a*-mRNA; (b,c) single-cell IF quantification of average N-term-Flag and C-term-HA signals detectable in neuronal somata; (d) “N-term-Flag to *Flag-rnoGrin1.203-HA*-mRNA” ratio, as an index of rnoGrin1.203 translational initiation; (e) “C-term-HA to N-term-Flag” ratio, as an index of rnoGrin1.203 translational elongation. In all five graphs, results normalized against *Foxg1*-LOF samples. *n* is the number of biological replicates. These are: (a) independently cultured and engineered preparations, originating from a common neural cell pool; (b,c,e) single-cells, evenly taken from multiple, independent *Foxg1*-LOF and control cultures. Statistical evaluation of results was performed by one-way ANOVA, one-tailed and unpaired. * *p*<0.05, *** *p*<0.001. Errors bars indicate s.e.m. Included are (*bottom-right*) examples of anti-Flag and anti-HA immunostainings in *Foxg1*-LOF and control cultures. Scalebars, 50μm.

Next, we wondered: what about the ex20-Grin1 isoform? Unfortunately we could not address this point by a PLA-run-off assay, coupling anti-Grin1-COOH-term with anti-puromycin, because of the close proximity between the corresponding epitope and the protein terminus, and the consequent difficulty in setting up a suitable harringtonine treatment time. To circumvent this conundrum, we built a dedicated translation sensor based on *rnoGrin1.203,* namely the rat ortholog of the exon 20-including, *mmuGrin1.*201 isoform. Specifically, this was a TetON-controlled transgene, encoding for a double-tagged, ^N-term^Flag-rnoGrin1.203-^C-term^HA chimera. We delivered it to *Foxg1*-LOF and control E16.5 neocortical murine cultures, and we acutely activated it by doxycycline. 12 hours later, we evaluated levels of Flag and HA epitopes by quantitative immunofluorescence, and we calculated the “HA epitope/Flag epitope” ratio, as an index of polypeptide elongation **(Figure 4B)**. Compared to *Foxg1*-LOF samples, such ratio was increased by +16.0±3.7% in controls (*p*<0.001 and *n*=55,68) (**Figure 4B,e graph**), suggesting that *Foxg1* promotes elongation of the *mmuGrin1.*201-encoded polypeptide.

Moreover, we took advantage of this sensor to further assay *Foxg1* impact on ribosome *engagement* to *ex20*-*Grin1*-mRNA. We noticed that, normalized against *Foxg1*-LOF samples, controls displayed a pronounced downregulation of *Flag-rnoGrin1.203-HA*-mRNA (−39.6±11.7%, with *p*<0.018 and *n*=4,5), and poorly affected Flag and HA epitope levels **(Figure 4B**,**a-c graphs)**. In this way the “Flag epitope/*Flag-rnoGrin1.203-HA*-mRNA” ratio, an index of translation initiation, was upregulated by +56.0% in controls compared to *Foxg1*-LOF samples **(Figure 4B**,**d graph)**, corroborating the positive *Foxg1* impact on initiation of *mmuGrin1.*201 translation.

### Foxg1 physically interacts with selected translation factors

We have shown that Foxg1 enhances translation of Grin1. Next question was: does Foxg1 act in this context (1) as a canonical nuclear transcription factor, tuning expression of translation factor genes, or (2) straightly as a cytoplasmic “translation modulator”? Previous Foxg1 interaction screening reports (Li et al., 2015; Stelzl et al., 2005; Weise et al., 2019) as well as limited responsivity of translation factors’ mRNA levels to *Foxg1* overexpression (**Table S1**) suggested us that type (2) mechanisms might be prevailing.

To preliminarily cast light on this issue, we took advantage of HEK293T preparations, expressing Foxg1 and distinct translation-related, putative interactors of it, tagged by V5 or Flag. We specifically focused on EIF4E, EEF1D, EEF1G, PUM1, evaluating their interaction with Foxg1 by quantitative proximity ligation assay (qPLA), driven by anti-Foxg1 and anti-tag antibodies (**Figure 5A,left**). Compared with their technical controls (“αFoxg1 only” and “αtag only”), EEF1D and EIF4E gave robust PLA signals. Normalized against averages of such controls, these equalled 3.4±0.6 (with *p*_vs-αFoxg1-only_<0.04, *p*_vs-αtag-only_<0.05 and *n*=2,2,2), and 3.7±0.8 (with *p*_vs-αFoxg1-only_<0.04, *p*_vs-αtag-only_<0.05 and *n*=2,2,2), respectively, suggesting that EEF1D and EIF4E interaction with Foxg1 is genuine. Conversely, EEF1G and PUM1 signals hardly overcame the corresponding controls (**Figure 5A,right**).

**Figure 5.**
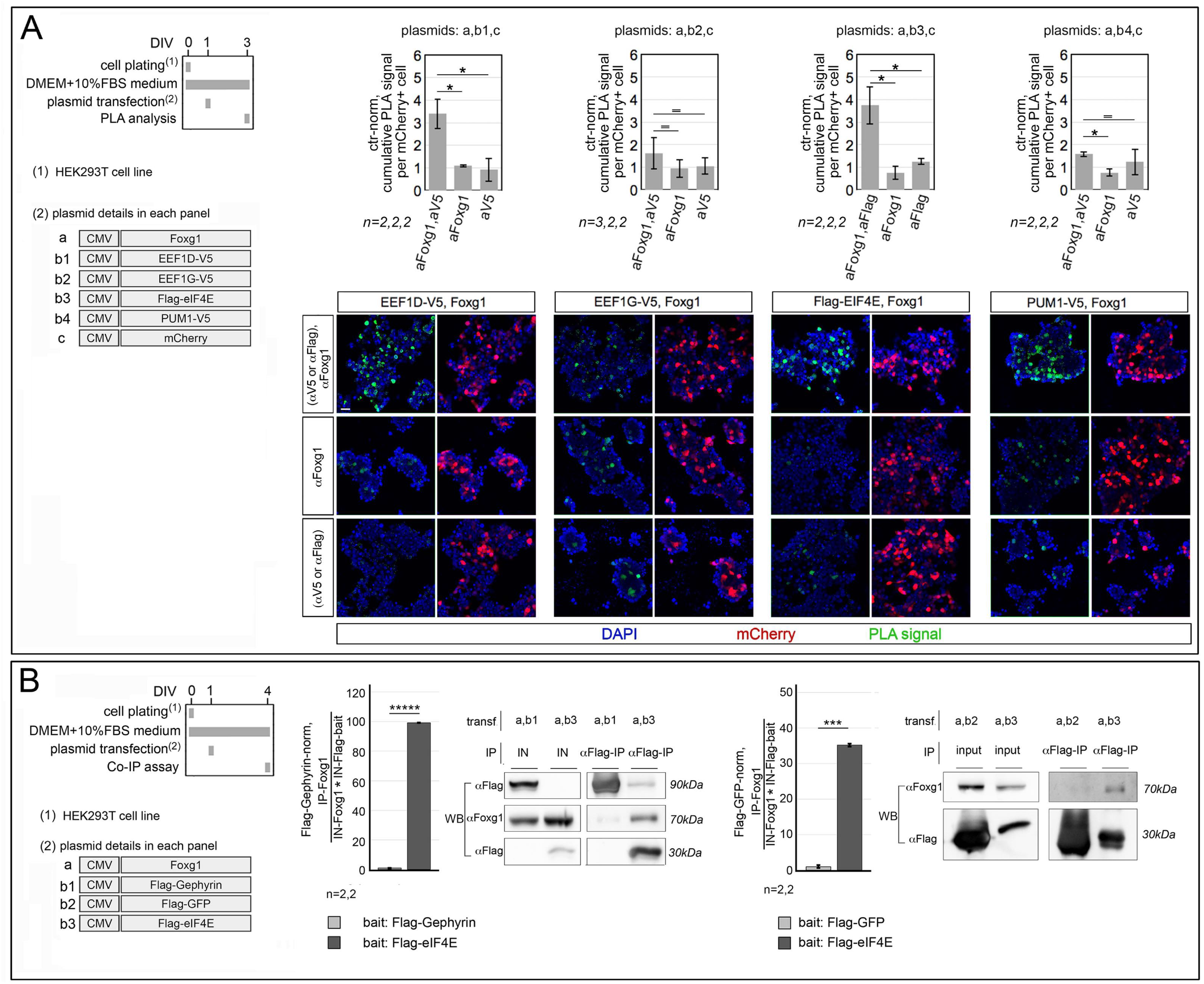
Assessment of Foxg1-protein interaction with selected translation factors in engineered HEK293T cells. **(A)** Interaction evaluation based on proximity ligation assay (PLA). To left, protocols and lentiviral vectors used, to right, graphical representation of results (upper row) and examples of primary data (lower row). Cells co-transfected, by Foxg1- and Flag-TranslationFactor-or TranslationFactor-V5-encoding transgenes, and further transfected by an mCherry transgene, as an internal control. Interaction signals revealed by anti-Foxg1/anti-Flag-or anti-Foxg1/anti-V5-driven PLAs, mCherry revealed by direct fluorescence. For each co-transfection type, shown is the cumulative “green” PLA signal per transfected mCherry^+^ cell, normalized against the average values of the two corresponding negative controls (each obtained by omitting either primary antibody). **(B)** Evaluation of Foxg1/Eif4e interaction by immuno-precipitation-western-blot (IP-WB) analysis. To left, protocols and lentiviral vectors used, to right, graphical representation of results, with examples of primary data. Cells double transfected, by Foxg1- and Flag-Eif4e-encoding transgenes (negative controls: Flag-Gephyrin and Flag-EGFP), IP by anti-Flag, immuno-detection by anti-Foxg1. Densitometric IP-protein values double normalized, against Foxg1 and Flag-protein inputs, and further normalized, against negative controls. Throughout figure, *n* is the number of biological replicates, i.e. independently cultured and engineered preparations, originating from a common cell pool. Statistical evaluation of results performed by one-way ANOVA, one-tailed and unpaired. * *p*<0.05, *** *p*<0.001, ***** *p*<0.00001. Errors bars indicate s.e.m. Scalebars, 50μm.

To confirm Foxg1/EIF4E interaction (not included in previously published interaction reports), we further interrogated HEK293T cells cotransfected with Foxg1- and Flag-EIF4E-encoding transgenes, via quantitative-immunoprecipitation-western blot (qIP-WB) assays. Negative controls were set replacing *Flag-EIF4E* by *Flag-Gephyrin* and *Flag-GFP* transgenes (**Figure 5B,left**). Immunoprecipitated by anti-Flag and probed upon WB separation by anti-Foxg1, lysates of *Foxg1/Flag-EIF4E*-transduced cells specifically gave a robust Foxg1 enrichment. Normalized against the product of Foxg1 and Flag-protein inputs, and further normalized against *Flag-Gephyrin*- and *Flag-GFP*-transduced controls, IP-Foxg1 equalled 99.1±0.1 (with p<2.0*10^-6^ and *n*=2,2) and 35.2±0.4 (with p<2.0*10^-4^ and *n*=2,2), respectively (**Figure 5B,right**).

In a few words, within HEK293T cells overexpressing them, Foxg1 interacts with standard factors implicated in both translation initiation (EIF4E) and polypeptide elongation (EEF1D). Next, we wondered: does this also apply to the corresponding endogenous proteins of neocortical neurons? and, if so, which the relevance of that to Grin1 translation?

To address this issues, as a proof-of-principle, we scored primary neocortical cultures for the interaction between *endogenous* Foxg1 and *endogenous* Eif4e, by quantitative proximity ligation assay (qPLA) (**Figure 6A,left**). Compared with technical controls (“αFoxg1 only” and “αEif4e only”), the assay gave a robust signal. Normalized against controls’ average, the number of PLA-spots per cell equalled 3.6±0.6 (with *p*_vs-αFoxg1-only_<0.01, *p*_vs-αEif4e-only_<0.01 and *n*=4,4,4) (**Figure 6A**, graph a). Remarkably, a similar result was obtained, when restricting the analysis to neurites only (PLA signal=3.2±0.4, with *p*_vs-αFoxg1-only_<0.003, *p*_vs-αEif4e-only_<0.02 and *n*=4,4,4) (**Figure 6A**,**graph b**). All this suggests that *endogenous* Foxg1 and Ei4e genuinely interact within neocortical neurons, including their neurites. Next, to assess relevance of such interaction to Grin1 translation, we tried to dampen the former and evaluate consequences of that on the latter. Specifically, by means of a lentiviral vector, we transduced neuronal cultures with a transgene encoding for the mmu-Foxg1 aa357-381 polypeptide, harboring the putative, Eif4e-binding YATHHLT motif. Then, we monitored Foxg1/Eif4e interaction as well as nascent Grin1, (n)Grin1, levels. As expected, compared to a scrambled control, the Foxg1-Eif4e PLA signal was lowered, by −16.3%±3.8% (with p<0.05 and n=3,3) (**Figure 6B**,**graph a**), which resulted into a −29.9%±2.7% decrease of (n)Grin1 (p<0.02 and n=3,3) (**Figure 6B**,**graph b**). All that further suggests that Foxg1-Eif4e interaction contributes to mediate the positive impact that Foxg1 exerts on Grin1 translation.

**Figure 6.**
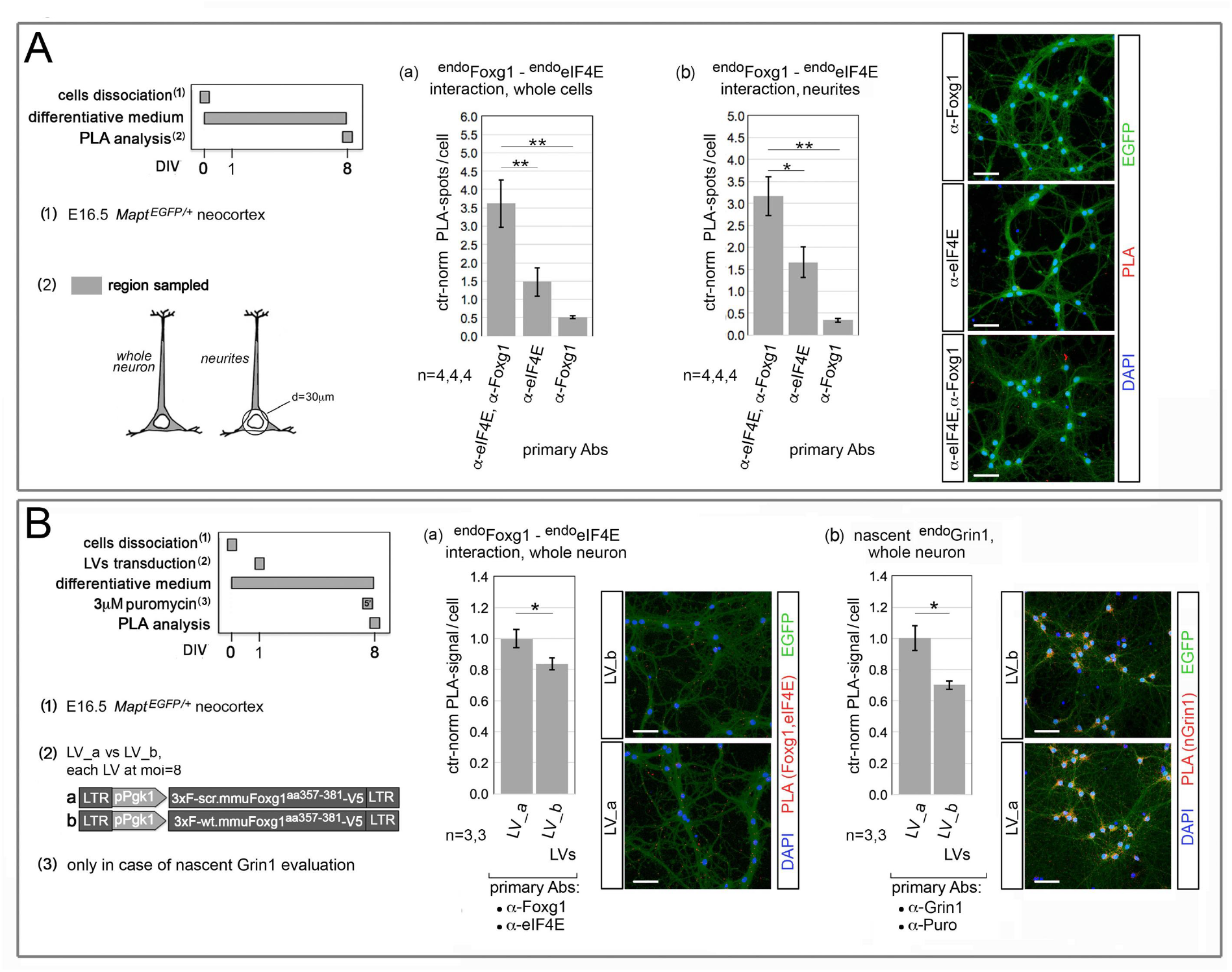
Assessment of Foxg1-Eif4e interaction (A) and its functional relevance to Grin1 translation (B) in neocortical neurons. **(A)** PLA assesment of endogenous-Foxg1/endogenous-Eif4E interaction (^endo^Foxg1-^endo^**Eif4e**), in whole neurons, or restricted to neurites. To left, protocols and lentiviral vectors used, to right, results. Assays run on cultures of *Mapt^EGFP^* neocortical neurons, interaction signals revealed by anti-Foxg1- and anti-Eif4e-driven PLA, performed on whole neurons (graph a) or restricted to neurites (graph b), according to graphically displayed criteria. Here, graphs report the numbers of spots/cell, normalized against the average of the two corresponding negative controls (each obtained by omitting either primary antibody). **(B)** PLA assesment of ^endo^Foxg1-^endo^eIF4E interaction (graph a) and nascent ^endo^Grin1 levels (graph b) in whole neurons, upon lentivirus-mediated over-expression of a tagged polypeptide including aa357-381 of murine Foxg1 protein (LV_b). A scrambled version of this polypeptide was used as a control (LV_a). To left, protocols and lentiviral vectors used, to right, results. Here, shown are cumulative PLA signals per cell, normalized against controls. Throughout figure, *n* is the number of biological replicates, i.e. independently cultured and engineered preparations, originating from a common cell pool. Statistical evaluation of results performed by one-way ANOVA, one-tailed and, paired or unpaired, in the case of (A) or (B), respectively. * *p*<0.05, ** *p*<0.01. Errors bars indicate s.e.m. Throughout figure, included are examples of primary data referred to by graphs. Scalebars, 50μm.

### Foxg1 physically interacts with *Grin1*-mRNA

To further support the hypothesis that Foxg1 promotes Grin1 synthesis as a translation factor, we investigated if Foxg1 interacts with *Grin1*-mRNA. To this aim, firstly, we quantified the fraction of endogenous *Grin1*-mRNA immunoprecipitated by an anti-Foxg1 antibody in lysates of E16.5+DIV8 neocortical neurons, by RNA immuno-precipitation (RIP)-qRTPCR (**Figure 7A**). This fraction exceeded the IgG background by 17.6±7.4 folds (with *p*<0.05, *n*=4,5) (**Figure 7A**, graph a); moreover, compared to “wild type” control, it showed a declining trend upon *Foxg1* knock-down (normalized against IgG, 10.3±1.5 vs 20.1±8.7, with *p*<0.15 and *n*=2,2) (**Figure 7A**, graph b). Then, we scored RNA extracted from neurons overexpressing a Foxg1-EGFP chimera and immunoprecipitated by an anti-EGFP antibody, for *Grin1*-mRNA enrichment. Remarkably, such enrichment equalled 6.1±0.8, upon normalization against PLAP expressing controls (with *p*<0.02, *n*=2,2) (**Figure 7A**, graph c). Altogether these results indicate that, within neocortical neurons, *endogenous* Foxg1 protein interacts with *endogenous Grin1*-mRNA.

**Figure 7.**
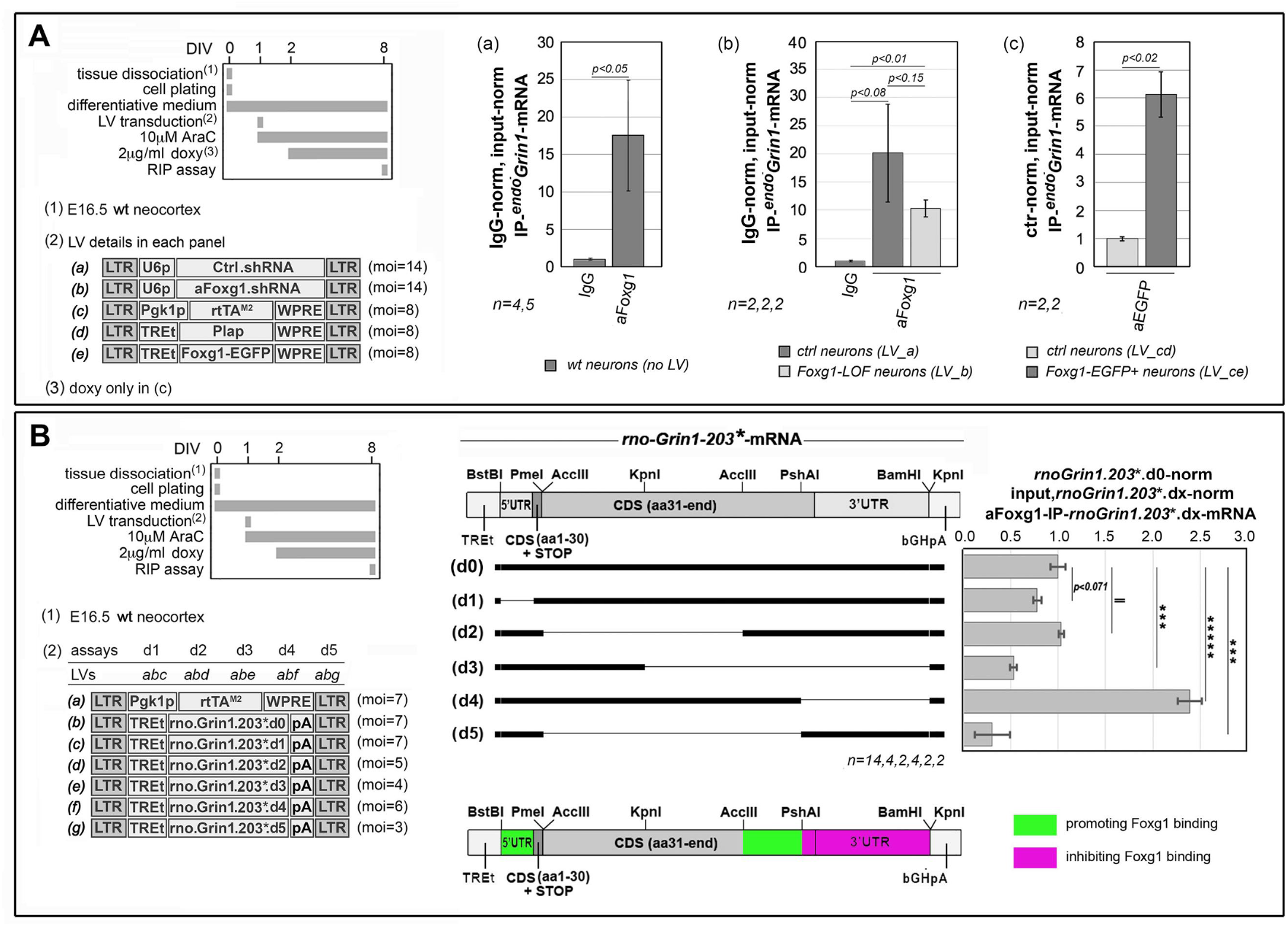
Evaluation of Foxg1-protein/*Grin1*-mRNA interaction in neocortical neurons, by RNA immunoprecipitation (IP) qPCR (qRIP-PCR) assays. (A) Immuno-precipitation of Foxg1-bound, endogenous *Grin1*-mRNA in neocortical neurons. To left, protocols and lentiviral vectors used, to right, results. Anti-Foxg1-IP fraction of endogenous *Grin1*-mRNA in neurons expressing naive (a) or decreased (b) levels of *Foxg1*-mRNA. Results double normalized, against input-RNA and IgG-IP samples. Anti-EGFP-IP fraction of endogenous *Grin1*-mRNA in neurons expressing a lentivector-driven, *Foxg1-EGFP* transgene or a *Plap* control (c). Results double normalized, against input-RNA and control samples. **(B)** Mapping determinants of Foxg1-protein binding on a heterologous *rno-Grin1*-mRNA, encoded by a lentiviral transgene. To left, protocols and lentiviral vectors used, to right, results. Here, a number of partially overlapping deletions were generated starting from the full-length cDNA (d0), by standard molecular cloning techniques, so giving rise to five distinct mutants (d1-d5). To prevent toxicity originating from *chronic*, exaggerated Grin1 expression and potential artifacts stemming from differential protection of *rnoGrin1*-mRNA by translating ribosomes, in all constructs a stop codon was inserted in a fixed position, between codons 30 and 31, so resulting into modified transcripts (*rno-Grin1-203*\* and derivatives). To quantify the impact of each deletion, neural cultures were co-transduced with lentiviral mixes encoding for different combinations of full-length (d0) and mutant (dx) transgenes. Anti-Foxg1-IP fractions of mutant rno*Grin1*-mRNAs, primarily normalized against the corresponding inputs, were diminished by the corresponding IgG-IP backgrounds and finally renormalized against the average full-length fraction. At the bottom, a color-coded cartoon summarizes the positive or negative impact that distinct transcript domains exert on *Grin1*-mRNA/Foxg1-protein interaction. Througout figure, *n* is the number of biological replicates, i.e. independently cultured and engineered preparations, originating from a common neural cell pool. Statistical evaluation of results was performed by one-way ANOVA, one-tailed and unpaired. *** *p*< 0.001, ***** *p*< 0.00001. Errors bars indicate s.e.m.

Next, to tentatively identify *Grin1*-mRNA domains needed to bind Foxg1 protein, we co-transduced murine neocortical neurons with TetON-controlled, intronless transgenes, encoding for the *Rattus norvegicus Grin1*-203 transcript (including exon 20 and orthologous to the murine *Grin1*-201 isoform) and artificially-deleted variants of it. [Within these transgenes, to prevent toxicity induced by chronic *Grin1* overexpression and potential artifacts due to differential protection of *rnoGrin1*-mRNA by translating ribosomes, a stop codon was inserted between codons 30 and 31 (*rnoGrin1*.203*)]. Then, we immunoprecipitated RNA originating from these cultures by anti-Foxg1 and normalized the IP-*Grin1*-mRNA fraction peculiar to each deletion against the IP fraction of full-length *rnoGrin1*.203*.d0. Finally, we critically evaluated the relevance of distinct *Grin1*-mRNA segments to anti-Foxg1 immuno-precipitability (**Figure 7B**). We observed that the two variants missing the AccIII-PshAI fragment at the Grin1-cds 3’ end, *rnoGrin1*.203*.d3 and *rnoGrin1*.203*.d5, specifically displayed a normalized IP fraction far below 1 (0.53±0.04 with *p*<0.002 and *n*=4, and 0.30±0.19, with *p*<0.002 and *n*=2, respectively), pointing to a pivotal role of this fragment in the interaction with Foxg1. Next, the removal of the original 3’UTR, peculiar to *rnoGrin1*.203*.d4, increased the IP fraction up to 2.39±0.13 (with *p*<5*10^-6^ and *n*=2), suggesting that such domain may normally antagonize Foxg1 recruitment to *Grin1*-mRNA. Last, a declining IP trend was also detectable in *rnoGrin1*.203*.d1, missing the 5’UTR (0.78±0.04, with *p*<0.07 and *n*=4), further implicating the 5’UTR in Foxg1 recruitment to *Grin1*-mRNA (**Figure 7B**). Altogether, these results corroborate the specificity of Foxg1/*Grin1*-mRNA interaction and provide a coarse-grained, tentative framework for its articulation.

### Foxg1 is needed to achieve proper homeostatic tuning of neuronal *Grin1*-mRNA translation

*Grin1* is a key player implicated in neuronal plasticity and, in turn, it is subject of intricate, activity-dependent post-transcriptional regulation (Baez et al., 2018; Dörrbaum et al., 2020; Lau and Zukin, 2007; Paoletti et al., 2013). We previously observed that exposing E16.5+DIV8 neocortical cultures to 55mM KCl resulted into a dramatic drop of (n)Grin1 level, that was partially rescued upon transferring the same cultures to a low K^+^-containing medium. This points to a dedicated mechanism taking care of homeostatic translation tuning (our unpublished results).

To evaluate relevance of Foxg1 levels to such tuning, we compared the impact of high extracellular K^+^ on Grin1 translation in *Foxg1*-LOF vs wild-type neural cultures **(Figure 8**, left). As expected, in wild type neurons we confirmed the previously observed colIapse of (n)Grin1 evoked by acute 55mM K^+^ (to 4.4±0.9%o of unstimulated wild-type samples, with p_vs-wt-ctr_<0.0005 and n=3,3), as well as the partial rebound of (n)Grin1 levels upon retransferring cultures to a low K^+^ medium (to 59.0±4.3%, with p_vs-wt-K5’_<0.0002, p_vs-wt-ctr_<0.02, n=3,3,3). Conversely, when *Foxg1* was knocked-down, (a) basal Grin1 translation was reduced to 49.9±3.7% (with p_vs-wt-ctr_<0.006 and n=3,3), (b) the exposure of *Foxg1*-LOF cultures to high K^+^ reduced nGrin1 to 9.1±0.8% (normalized against wt_ctr), with p_vs-Foxg1.LOF-ctr_<0.00003 and n=3,4, and (c) the subsequent re-transfer of these cultures to standard potassium allowed (n)Grin1 to rebound to 27.3±7.2% (again normalized against wt_ctr), with p_vs-Foxg1.LOF-K5’_<0.03, p_vs-Foxg1.LOF-ctr_<0.03 and n=3,4,4 **(Figure 8**, right). In other words, compared to controls, *Foxg1* knock-down dampened the early homeostatic response to high K^+^ by about four-folds (4.4%-vs-100.0% and 9.1%-vs-49.9%, respectively, with p_(genotype/K+)interaction_<0.002, as assessed by 2-ways ANOVA).

**Figure 8.**
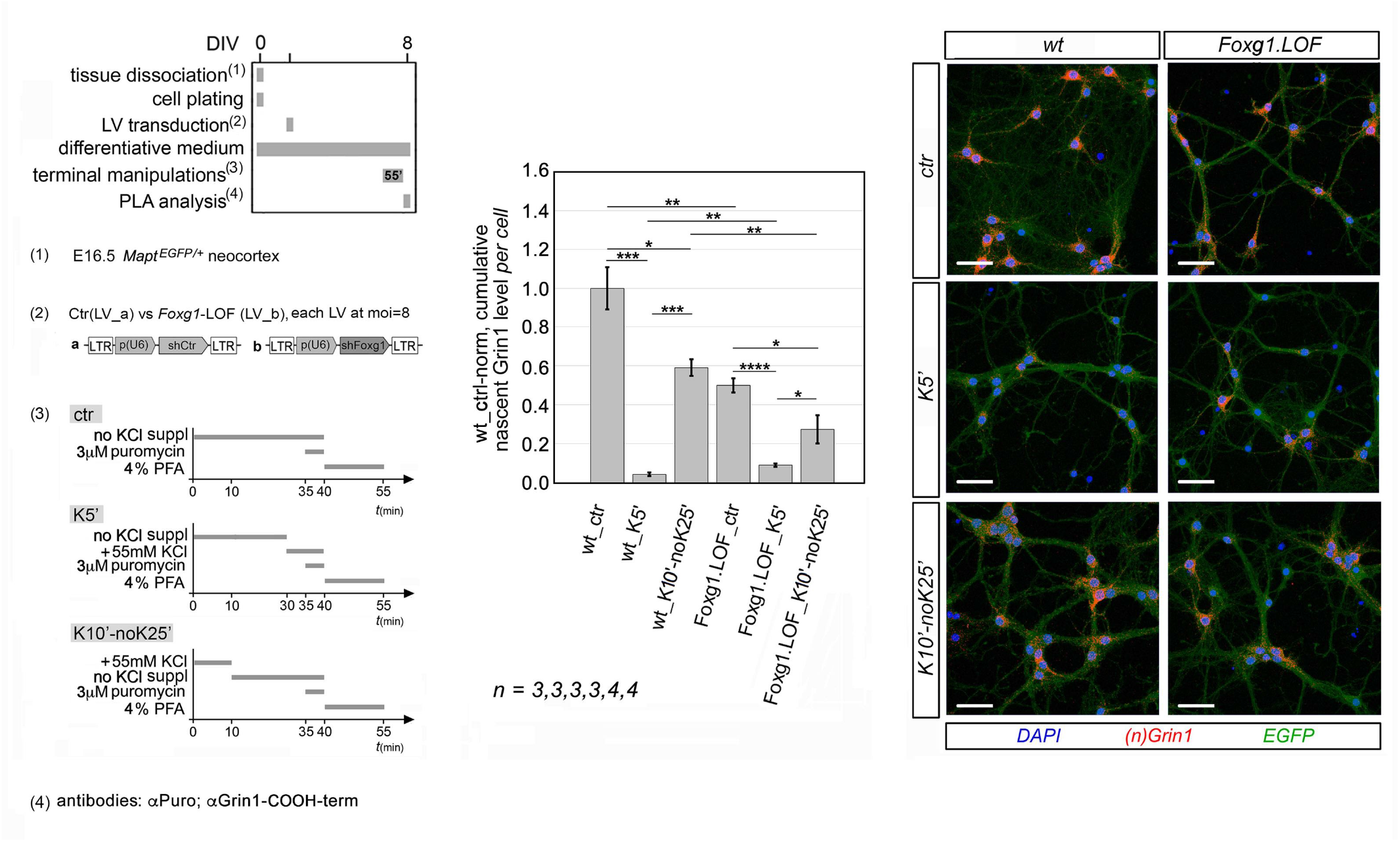
Foxg1 relevance to homeostatic *Grin1*-mRNA translational tuning. To left, protocols (including lentiviruses employed, and operational details of transient neuronal stimulation), to right, results. Impact of *Foxg1*-down-regulation on (n)Grin1 levels, following acute exposure of neocortical neurons to high extracellular potassium (*K5’*) and their return to not-K^+^-supplemented medium (*K10’-noK25’*). *Foxg1* knock-down elicited via shRNA-encoding lentivirus. (n)Grin1 evaluated by anti-Grin1-COOH-term/anti-puromycin-driven proximity ligation assay (PLA). Results normalized against unstimulated controls (*wt_ctr*). Included are examples of primary data. *n* is the number of biological replicates, i.e. independently cultured and engineered preparations, originating from a common cell pool. Scalebars, 50 μm. Statistical evaluation of results performed by t-test, one-tailed and unpaired, and 2-ways ANOVA. * *p*< 0.05, ** *p*< 0.01, *** *p*<0.001, **** *p*<0.0001. Errors bars indicate s.e.m.

In synthesis, we found that Grin1 *de novo* synthesis undergoes a prominent and reversible, homeostatic regulation, and Foxg1 is instrumental to that.

### Widespread impact of *Foxg1* on mRNA engagement to ribosomes

We wondered if *Foxg1* impact on translation is peculiar only to a few genes including *Grin1* or is it a pervasive phenomenon. To get a preliminary insight into this issue, we systematically sequenced ribosome-engaged-mRNA (trapRNAseq) purified from *Foxg1*-OE and control cultures (as in **Figure 2A,left),** and compared it to total-mRNA originating from corresponding sister cultures (totRNAseq) ((Tigani et al., 2020); Artimagnella and Mallamaci, doi: temporarily restricted).

For the sake of simplicity, we took into account trapRNAseq and totRNAseq reads belonging to the only principal isoform of each gene (according to APPRIS annotation) (Rodriguez et al., 2013). We calculated log_2_“expression fold change” values (log_2_FC) peculiar to trapRNA and totRNA samples and evaluated statistical significance of results by DESeq2 software (Love et al., 2014). Next, we scored each gene on the basis of the “log_2_FC(trapRNAseq)-log_2_FC(totRNAseq)” difference (hereafter Δlog_2_FC), as a measure of Foxg1-dependent stimulation of ribosomal mRNA engagement and a presumptive index of Foxg1-driven promotion of its translation. Finally, we evaluated statistical significance of results by Ribodiff software (Zhong et al., 2017).

Upon filtering out low-expressed genes as well as those with p_adj_≥ 0.1’, we found 183 genes with Δlog_2_FC>0.5 (i.e. with Foxg1 presumptively promoting their translation) and 175 genes with Δlog_2_FC<-0.5 (i.e. with Foxg1 presumptively antagonizing their translation). Categorized by Δlog_2_FC value, these genes largely fell within the “1.0 to 1.5” and the “-1.5 to −1.0” intervals (66 and 72 genes, respectively) (**Figure 9A).** As shown by GO analysis, such genes preferentially encode for proteins (a) involved in synaptic signalling, behaviour, memory, fatty acid catabolism, (b) localized at plasmamembrane and synapses, and (c) acting as channels, neurotransmitter receptors, transmembrane transporters, and transcription factor**s (Table S2A**).

**Figure 9.**
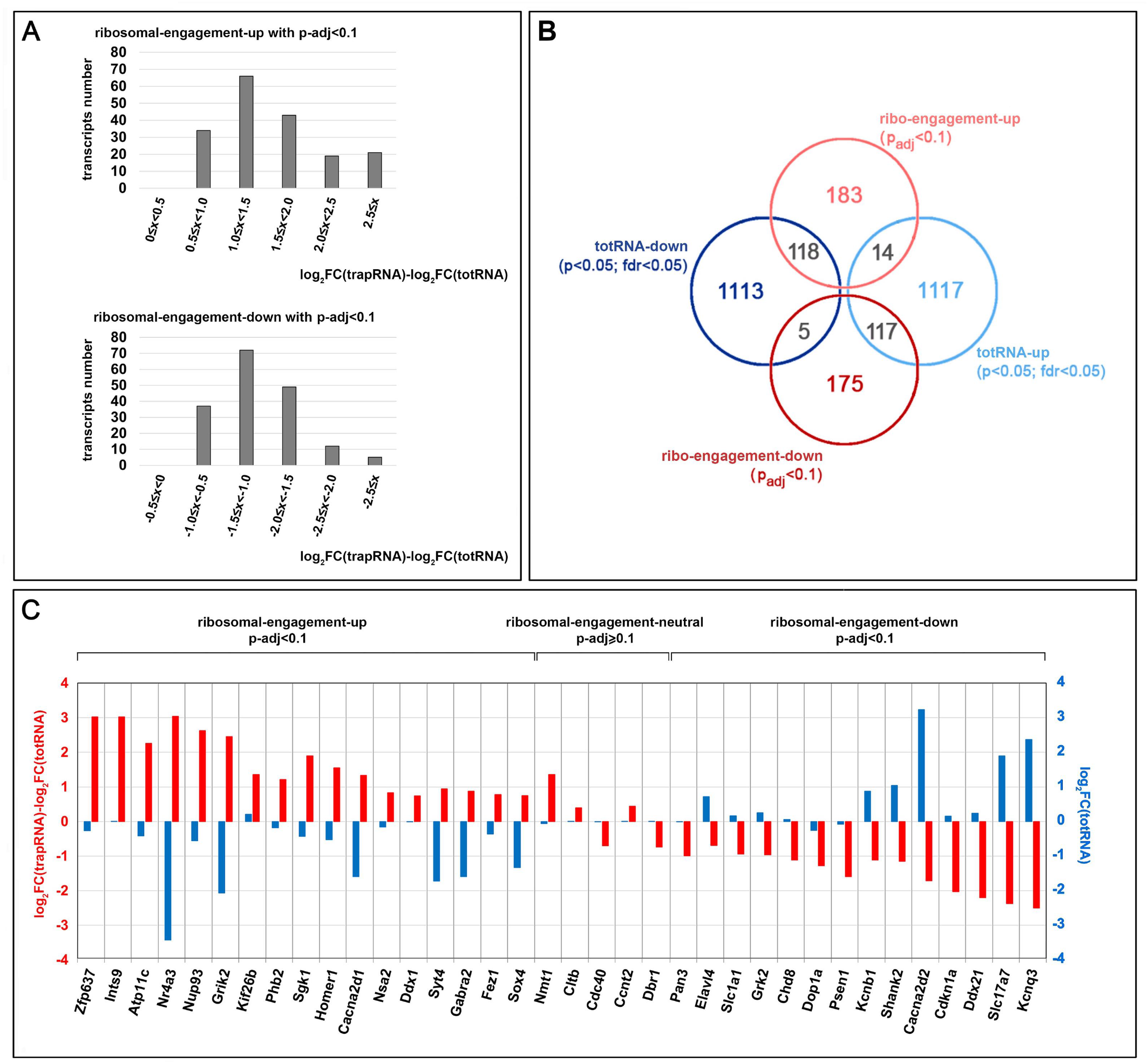
Evaluation of Foxg1 impact on ribosomal mRNA engagement by TRAP-Seq/RNA-Seq. (**A**) Transcripts distribution by “log_2_FC(trapRNAseq)-log_2_FC(totRNAseq)” (or 1′log_2_FC). To top, transcripts with 1′log_2_FC>0 and p_adj_<0.1 (“ribosomal engagement up”), to bottom transcripts with 1′log_2_FC<0 and p_adj_<0.1 (“ribosomal engagement down”), (**B**) Venn’s diagram representation of genes distribution among the four categories: “ribo-engagement-up” (1′log_2_FC>0 and p_adj_<0.1), “ribo-engagement-down” (1′log_2_FC<0 and p_adj_<0.1), “totRNA-up” (log_2_FC>0, with p<0.05 and fdr<0.05), and “totRNA-down” (log_2_FC<0, with p<0.05 and fdr<0.05). (**C**) Representative examples of genes falling in the three categories: “ribosomal engagement up”, “ribosomal engagement neutral” (i.e. with p-adj≥ 0.1), and “ribosomal engagement down”. Here, for each gene plotted are “log_2_FC(trapRNAseq)-log_2_FC(totRNAseq)” and “log_2_FC(totRNAseq)”.

Next, we further classified these genes as for their *Foxg1*-driven totRNA dynamics. We found that among 183 genes with increased ribosomal engagement, as many as 118 displayed reduced totRNA and only 14 increased totRNA. Symmetrically, among 175 genes with decreased ribosomal engagement, 117 and 5 had upregulated and downregulated totRNA, respectively (**Figure 9B**). All that results in a variegated scenario, as shown in **Figure 9C**.

Next, to exclude possible artifactual results originating from Foxg1-dependent alteration of pre-mRNA maturation, we re-analyzed primary totRNA data (Artimagnella and Mallamaci, 2019) by CASH (Wu et al., 2018) and ROAR (Grassi et al., 2016) softwares. Interestingly, we found that, upon Foxg1 overexpression, only (7+14)=21 of the (183+175)=358 genes “with altered ribosomal engagement” mentioned above displayed altered splicing and polyadenylation, respectively (**Table 1**; Artimagnella and Mallamaci, doi: temporarily restricted).

**Table 1.** Distribution of Foxg1-sensitive splicing, Foxg1-sensitive polyadenylation, and mRNA interaction with Foxg1 protein, among gene transcripts characterized by altered ribosomal engagement and/or progression upon Foxg1 over-expression.

|  | total | with altered splicing (3) | with altered polyadenylation (4) | with Foxg1-interacting mRNA(5) |
| --- | --- | --- | --- | --- |
| genes with altered ribosomal engagement (1) | 358 | 7 | 14 | 138 |
| genes with altered ribosomal progression (2) | 328 | 6 | 17 | 93 |

(1) identified by integrated evaluation of totRNA-Seq and TRAP-Seq data; satisfying " $\Delta\log_2FC \neq 0$ ; $p_{adj} < 0.1$ ".
(2) identified on the basis of distribution of TRAP-Seq-reads along every transcript; satisfying " $f_{boi} - zscore \geq 3$ ".
(3) identified by cash software, with $-0.1 \geq \Delta\psi \geq 0.1$ ; $fdr < 0.05$ .
(4) identified by roar software, with $1/1.2 \geq r \geq 1.2$ ; $fdr < 0.05$ .
(5) identified by aFoxg1RIP-Seq, based on the occurrence of Foxg1-protein/mRNA interaction peaks with $aFoxg1/IgG\_enrichment \geq 2$ and $fdr < 0.1$ (mRNAs taken into account sharing $\geq 1$ peak in $\geq 2$ out of 3 biological replicates; calculations restricted to the main isoform of each mRNA).

Then, to further corroborate our findings, we systematically interrogated mRNAs “with altered ribosomal engagement” for a possible interaction with the Foxg1 protein. For this purpose, we relied on sequencing of RNA extracted from E16.5+DIV8 pallial cultures and immunoprecipitated by an aFoxg1 antibody, taking selectively into account exonic reads representative of mature mRNAs (here we referred to the principal splicing isoform, according to APPRIS annotation (Rodriguez et al., 2013)). We monitored the distribution of these reads by Sicer software, comparing aFoxg1-IP samples with IgG-treated controls. Foxg1/mRNA interaction peaks with aFoxg1/IgG_enrichment≥2 and fdr<0.05 resulting from this analysis were further taken into account, and mRNAs sharing ≥1 peak in ≥2 out of 3 biological replicates were considered as interacting with the Foxg1 protein. Specifically, 2857 distinct mRNAs fulfilled this requirement and, interestingly, among the 358 genes “with altered ribosomal engagement” mentioned above, as many as 138 encoded for them (**Table 1**; Artimagnella and Mallamaci, doi: temporarily restricted).

Finally, to primarily validate this approach, we selected *Sgk1* and *Homer1*, namely two genes presumptively undergoing *Foxg1*-OE-driven translational enhancement (Δlog_2_FC equalling +1.89, with padj<0.04, and +1.54, with padj<0.01, respectively) in the face of a significative downregulation of the corresponding mRNAs (−24.94%, with p<10^-4^, and −46.24%, with p<10^-21^, respectively), and we monitored the synthesis rate of their protein products in *Foxg1*-OE neurons by puro-PLA. We found that, compared to controls, such rate was increased in the case of *Sgk1* (by 3.09±0.54-folds, with p<0.006 and n=4,4), and barely shifted upward in the case of *Homer1* (1.12±0.15, p=0.28, n=5,5). Taking into account the underlying, declining mRNA dynamics’, these results unambiguously point to a positive *Foxg1* impact on both *Sgk1* and *Homer1* translation gains. Conversely, the translation rate of *Nmt1,* displaying no statistically significant Δlog_2_FC value or variation in ^tot^mRNA level, was not affected upon *Foxg1* overexpression (**Figure 11A**).

### *Foxg1* impact on ribosomes progression along mRNAs

We further mined our TrapSeq data, aiming at unveiling a possible impact of *Foxg1* expression levels on ribosomal progression along mRNAs. For this purpose, we assumed that, because of random mechanical fragmentation undergone by “ribo-trapped” mRNA during the immuno-precipitation procedure, reads location should provide information about the position occupied by the 60S subunit along the mRNA-cds. Specifically, for each gene, we took into account the principal isoform (according to APPRIS annotation) (Rodriguez et al., 2013), and, for each transcript, we allotted reads to adjacent, 125 bases-wide cds bins. Next, considering each bin/bin boundary as a potential bottleneck for ribosome advancement, we calculated the corresponding ribosome progression index (rpi), as the ratio among reads falling downstream and upstream of such boundary (**Figure 10A**). For each boundary, we averaged rpi’s of the three *Foxg1*-OE replicates and those of the four controls, and we annotated boundaries with log_2_FC(rpi)≥1 and p<0.05 as “boundaries of interest, up” (boi_up_’s). Then, we evaluated the frequency of such boundaries over the full cds (f_boi.up_), as a global, gene-specific index of *Foxg1-*dependent *promotion of* ribosomal progression. In parallel, referring to boundaries with log_2_FC(rpi)≤-1 and p<0.05 (boi_down_’s), we similarly obtained an alternative, gene-specific index of *Foxg1-*dependent *inhibition of* ribosomal progression (f_boi.down_). Finally, to deal with potential false positives originating from random, non-*Foxg1*-dependent variability of ribosome progression, we built all the 34, (4+3)-type permutations of our samples-set and, on each permutation, we repeated the above described analyses. At the end, for each gene, we calculated the f_boi.up_ and the f_boi.down_ z-scores, and we filtered out potential candidates with z-scores<3 (**Figure 10A**; Artimagnella and Mallamaci, doi: temporarily restricted). Among genes with z-scores≥3 (likely undergoing *Foxg1*-dependent modulation of ribosomal progression), 165 harbored at least one boundary with log_2_FC(rpi)≥1 and p<0.05 (boi_up_), conversely 163 displayed at least one boundary with log_2_FC(rpi)≤-1 and p<0.05 (boi_down_). In both cases, the z-score distribution was relatively flat and the correlation between z-score and median log_2_FC(rpi) very low (**Figure 10B**). As expected, moving from low-to high-f_boi_ z-score genes, we found a progressively more pronounced differential distribution of reads, preferentially clustered towards the *Foxg1*-OE cds-3’ in boi_up_-rich genes and towards the control cds-3’ in boi_down_-rich ones (**Figure 10C**). As shown by GO analysis, genes characterized by Foxg1-modulated ribosome distribution along their transcripts preferentially encode for proteins implicated in fatty acid catabolism, Na^+^ binding, AMP binding, and serine/threonine kinase activity (**Table S2B**).

**Figure 10.**
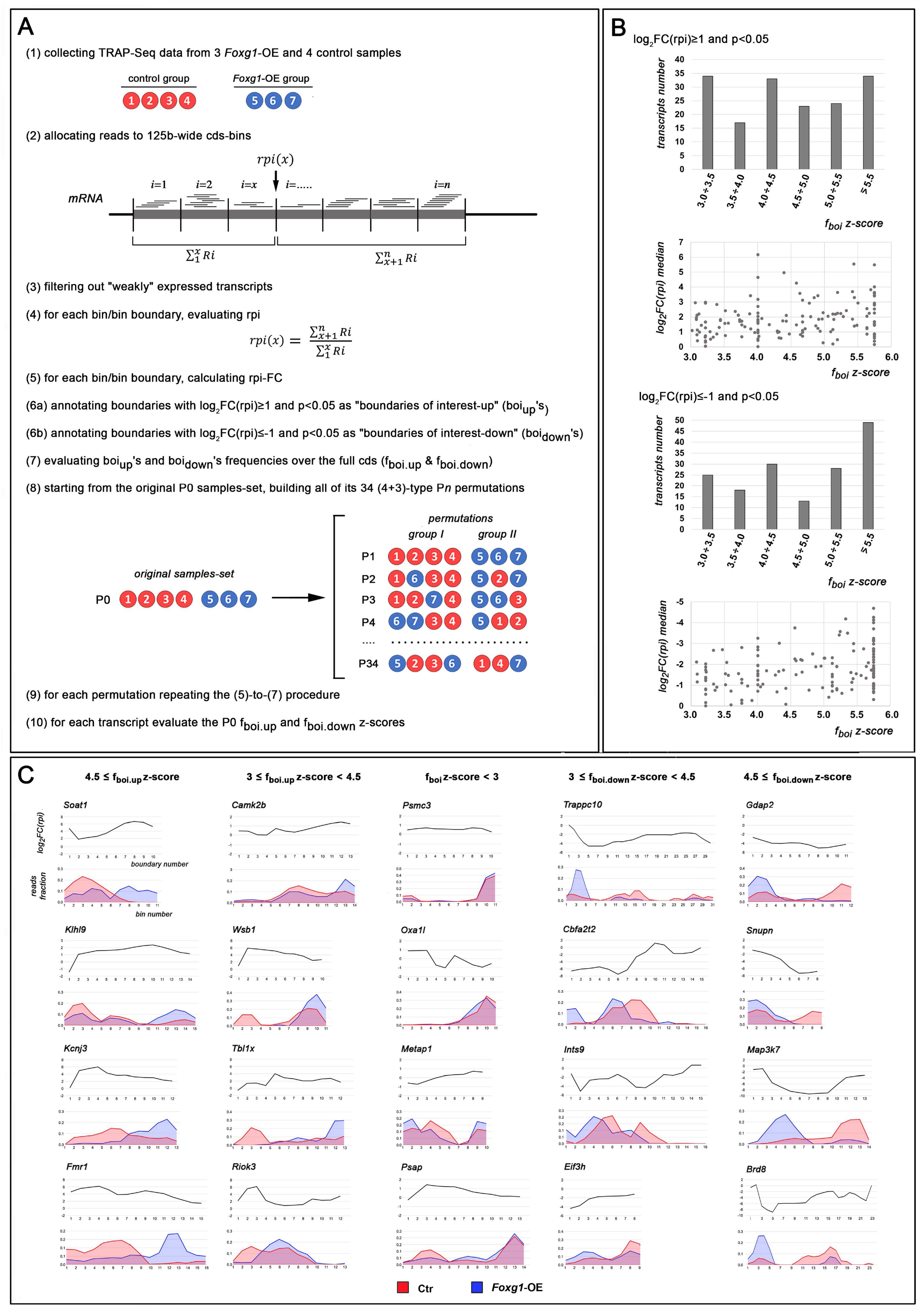
Evaluation of Foxg1 impact on ribosome progression along mRNA by TRAP-seq. **(A)** Step-by-step protocol for *Foxg1*-OE TRAP data mining. Here, *R_i_* indicates the number of reads mapped within the #i bin; *n* is the total bin number; *rpi* is the “ribosome progression index”. **(B)** Transcripts distribution by f_boi_z-score, and correlation between f_boi_z-score and median log_2_FC(rpi), as evaluated for transcripts with “log_2_FC(rpi)≥1, p<0.05” (top graphs) and “log_2_FC(rpi)≤-1, p<0.05” (bottom graphs). **(C)** Examples of genes falling in five different categories, on the basis of their f_boi_z-score values. For each gene, shown are log_2_FC(rpi) progression against bin/bin boundary number (top graph) and reads fraction progression against bin number (bottom graph).

Again to exclude possible artifactual results originating from *Foxg1*-dependent alteration of pre-mRNA maturation, we monitored results of CASH and ROAR profiling of total RNA from *Foxg1*-OE and control neocortical cultures. Interestingly, we found that, upon Foxg1 overexpression, only (6+17)=23 of the (165+163)=328 genes “with altered ribosomal progression” mentioned above displayed altered splicing and polyadenylation, respectively (**Table 1**; Artimagnella and Mallamaci, doi: temporarily restricted).

Then, to further corroborate our findings, we intersected such (165+163)=328 genes “with altered ribosomal progression” with the 2857 ones encoding for Foxg1-interacting mRNAs, previously identified by aFoxg1-RIP/Seq. Interestingly, it turned out that 46/165 and 47/163 genes “with altered ribosomal progression” encodeed for Foxg1-interacting mRNAs (**Table 1**; Artimagnella and Mallamaci, doi: temporarily restricted).

Finally, to preliminarly assess the validity of this approach, we evaluated translation rates of *Camk2b* and *Fmr1*, namely two boi_up_-rich genes (with 3.0 ≤ f_boi.up_ z-score < 4.5, and f_boi.up_ z-score > 4.5, respectively), characterized by diversified reads distributions along their cds’ in *Foxg1*-OE vs control samples (**Figure 10C**), and mRNA expression levels not affected upon *Foxg1* manipulation (Artimagnella and Mallamaci, doi: temporarily restricted). For this purpose, we employed a dedicated puro-PLA run-off assay, similar to the one used for Grin1 (**Figure 4** and **S2**). In the case of *Camk2b*, upon setting the *t_i_* time to 4.5 min, we found that the *t_0_*-normalized decline of the PLA signal (−22.2±0.2% in controls), was remarkably exacerbated in *Foxg1*-OE samples (−47.2±3.5% with p<0.0011 and n=3,3) (**Figure 11B**). This points to an overt positive impact exerted by Foxg1 overexpression on ribosome progression along *Camk2b*-mRNA. It provides a first positive assessment of the predictive power of the bioinformatic strategy we employed. Viceversa, in the case of *Fmr1*, upon setting the *t_i_* time to 6 min, we found that the *t_0_*-normalized decline of the PLA signal (−54.5±8.7% in controls), was *reduced* in *Foxg1*-OE samples (−27.3±8.8% with p<0.027 and n=6,6) (**Figure 11B**). It is possible that, in this case, rather than simply originating from faster holoribosome progression through the very body of the cds, the preferential clustering of ^trap^mRNA reads detectable in the 3’ half of it upon *Foxg1*-OE might reflect some pre-terminal holoribosome accumulation, due to an alternative, 3’-terminal bottleneck evoked by this treatment (**Fig. 10C**).

**Figure 11.**
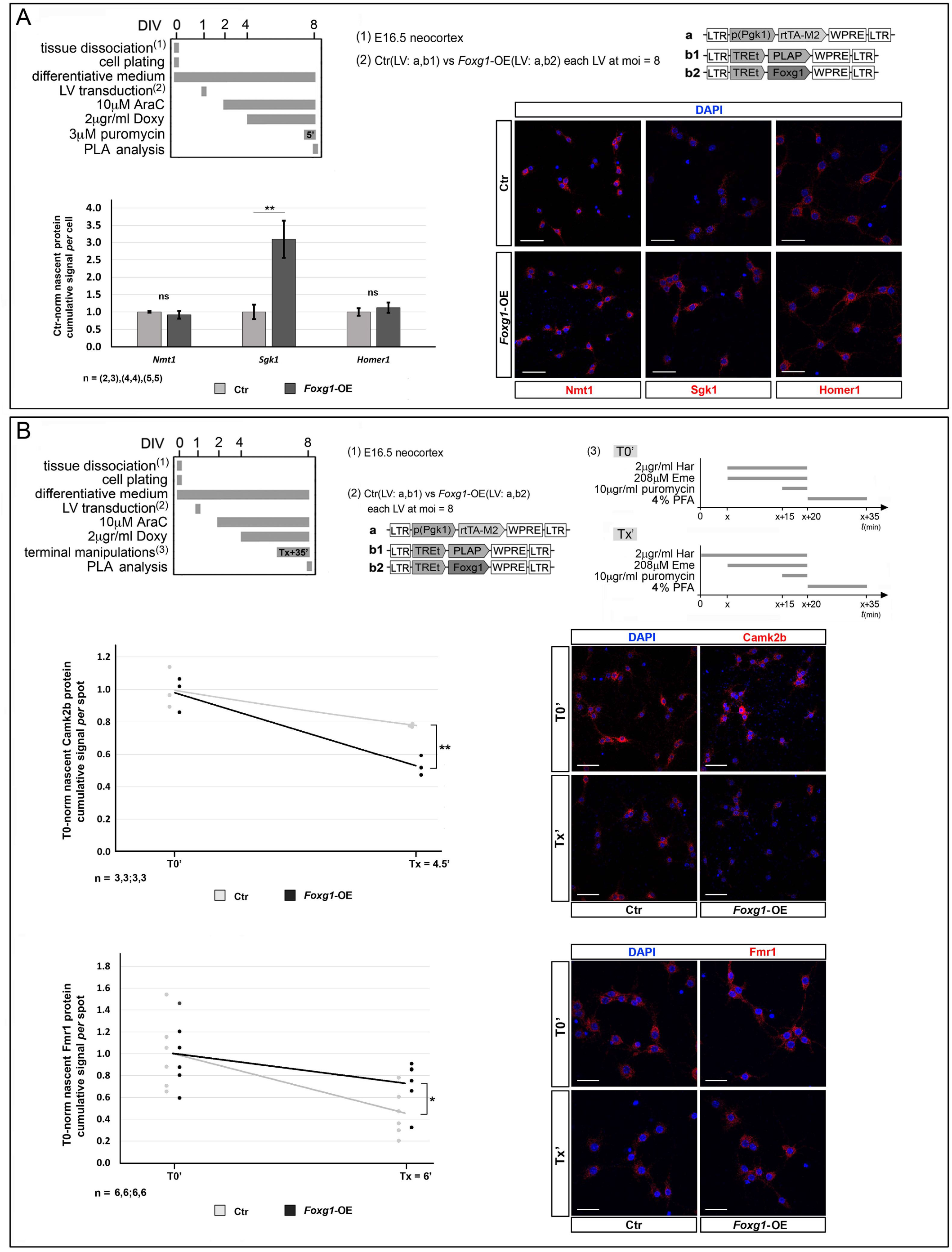
Examples of *Foxg1*-OE TRAP-seq data validation for ribosomal engagement (A) and ribosome progression analysis (B). **(A)** To top, protocols and lentiviruses employed for Foxg1 overexpression, to bottom, results. Graph represents quantitative confocal immunofluorescence (qIF) assessment of nascent Nmt1, Sgk1, and Homer1 protein, performed upon *Foxg1* over-expression, terminal (5’) puromycin administration and subsequent anti-protein/anti-puromycin-driven PLA. As indices of nascent protein levels, shown are average cumulative qIF signals per cell. **(B)** To top, protocols (including lentiviruses employed, and operational details of the translational run-off assay), to bottom, results. Graphs represent progression of nascent Camk2b and Fmr1 levels, evaluated by anti-Camk2b/ or anti-Fmr1/anti-puromycin-driven PLA, upon *Foxg1* down-regulation, in basal conditions (T0’) as well as 4 min 30 sec (case CamK2b) or 6 min (case Fmr1) after 2μg/ml harringtonine (har) blockade of translation initiation (Tx’). In both cases, ribosome progression was subsequently inhibited by 208μM emetine (eme), and nascent polypeptides were terminally labeled by 10μg/ml puromycin (puro). For each genotype, results normalized against (T0’) average values. Superimposed, linear trendlines. Throughout the figure, statistical evaluation of results performed by ANOVA test (one-tail, unpaired). * *p*<0.05; ** *p*<0.01. Errors bars indicate s.e.m. *n* is the number of biological replicates, i.e. independently cultured and engineered preparations, originating from a common neural cell pool. Included are examples of primary PLA data referred to by the corresponding graphs. Scalebars, 50μm.

## DISCUSSION

Here, inspired by the detection of Foxg1 protein in neuritic cytoplasm of neocortical pyramids (**Figure 1**), we investigated its potential implication in translation of selected neuronal genes, and we documented an impact of *Foxg1* on ribosomal engagement of *Grin1*-mRNA (**Figure 2A,B**). Next, we showed that Foxg1 increases Grin1 protein level by enhancing translation of its mRNA, while not ameliorating its stability (**Figure 2C-F** and **3**). Such enhancement was apparently due to increased translational initiation and, possibly, polypeptide elongation (**Figure 4**). Mechanisms underlying these phenomena included Foxg1 protein interaction with Eif4e and Eef1d (**Figure 5** and **6**) as well as with *Grin1*-mRNA (**Figure 7**). Moreover, we found that *Grin1*-mRNA translation undergoes a prominent (and reversible), homeostatic regulation and Foxg1 is instrumental to that (**Figure 8**). Finally, a dedicated TRAP-seq survey showed that, not just peculiar to *Grin1*, functional Foxg1 implication in translation control (both initiation and ribosome progression) apparently is a pervasive phenomenon, affecting hundreds of neuronal genes (**Figures 9-11**).

The localization of Foxg1 in early born, neocortical glutamatergic neurons, outside of the nucleus, had been already reported (Regad et al., 2007). Here we showed that Foxg1 is specifically detectable in soma, dendrites and axons of the majority of pallial pyramids, including mitochondria as well as cytoplasm (**Figure 1**).

Based on higher *Grin1*-mRNA levels in ribosome-engaged-compared to not-ribosome-engaged-RNA of *Foxg1*-OE neurons (**Figure 2B**), we inferred a likely positive impact of Foxg1 on *Grin1* translation. However, hyper-recruitment of an mRNA to ribosomes, as documented by ^anti-Rpl10a^TRAP analysis, does not imply *per se* an increased synthesis of its protein product, but it could alternatively reflect enhanced stalling of the holoribosome on such mRNA, ultimately resulting in *reduced* protein outcome of translation. To disambiguate this issue, we subsequently compared levels of *Grin1* mRNA and protein. We found that higher *Foxg1* level led to an increased “Grin1-protein to *Grin1*-mRNA” ratio (**Figure 2F**) in the absence of Grin1-protein stabilization (**Figure 3C**), as well as to increased puromycin-tagged, nascent Grin1 (**Figure 3A**). That allowed us to definitively validate the aforesaid inference. Intriguingly, a substantial fraction of nascent Grin1 was detected in neurites (**Figure 3A**), consistently with previously reported localization of the corresponding mRNA in these structures (Ainsley et al., 2014; Cajigas et al., 2012; Kügelgen and Chekulaeva, 2020; Middleton et al., 2019). To notice, *Foxg1* did not drive any appreciable, generalized enhancement of translation (**Figure 3B**).

Synthesis rate of a given polypeptide depends not only on initiation of its translation, but it also reflects the speed at which it is elongated. In this respect, combined use of harringtonine and puromycin had already been employed to assay cumulative, proteome-wide polypeptide elongation rates (Argüello et al., 2018). Here, by means of PLA, we re-adapted this method to evaluate elongation rates of *specific* polypeptides in distinctive sub-cellular locales (**Figure S2**). Unfortunately, albeit technically working, this approach did not allow us to document any *Foxg1*-driven change of this rate, as cumulatively evaluated on *all* Grin1 isoforms (**Figure 4A**). Conversely, focusing our attention on a specific *Grin1* isoform, namely the ex20-encoding *mmuGrin1.201*-mRNA, and employing an ad hoc designed sensor, we were able to document a likely, *Foxg1*-driven promotion of polypeptide elongation. However, based on comparative immuno-profiling of two artificial epitopes appended to N- and C-terms of Grin1 (**Figure 4B**), this result requires a cautious interpretation. In fact, albeit providing information on polypeptide elongation rates (**Figure 4B,graph e**), the C/N signal ratio employed in this assay also depends on tags accessibility. This is a parameter hardly sensitive to dynamics of primary protein folding (Kubelka et al., 2004), however potentially sensitive to concomitant, Foxg1-dependent post-transcriptional protein modifications.

As for molecular mechanisms underlying Foxg1 impact on *Grin1* translation, we achieved multiple pieces of evidence pointing to it as a “translation modulator”. In fact, beyond its detection in neuronal cytoplasm (**Figure 1**), we found that Foxg1 interacts with Eef1d and Eif4e, and partial inhibition of its interaction with the latter resulted in a substantial decline of Grin1 translation (**Figure 5** and **6**). Moreover, Foxg1 binds to *Grin1*-mRNA (**Figure 7**). To note, our PLA-based investigation of Foxg1/Eef1d association confirmed results of previous high-throughput mass spectrometry (MS) screenings in HEK293T and N2A cells (Li et al., 2015; Weise et al., 2019). Conversely, Foxg1 interaction with Eif4e, we proved by both IP-WB analysis and PLA, is novel. Similarly, while an interaction of Foxg1 with ncRNAs (miRNA precursors) has been already reported (Weise et al., 2019), our finding of neuronal Foxg1 interaction with mRNA has been not yet documented. Needless to say, Foxg1 association to Eif4e and Eef1d resonates with presumptive Foxg1 implication in translation initiation and polypeptide elongation, respectively (Jackson et al., 2010; Pelletier and Sonenberg, 2019).

It has been shown that acute stimulation of hippocampal pyramids by high extracellular potassium may evoke a fast increase of cap-dependent translation (Srivastava et al., 2012). Moreover, *Grin* genes - which encode for subunits of the heteromeric NMDA receptor - undergo an intricate, multi-step regulation, needed for proper setting of integrative properties of neocortical pyramids (Paoletti et al., 2013). In this context, specific and reversible, high K^+^-driven *downregulation* of Grin1 translation (**Figure 8**) might represent the experimental correlate of specific physiological mechanisms contributing to homeostatic scaling of neuronal response to glutamate (Dörrbaum et al., 2020).

Next, *Foxg1* has been recently shown to promote activity and excitability of neocortical neurons, largely via a profound impact on their transcriptome (Tigani et al., 2020). Consistently, *Foxg1*-depleted hippocampal neurons display reduced NMDA currents and defective long term potentiation (LTP) (Yu et al., 2019). In this respect, *Foxg1*-dependent modulation of *Grin1* translation (**Figures 3** and **8**) might be a key mechanism concurring to both these effects.

Finally, we have recently shown that *Foxg1* is transiently upregulated by neuronal hyperactivity (Fimiani et al., 2016; Tigani et al., 2020). In this way, delayed, *Foxg1*-mediated promotion of Grin1 translation, following episodes of intense electrical activity, might contribute to normal dynamic shaping of pyramid excitability, and its absence might impair neuronal plasticity, contributing to major cognitive deficits of *FOXG1*-haploinsufficient patients (Mitter et al., 2018; Vegas et al., 2018; Wong et al., 2019).

The involvement of a neurodevelopmental transcription factor in control of mRNA translation is not novel. It has already been reported in a few cases, including those of Bicoid (Niessing et al., 2002), Emx2 (Nédélec et al., 2004), and En2 homeoproteins (Brunet et al., 2005). In our case, we found that Foxg1 implication in translation is not limited to *Grin1* only, but it likely is a pervasive phenomenon, affecting hundreds of genes (**Figures 9** and **10**), among which a large subset encoding for proteins involved in neuronal metabolism and activity (**Table S2**). In a subset of cases we got robust evidence of physical interaction between mRNAs subject of *Foxg1* translational control and the Foxg1 protein (**Table 1),** suggesting that - at least in such cases - the latter may work as a “translation factor”. To note, the number of mRNA interactors of Foxg1, 2857, largely exceeded the number of those further undergoing Foxg1 control of translation, 138+46+47=231, pointing to a likely Foxg1 involvement in other aspects of post-transcriptional gene tuning.

Remarkably, albeit our quantification of ribosome engagement and progression was intentionally restricted to the principal isoform of each polypeptide-encoding transcript, as such isoform often shares a large subset of its exon/intron architecture with minor ones, a number of reads originating from the latter was likely mis-attributed to the former. Next, since different translational gains may apply to distinct isoforms, a change in isoform ratio originating from *Foxg1*-dependent modulation of alternative splicing and/or polyadenylation might have resulted into an artifactual impact of *Foxg1* overexpression on ribosome engagement and progression parameters. To address this issue, we re-analyzed primary totRNA-Seq data from *Foxg1*-OE and control cultures by CASH and ROAR softwares. It turned out that only a minority of presumptive translational targets of *Foxg1* regulation underwent *Foxg1*-dependent modulation of splicing and/or polyadenylation patterns (**Table 1**; Artimagnella and Mallamaci, doi: temporarily restricted), therefore allowing us to fix this concern. To note, while running these controls, we documented an additional impact of Foxg1 on two steps of pre-mRNA maturation, i.e. splicing and polyadenylation. We will address these novel aspects of Foxg1 biology in a forthcoming, dedicated study.

As said above, we have shown that integrated mining of trap- and total-RNA data can provide evidence of Foxg1 control over ribosomes engagement to mRNA, while binning of trap-RNA reads may unveil Foxg1 control of ribosomes progression along it. However, the interpretation of results originating from such approaches deserves caution. This applies firstly to the evaluation of the Δlog_2_FC parameter. For example, rather than simply reflecting *enhanced translation initiation*, Δlog_2_FC values above 0 might also alternatively originate from pronounced ribosome stalling by the kozak motif. Consistently with this prediction, we found that 8 transcripts out of 183 ones with Δlog_2_FC>0 (see **Figure 9B**), were also characterized by “average_log_2_FC(rpi)<0 and boi.down-z-score>3”. In such cases, Foxg1 could actually *limit* baseline translation (possibly paving the way to subsequent, prompt completion of it, upon the arrival of due inputs). In a symmetrical way, Δlog_2_FC values below 0 might originate from extremely fast ribosome progression along the cds and anticipated detachment from it. Again, consistently with this prediction, we found that 2 transcripts out of 175 ones with Δlog_2_FC<0 (see **Figure 9B**), were also characterized by “average_log_2_FC(rpi)>0 and boi.up-z-score>3”. Here, Foxg1 might elicit a *hyperacute arousal of translation*, just by relieving ribosomal stalling. Finally, beyond Δlog_2_FC issues, even the *rpi* (**Figure 10A**) has an intrinsically limited predictive power, similarly to the corresponding Ribo-seq parameters (Ingolia et al., 2009). In fact, it provides only a static snapshot of presumptive ribosome distribution along mRNA and no direct information about the actual speed at which ribosomes move. For all these reasons, TRAP-seq data mandatorily require to be integrated by experimental investigation of the *actual* rate at which polypeptides of interest are synthesized.

Prompted by these considerations, we challenged results of our total/TRAP-seq analyses, firstly by assessing translation rates of *Sgk1*-and *Homer1*-mRNA, namely two transcripts apparently undergoing *Foxg1*-driven promotion of ribosome engagement. In both cases, integrated evaluation of puro-PLA results and ^tot^mRNA dynamics’ pointed towards an overt increase of the translational gain evoked by *Foxg1* overexpression (**Fig. 11A**). Next, we focused our attention on *Camk2b*-and *Fmr1*-mRNA, namely two transcripts showing 3’-ward displaced ^trap^mRNA reads upon *Foxg1* overexpression. We measured the temporal decline rate of their translation upon harringtonin blockade of translation initiation, as an index of ribosome progression along their cds’. For this purpose, we employed a “puro-PLA run-off” assay, i.e a novel method we developed to evaluate ribosome advancement speed along specific mRNA-cds’ (**Figure S2**). As expected, this method provided evidence of faster ribosomal progression through *Camk2b*-cds, upon *Foxg1*-OE. In case of *Fmr1*, it conversely pointed to the alternative emergence of a novel ribosomal pausing site, likely evoked by *Foxg1-OE* towards the 3’ end of *Fmr1*-cds (**Figure 11B**). Of course, all that was only a proof-of-principle, “wet validation” of our procedure, which needs to be corroborated by further experimental work. This will be the subject of dedicated follow-up studies.

To note, albeit providing us with only coarse-grained information about ribosome location along mRNA, our reanalysis of “cheap” TRAP-seq data allowed us to identify as many as >300 genes characterized by a robustly diversified ribosome association to distinctive mRNA regions, dependent on *Foxg1* expression levels (**Figure 10B,C**). This suggests that, aiming to explore the impact of specific independent variables on ribosomal progression rates, mining publicly available TRAP-seq data might be an advisable approach, prior to moving to more expensive, state-of-art Ribo-seq profiling.

Intriguingly, in a number of cases including *Grin1*, we also found that a large subset of genes characterized by statistically significant Δlog_2_FC>0 displayed a robust downregulation of their total-mRNA (118/183), and viceversa for those with Δlog_2_FC>0 (117/175) (**Figure 9B,C**, and Artimagnella and Mallamaci, doi: temporarily restricted). In the former case, reminiscent of activity-driven regulation of Npas4 and Arc (Brigidi et al., 2019; Steward et al., 2018), the very same effector, Foxg1, might promote a rapid arousal of the protein, while however limiting the temporal duration of its overexpression. In the latter, Foxg1 could conversely elicit a slow protein upregulation followed by a delayed, fast decrease of it. Evolutionarily speaking, the multilevel regulation of the same target genes by a unique molecular effector is a rare and thermodynamically demanding phenomenon. Such phenomenon could ease the portability/selectability of temporally structured expression programs (in the minutes/hours range).

Mainly known as a transcription factor patterning the terminal brain and ruling its histogenesis, Foxg1 has been also detected outside the nucleus, however biological meaning of that has been clarified only to a limited extent (reviewed by (Hou et al., 2020)). Our implication of Foxg1 in translational control of a large number of neuronal genes points to this factor as a multi-scale, temporal modulator of neocortical pyramid plasticity. Interesting per se as well as for its profound neuropathogenic implications, this issue will be specifically investigated in a future, dedicated follow-up study.

## MATERIALS AND METHODS

### Animal handling

#### Permissions

Animal handling and subsequent procedures were in accordance with European and Italian laws (European Parliament and Council Directive of 22 September 2010 (2010/63/EU); Italian Government Degree of 04 March 2014, no.26). Experimental protocols were approved by SISSA OpBA (Institutional SISSA Committee for Animal Care) and authorized by the Italian Ministery of Health (Auth. No 22DAB.N.4GU).

#### Rodent strains, animal handling and neural tissue dissection

In this study, the following rodent models were employed:

– wild type (*wt*) CD1 strain mice [purchased from Envigo Laboratories, Italy];

– transgenic *Gt(ROSA)26Sor^tm1.1(CAG-EGFP/Rpl10a,-birA)Wtp^/J* mice, throughout the text referred to as *Rpl10a*^EGFP-Rpl10a/+^ (Zhou et al., 2013) [founders purchased from Jackson Laboratories, USA, Jax #022386; transgenic line maintained according to Jackson’s instructions];

– transgenic *Mapt^EGFP/+^* mice (Tucker et al., 2001) [founders purchased from Jackson Laboratories, USA, Jax #004779; transgenic line transferred to CD1 background (>20 backcrossing generations)];

– *wt* Wistar rats [generated at the SISSA animal facility starting from founders purchased from Envigo Laboratories, Italy].

Mutant mouse embryos were obtained by crossing *wt* females to mutant or *wt* males, and were staged by timed breeding and vaginal plug inspection. Pregnant dams were killed by cervical dislocation. *Rpl10a*^EGFP-Rpl10a/+^ and *Mapt^EGFP/+^* mouse embryos were distinguished from their *wt* littermates by UV lamp inspection.

Rat pups were anesthetized with CO_2_ and sacrificed by decapitation.

Mouse and rat neural tissue was dissected out in sterile ice-cold 1X-phosphate buffered saline (PBS) supplemented with 0.6% D-glucose (Sigma), under sterile conditions.

### Plasmids and lentiviruses

Plasmids employed in this study include:

– LV_pU6-shFoxg1 (Sigma SHCLND-NM_008241, TRCN0000081746); see Figures 2,3C,4B and 7A.

– LV_pU6-shFoxg1-DPuroR [built by removing the SacII/SacII fragment, including the 5’ end portion of puromycin resistance cds and its upstream hPGK-promoter, from “LV_pU6-shFoxg1”; annotated as “LV_pU6-shFoxg1” in Figures 3A-B,4A, 8 and 11].

– LV_pU6-shCtrl (Chiola et al., 2019).

– LV_pPgk1-rtTA2S-M2 (Spigoni et al., 2010).

– LV_pPgk1-EGFP (Brancaccio et al., 2010).

– LV_TREt-Foxg1 (Raciti et al., 2013).

– LV_TREt-PLAP (Falcone et al., 2019).

– LV_pPgk1-mCherry (Falcone et al., 2019).

– LV_pPgk1-3xF-wt.mmuFoxg1^aa357-381^-V5 [built by replacing the AgeI/SalI EGFP-cds fragment of LV_pPgk1-EGFP, by the AgeI/SalI wt.mmuFoxg1^aa357-381^-V5 module (as detailed in **Table S3**)]

– LV_pPgk1-3xF-scr.mmuFoxg1^aa357-381^-V5 [built by replacing the AgeI/SalI EGFP-cds fragment of LV_pPgk1-EGFP, by the AgeI/XhoI scr.mmuFoxg1^aa357-381^-V5 module (as detailed in **Table S3**)]

– LV_TREt-Foxg1-EGFP [built by replacing the SrfI/ApaI fragment of LV_TREt-Foxg1 (including the last 161nt of Foxg1-cds) with the “SrfI-Foxg1(cds-3’term)-EGFP-ApaI” fragment, detailed in **Table S3**].

– LV_CMV-Flag-eIF4E (lentivirus of second generation; Addgene plasmid #38239).

– CMV-Flag-GFP (Addgene plasmid #60360).

– CMV-Flag-Gephyrin (a gift from E.Cherubini’s Lab).

– LV_CMV-EEF1G-V5 (DNASU Plasmid Repository, HsCD00434091).

– LV_CMV-EEF1D-V5 (DNASU Plasmid Repository, HsCD00444454).

– LV_CMV-PUM1-V5 (DNASU Plasmid Repository, HsCD00438817).

– LVrc_TREt-pl-BGHpA [built by replacing the “pPgk1-EGFP-WPRE” fragment of “LV_pPgk1-EGFP” by a “TREt-polylinker-BGHpA” stuffer, in a 3’LTR-to-5’LTR orientation (as detailed in **Table S3**)]

– LVrc_TREt-Flag-rnoGrin1-203-HA [built by tagging the *rnoGrin1-*203 cds by a 3xFlag N-term epitope-cDNA and 3xHA C-term epitope-cDNA (“Flag-rnoGrin1-203-HA” fragment, detailed in **Table S3**) and transferring the resulting double-tagged *rnoGrin1-203*-cDNA into ^filled-in^XhoI/XbaI-cut “LVrc_TREt-pl-BGHpA” vector].

– LVrc_TREt-rnoGrin1-203*.full [built by introducing a STOP codon and a polylinker after codon 30 of *rnoGrin1-203* cDNA and transferring the resulting “rnoGrin1-203*.full” fragment (detailed in **Table S3**) into ^filled-in^XhoI/XbaI-cut “LVrc_TREt-pl-BGHpA” vector].

– LVrc_TREt-rnoGrin1-203*.d1 (5’utr deletion) [built by replacing the BstBI/PmeI fragment of “LVrc_TREt_rnoGrin1-203*.full” with the synthetic “BstBI-GAGCTC-(rnoGrin1-203*:1-30aa)-STOP-PmeI” module].

– LVrc_TREt-rnoGrin1-203*.d2 (cds1 deletion) [built by removing the AccIII/AccIII fragment from “LVrc_TREt_rnoGrin1-203*.full”].

– LVrc_TREt-rnoGrin1-203*.d3 (cds2-3’utr deletion) [built by removing the KpnI/KpnI fragment from “LVrc_TREt_rnoGrin1-203*.full”].

– LVrc_TREt-rnoGrin1-203*.d4 (3’utr deletion) [built by removing the PshAI/BamHI fragment from “LVrc_TREt_rnoGrin1-203*.full”].

– LVrc_TREt-rnoGrin1-203*.d5 (cds3 deletion) [built by removing the PmeI/PshAI fragment from “LVrc_TREt-rnoGrin1-203*.full”].

Starting from a subset of these plasmids, self-inactivating lentiviral vectors (LV) were generated and titrated as previously described (Brancaccio et al., 2010).

### Primary cortico-cerebral cell cultures

Cortical tissue from E16.5 mice was chopped to small pieces for 5 minutes (min), in the smallest volume of ice-cold 1X PBS - 0,6% D-glucose - 5mg/ml DNaseI (Roche #10104159001) solution. After chemical digestion in 2.5X trypsin (Gibco #15400054) - 2mg/ml DNaseI (Roche) for 5 min, and trypsin inhibition with DMEM-glutaMAX (Gibco) - 10% FBS (Euroclone) - 1X Pen-Strep (Invitrogen), cells were spinned down and transferred to differentiative medium [Neurobasal-A (Gibco), 1X Glutamax (Gibco), 1X B27 supplement (Invitrogen), 25μM L-glutamate (Sigma), 25μM β-Mercaptoethanol (Gibco), 2% FBS (Euroclone), 1X Pen/Strept (Invitrogen), 10pg/ml fungizone (Invitrogen)]. Cells were counted and plated as follows:

a. in case of RNA profiling (totalRNA-, TRAP- and RIP-qRTPCR assays) and western blot experiments, cells were plated onto 0.1mg/ml poly-L-Lysine (Sigma #P2636) pre-treated 12-multiwell plates (Falcon) at 8×10^5^ cells/well in 0.6-0.8 ml differentiative medium;
b. in case of immunofluorescence and PLA assays, cells were plated onto 0.1mg/ml poly-L-Lysine pre-treated 12mmØ glass coverslips in 24-multiwell plates (Falcon) at 1×10^5^ cells/well in 0.6-0.8 ml differentiative medium.

In general, when required and as indicated in each Figure, lentiviral infection was done at DIV1; TetON regulated transgenes were activated by 2μg/ml doxycycline (Clontech #631311) administration; 10μM Cytosine β-D-arabinofuranoside (AraC; Sigma #C6645) was acutely added to the medium at DIV1. Cells were kept in culture for 8 days.

### Primary hippocampal cell culture and their live-imaging

Hippocampal tissue from P2 Wister rats was chopped to small pieces and digested by 5mg/ml trypsin (Sigma #T1005) - 5mg/ml DNaseI for 5 min. Dissociated cells were plated at 4×10^4^ cells/ml onto 35mm glass dishes (Ibidi) pre-treated with 0.5mg/ml poly-D-lysine (Sigma) for 1 h at 37°C, in Minimum Essential Medium (MEM) with GlutaMAX™ (Gibco) supplemented with 10% FBS, 0.6% D-glucose, 15 mM Hepes, 0.1mg/ml apo-transferrin, 30μg/ml insulin, 0.1 μg/ml D-biotin, 1 μM vitamin B12 (all from Sigma), and 2.5 μg/ml gentamycin (Invitrogen). At DIV2, hippocampal neurons were engineered with “LV_pPgk1-rtTA2S-M2” and “LV_TREt-Foxg1-EGFP” transgenes. At DIV4, the expression of *Foxg1*-*EGFP* chimera was activated by administration of 2μg/ml doxycycline. Finally, at DIV7, 50nM Mitotracker dye (Life Technologies #M7512) was added to hippocampal cultures, and confocal images were acquired after 30 min. Live fluorescent imaging was done with a confocal microscope (NIKON A1R) equipped with 488nm and 594nm laser excitation light and a 60×oil immersion objective (N.A. 1.40), keeping samples at 37°C, 5% CO_2_, and 95% humidity.

### HEK293T cell cultures

HEK293T cells were used for lentivirus production, lentivirus titration (Brancaccio et al. 2010), as well as to evaluate protein-protein interactions via co-immunoprecipitation (co-IP) and proximity ligation assay (PLA). HEK293T cells were cultured in DMEM-glutaMAX - 10% FBS - 1X Pen-Strep, on 6-multiwell plates at 1.2×10^6^ cells/well (for co-IP assays) or on 0.1mg/ml poly-L-Lysine pre-treated 12mmØ glass coverslips in 24-multiwell plates at 3×10^5^ cells/well (for PLA assays). In all cases, cells were transfected by LipoD293 (SignaGen laboratories #SL100668) at DIV1, according to manufacturer’s instructions. Cells were further kept in culture for 3 and 2 days, for co-IP and PLA assays, respectively, and finally analyzed.

### Immunofluorescence assays

Neural cell cultures were fixed by ice-cold 4% PFA for 15-20 min and washed 3 times in 1X PBS. Samples were subsequently treated with blocking mix (1X PBS; 10% FBS; 1mg/ml BSA; 0.1% Triton X-100) for at least 1 hour at room temperature (RT). After that, incubation with primary antibodies was performed in blocking mix, overnight at 4°C. The day after, samples were washed 3 times in 1X PBS - 0.1% Triton X-100 for 5 minutes and then incubated with secondary antibodies in blocking mix, for 2 hours at RT. Samples were finally washed 3 times in 1X PBS - 0.1% Triton X-100 for 5 minutes, and subsequently counterstained with DAPI (4’, 6’-diamidino-2-phenylindole) and mounted in Vectashield Mounting Medium (Vector). The following primary antibodies were used: anti-Tubβ3, mouse monoclonal, (clone Tuj1, Covance #MMS-435P, 1:1000); anti-Foxg1, rabbit polyclonal (gift from G.Corte, 1:200); anti-Psd95, mouse monoclonal (clone 6G6-1C9, Abcam #ab2723, 1:500); anti-Smi312, mouse monoclonal (Abcam #ab24574, 1:1000); anti-Flag, mouse monoclonal (clone M2, Sigma #F1804, 1:1000); anti-HA, rat monoclonal (clone 3F10, Roche #11867423001, 1:500); anti-puromycin, mouse monoclonal (clone 12D10, Millipore #MABE343, 1:4000). Secondary antibodies were conjugates of Alexa Fluor 488, and Alexa Fluor 594 (Invitrogen, 1:600).

### Proximity ligation assays (PLAs), puro-PLAs, puro-PLA-run-off assays

PLA assays were performed according to manufacturer’s instructions (Duolink™ PLA Technology, Sigma). Briefly, cells were fixed for 15-20 min in ice-cold 4% PFA, washed 3 times in 1X PBS, permeabilized in 1X PBS - 0.1% Triton X-100 for 1h at RT, blocked for 1 h at 37°C in Duolink blocking buffer and incubated for 3h/overnight at RT with mouse and rabbit primary antibodies (as indicated in the corresponding Figures). Afterwards, samples were washed 3 times for 5 min in Duolink buffer A, and then incubated for 1 h at 37°C with Duolink anti-mouse MINUS and anti-rabbit PLUS probes, both co-diluted 1:5 in Duolink antibody dilution buffer. Next, samples were washed 3 times for 5 min in buffer A, incubated for 30 min at 37°C in Duolink ligase diluted 1:40 in 1X ligation buffer, washed again 3 times in buffer A, and incubated for 100 min at 37°C in Duolink polymerase diluted 1:80 in 1X green or red amplification buffer. Finally, samples were washed 2 times for 10 min in Duolink buffer B, and 1 time in 1:100 buffer B for 1 min, and mounted in Duolink mounting medium with DAPI. Then, by 48 hours, confocal images were acquired.

Puro-PLA samples (tom Dieck et al., 2015), were prepared as indicated in the corresponding Figures and schematized in Figure S2. Briefly, cortico-cerebral cells were pulsed for 5 min with 3μM puromycin (Sigma #P8833) or with 1X PBS (negative control), and, immediately afterwards, fixed in ice-cold 4% PFA for 15 min. Then, they were processed by standard PLA, as above.

Puro-PLA-run-off DIV8 samples were prepared as indicated in the corresponding Figures. In particular, before terminal puromycin labeling, cells were cumulatively exposed to 2μg/ml harringtonine (Abcam #ab141941), for 20’ or (20+x)’ depending on the “T0’” or the “Tx’” branch of the protocol, and 208μM emetine (Sigma #E2375), for 20 min. Finally, during the last 5’ of harringtonin/emetin treatment, unfinished polypeptides were labeled via further medium supplementation by 10μg/ml puromycin. Immediately afterwards, samples were fixed in ice-cold 4% PFA for 15 min and processed for standard PLA, as above.

The following primary antibodies were used: anti-Grin1 COOH-term, rabbit monoclonal [EPR2481(2)] (Abcam #ab109182, 1:500); anti-Grin1 NH2-term, rabbit polyclonal (Alomone #AGC-001, 1:500); anti-Camk2b, rabbit polyclonal (GeneTex #GTX133072, 1:500); anti-Nmt1, rabbit polyclonal (GeneTex #GTX130852, 1:500); anti-Sgk1, rabbit polyclonal (GeneTex #GTX54726, 1:200); anti-Homer1, rabbit polyclonal (GeneTex #GTX103278, 1:300); anti-Fmr1 rabbit monoclonal (Huabio #ET1703-70, 1:500); anti-Foxg1 ChIP-grade, rabbit polyclonal (Abcam #ab18259, 1:500); anti-puromycin, mouse monoclonal (clone 12D10, Millipore #MABE343, 1:1000); anti-eIF4E, mouse monoclonal (clone 5D11, Thermofisher #MA1-089, 1:100); anti-V5, mouse monoclonal (SV5-Pk1, Abcam #ab27671, 1:1000); anti-Flag, mouse monoclonal (clone M2, Sigma #F1804, 1:1000).

### Neuronal stimulation assays

Cortico-cerebral cultures were set up as described above (to see “Primary cortico-cerebral cell cultures”) and as detailed in **Figure 8**. Specifically, their terminal DIV8 manipulation was as follows. “K5” samples were pulsed with 55mM KCl-supplemented medium for 5 min. “K10-noK25” samples were firstly pulsed with 55mM KCl-supplemented medium for 10 min and then transferred to a conditioned medium, taken from unstimulated sister cultures, for 25 min. “Ctr” samples were kept in standard, not KCl-supplemented medium. Next, “K5”, “K10-noK25” and “Ctr” cells were all pulsed by 3μM puromycin for 5 min, and, immediately afterwards, fixed in ice-cold 4% PFA for 15 min.

### Photography and image analysis

#### Basic immunofluorescence

(**Figures 1A-F** and **3B**) αFoxg1-, αTubb3-, αPSd95-, αSmi312- and αPuro-immunoprofiled cells were photographed by a Nikon C1 confocal system equipped with 40X-oil objective. Photos were collected as 3μm Z-stacks (step = 0.3μm). Upon Z-stack flattening (max version), pictures were imported into Adobe Photoshop CS6, for subsequent processing. (**Figure 4B**) αFlag- and αHA-immunoprofiled cells were photographed by a Nikon Eclipse TI microscope, equipped with a 40X objective through the Hamamatsu 1394 ORCA-285 camera. (**Figures 3B** and **4B**) Collected as 1024×1024 (case Figure 3B) and 1344×1024 pixel images (case Figure 4B), photos were imported in Volocity 6.5.1 for analysis. Here, for each individual neuron an ROI was outlined by an operator blind of sample identity and background-subtracted, average αFlag and αHA, non-nuclear signals, as well as total-cell αPuro signal were collected.

#### PLA analysis

PLA-profiled cells were photographed by a Nikon C1 confocal system equipped with 40X-oil objective (**Figures 3A,5A,6,8** and **11**). Photos were collected as 2μm Z-stacks (step = 1μm) and 3μm Z-stacks (step = 1μm) of 1024×1024 pixel images, for **Figure 5A**, and **Figures 3A,6,8** and **11**, respectively. All primary images were generally analyzed with Volocity 6.5.1 software [here, positive spots were 3D-clusters including ≥1 voxels, each voxel corresponding to 0.1 μm^3^ and displaying a signal above 90 background standard deviations; for cumulative PLA signal calculation, only voxels above this threshold were taken into account]. Limited to **Figure 6A(b)**, files originating from flattened Z-stacks (max version) were imported into Adobe Photoshop CS6 and 2D-spots counting was performed manually, by an operator blind of sample identity. When appropriate (**Figures 3A** and **6A**), spot counting and/or cumulative signal evaluation was restricted to specific cell compartments (highlighted in grey, in idealized neuron silhouettes).

#### Common

Results of numerical image analysis were imported into Microsoft Excel, for subsequent processing. Finally, representative photos were edited for figures preparation, by ImageJ-Fiji and Adobe Photoshop CS6 softwares.

### Total RNA extraction

Total RNA was extracted from cells (**Figures 2D,E,4B**) using TRIzol Reagent (Thermofisher) according to the manufacturer’s instructions, with minor modifications. Briefly, for each biological replicate, a pellet including 300,000-800,000 cells was dissolved in 250-500μl of Trizol. RNA was precipitated using isopropanol and GlycoBlue (Ambion) overnight at −80°C. After two washes with 75% ethanol, the RNA was resuspended in 20μl sterile nuclease-free deionized water. Agarose gel electrophoresis and spectrophotometric measurements (NanoDrop ND-1000) were employed to estimate its concentration, quality and purity.

### Translating Ribosome Affinity Purification (TRAP) assay: RNA preparation

The TRAP assay was performed as previously described (Ainsley et al., 2014; Heiman et al., 2014) with minor modifications. For each TRAP reaction, 10μg of anti-GFP antibody, purchased from the Monoclonal Antibody Core Facility at the Memorial Sloan-Kettering Cancer Center (purified form of HtzGFP-19C8), were covalently bound to 1mg magnetic epoxy beads (Dynabeads Antibody Coupling kit, Life Technologies #14311D), according to manufacturer’s protocols, followed by BSA treatment to reduce non-specific binding. Antibody-coupled beads were resuspended at the concentration of 1mg/100μl. Cortico-cerebral cells, derived from *Rpl10a*^EGFP-Rpl10a/+^ embryos, were set up as described above (see “Primary cortico-cerebral cell cultures”) and as detailed in **Figure 2A**. At DIV8, cells were treated by supplementing medium with 0.1mg/ml cycloheximide (CHX; Sigma #C7698) at 37°C for 15 min. Then, cells were washed two times with ice-cold 1X PBS containing 0.1mg/ml CHX. 75μl ice-cold lysis buffer (see below) was added to each cells-containing well (12-multiwell plate) for 10 min on ice. Afterwards, cells were scraped and lysed by vigorously pipetting them up and down without creating bubbles. The lysate derived from two wells (about 1.6×10^6^ cells; corresponding to one biological replicate), was pooled. Upon addition to each replicate sample of 1/9 volume of 300mM 1,2-dihexanoyl-sn-glycero-3-phosphocholine (DHPC, Avanti Polar Lipids #850305), such sample was firstly centrifuged at 2000g for 10 min at 4°C. The supernatant was harvested and re-centrifuged, at 20000g for 10 min at 4°C. The resulting supernatant (about 150μl) was incubated with 100μl antibody-coupled beads for 1h at 4°C on a rotating wheel, at 10rpm. After incubation, beads were collected with a magnet: the immunoprecipitated component (TRAP-IP) bound to beads was washed four times with 1ml of ice-cold high-salt buffer (see below); the supernatant component (TRAP-SN) of each sample was stored on ice. [Lysis buffer: 20mM HEPES (Ambion), 150mM KCl (Ambion), 10mM MgCl_2_ (Ambion), 1%(vol/vol) NP-40 (Thermo Fisher Scientific), 1X EDTA-free protease inhibitors (Roche), 0.5mM DTT (Invitrogen), 0.1mg/ml cycloheximide, 10μl/ml rRNasin (Promega), 10μl/ml Superasin (Applied Biosystems). High-salt buffer: 20mM HEPES, 350mM KCl, 10mM MgCl_2_, 1%(vol/vol) NP-40, 1X EDTA-free protease inhibitors, 0.5mM DTT, 0.1mg/ml cycloheximide]. For each sample, RNA of TRAP-SN and TRAP-IP fractions were extracted with Trizol^®^ LS reagent (Thermofisher) according to manufacturer’s instructions, with minor modifications. The extraction procedure was repeated to improve RNA sample purity. RNA was finally precipitated using NaOAc, isopropanol and GlycoBlue overnight at −80°C, according to standard protocols. After two washes with 75% ethanol, the RNA was resuspended in 10μl sterile nuclease-free deionized water. Agarose gel electrophoresis and spectrophotometric measurements (NanoDrop ND-1000) were employed to estimate quantity, quality and purity of the resulting preparation.

### RNA immunoprecipitation (RIP) assay: RNA preparation

Cortico-cerebral cells were set up as described above (see “Primary cortico-cerebral cell cultures”) and as detailed in **Figure 7**. For each RIP reaction, 10μl of protein A/G Dynabeads (Thermofisher #492024) were coupled with 10μg of anti-protein of interest (POI; anti-Foxg1 ChIP-grade, rabbit polyclonal, Abcam #ab18259; anti-GFP, rabbit polyclonal, Abcam #ab290), or 10μg of rabbit IgG (Millipore #12370) as control, according to manufacturer’s protocols. Pre-clearing beads were prepared omitting antibody coupling. DIV8 cells were washed once with ice-cold 1X PBS. 75μl ice-cold lysis buffer (see below) was added to each cells-containg well (12-multiwell plate) for 10 min on ice. Afterwards, cells were scraped and lysed by vigorously pipetting them up and down without creating bubbles. The lysate derived from 10 wells (about 8×10^6^ cells; to be employed for one set of paired anti-POI/IgG assays), was pooled, pipetted up and down and kept 10 min on ice. Pipetting and incubation on ice were repeated. Next, each sample was centrifuged at 2000g for 10 min at 4°C. Then, the supernatant was re-centrifuged, at 16000g for 10 min at 4°C. The resulting supernatant was incubated with pre-clearing beads (pre-equilibrated in lysis buffer, see below) for 30 min at 4°C on a rotating wheel, at 10rpm. Then, the pre-clearing beads were removed with a magnet, and the supernatant was incubated with antibody-coupled beads (pre-equilibrated in lysis buffer), overnight at 4°C on a rotating wheel, at 10rpm. 10% of supernatant (Input, RIP-IN) was stored at −80°C. The day after, beads were collected with a magnet and the immunoprecipitated material bound to beads was harvested by washing them five times with 0.5ml of ice-cold high-salt buffer. [Lysis buffer: 25mM TRIS-HCl, 150mM KCl (Ambion), 10mM MgCl_2_ (Ambion), 1%(vol/vol) NP-40 (Thermo Fisher Scientific), 1X EDTA-free protease inhibitors (Roche), 0.5mM DTT (Invitrogen), 10ul/ml rRNasin (Promega), 10ul/ml Superasin (Applied Biosystems). High-salt buffer: 25mM TRIS-HCl, 350mM KCl (Ambion), 10mM MgCl_2_ (Ambion), 1%(vol/vol) NP-40 (Thermo Fisher Scientific), 1X EDTA-free protease inhibitors (Roche), 0.5mM DTT (Invitrogen)]. For each sample, immunoprecipitated RNA (RIP-IP) and Input (RIP-IN) were extracted with Trizol^®^ LS reagent according to manufacturer’s instructions, with minor modifications. The extraction procedure was repeated to improve RNA sample purity. RNA was precipitated using isopropanol and GlycoBlue overnight at −80°C, according to standard protocols. After two washes with 75% ethanol, the RNA was resuspended in 10ul sterile nuclease-free deionized water. Agarose gel electrophoresis and spectrophotometric measurements (NanoDrop ND-1000) were employed to estimate quantity, quality and purity of the resulting RNA.

### RNA quantitation: DNase treatment, Reverse Transcription and Real-Time quantitative PCR

DNA contaminants were removed from total-RNA, TRAP-SN, RIP-IN and RIP-IP samples by treating them with TURBO™ DNase (2U/μl) (Ambion) for 1h at 37°C, following manufacturer’s instructions. cDNA was produced via reverse transcription (RT) of the resulting preparations by SuperscriptIII™ (Invitrogen), primed by random hexamers, according to manufacturer’s instructions. For RT reactions, the following aliquots of RNA preparations were used: 1/10 TRAP-IP, 1/10 (DNA-free) TRAP-SN, 1/6 (DNA-free) IP- and IN-RIP, 0.5μg (DNA-free) total-RNA.

Following SuperscriptIII™ thermo-inactivation, the RT reaction (20 μl) was diluted 1:3 (in case of TRAP samples) or 1:5 (in case of RIP, and total-RNA samples), and 1-2μl of the resulting cDNA solution was used as substrate of any subsequent quantitative PCR (qPCR) reaction. Limited to intron-less amplicons and/or TRAP-IP, RIP-IN and RIP-IP samples, negative control PCRs were run on RT(-) RNA preparations. qPCR reactions were performed by the SsoAdvanced SYBR Green Supermix^TM^ platform (Biorad), according to manufacturer’s instructions, on a CFX BioRad thermocycler.

For each transcript under examination and each sample (i.e. biological replicate), cDNA was qPCR-analyzed in technical triplicate, and results averaged. In case of total-RNA and TRAP-IP and TRAP-SN samples, mRNA levels were normalized against *Rpl10a*-mRNA (Zhou et al., 2010). In addition, in case of TRAP samples, as indices of mRNA engagement to holoribosomes, IP/SN ratios were further calculated per each sample. In case of RIP samples, IP values were straightly normalized against IN values. Final results were averaged and the corresponding sem’s calculated, by Excel software.

The following oligonucleotides have been employed in this study:

*Psd95*/F: GCCGTGGCAGCCCTGAAGAACACA

*Psd95*/R: GCTGCTATGACTGATCTCATTGTCCAGG

*Foxg1(cds)*/F: GACAAGAAGAACGGCAAGTACGAGAAGC

*Foxg1(cds)*/R: GAACTCATAGATGCCATTGAGCGTCAGG

*Foxg1*(*5utr*)/F: TAGAAGCTGAAGAGGAGGTGGAGTGC

*Foxg1*(*5utr*)/R: CAGACCCAAACAGTCCCGAAATAAAGC

*Gria1*/F: TCCATGTGATCGAAATGAAGCATGATGGAATCC

*Gria1*/R: CGATGTAGGTTCTATTCTGGACGCTTGAGTTG

*pan-Grin1*/F: CGAGGATACCAGATGTCCACCAGACTAAAGA

*pan-Grin1*/R: CTTGACAGGGTCACCATTGACTGTGAACT

*ex20-Grin1*/F: CCGTGAACGTGTGGAGGAAGAACCT

*ex20-Grin1*/R: GTGTCTTTGGAGGACCTACGTCTCTTG

*Grid1*/F: AAGGACTGACTCTCAAAGTGGTGACTGTCTT

*Grid1*/R: CCTTAGCCAGTGCATCCAGCACATCTATG

*Gabra1*/F: AAACCAGTATGACCTTCTTGGACAAACAGTTGAC

*Gabra1*/R: GTGGAAGTGAGTCGTCATAACCACATATTCTC

*Slc17a6*/F: TTTTGCTGGAAAATCCCTCGGACAGATCTACA

*Slc17a6*/R: CTTACCGTCCTCTGTCAGCTCGATGG

*Bdnf2c*/F: CTTTGGGAAATGCAAGTGTTTATCACCAGGAT

*Bdnf4*/F: CTGCCTTGATGTTTACTTTGACAAGTAGTGACTG

*Bdnf(2c,4)*/R: GCCTTCATGCAACCGAAGTATGAAATAACCATAG

*Rpl10a*/F: CAGCAGCACTGTGATGAAGCCAAGG

*Rpl10a*/R: GGGATCTGCTTAATCAGAGACTCAGAGG

*F-rnoGrin1-H*/F: ACCTCCACCCTGGCCTCCAGCTT

*F-rnoGrin1-H*/R: GGGATAGCCAGCGTAATCTGGAACATC

*rnoGrin1.d*/F1: AGATCGCCCTCGACTTCGAAGAGC

*rnoGrin1.d*/F2: GTCGCACTCGCGCAACCCAGAG

*rnoGrin1.d*/F3: CCCAAGATCGTCAACATCGGCTGAGT

*rnoGrin1.d*/F4: CTGGCCGTGTGGAATTCAATGAGGATG

*rnoGrin1.d*/F5: TGCAGGATAGAAAGAGTGGTAGAGCAGA

*rnoGrin1.d*/F6: CAGTGGTGATGCCTAAAGGAATGTCAG

*rnoGrin1.d*/R1: CTCAGCACCGCCTCGAGTCCG

*rnoGrin1.d*/R2: ACTTCTGTGAAGCCTCAAACTCCAGCA

*rnoGrin1.d*/R3: TCCTCCCTCTCAATAGCGCGTCG

*rnoGrin1.d*/R4: TGTGGGTGACAGAAGTGGCGTTGAG

*rnoGrin1.d*/R5: GGGCAAACAACAGATGGCTGGCAACT

NB. rnoGrin1.d oligos employed for assays referred to by **Figure 7B** were associated as follows:

– d0/d1 assay: *rnoGrin1.d*/F1, *rnoGrin1.d*/F2*, rnoGrin1.d*/R1;
– d0/d2 assay: *rnoGrin1.d*/F3, *rnoGrin1.d*/R2, *rnoGrin1.d*/R4;
– d0/d3 assay: *rnoGrin1.d*/F4*, rnoGrin1.d*/F6, *rnoGrin1.d*/R5;
– d0/d4 assay: *rnoGrin1.d*/F5*, rnoGrin1.d*/F6*, rnoGrin1.d*/R5;
– d0/d5 assay: *rnoGrin1.d*/F3, *rnoGrin1.d*/R3, *rnoGrin1.d*/R4.

### TRAP-seq profiling

Produced as described in “Translating Ribosome Affinity Purification (TRAP) assay” section, TRAP-IP samples were sequenced by IGA Technology Services Srl. Libraries were produced using retrotranscribed cDNA previously amplified by Ovation Ultralow Library System V2 (NuGEN Technologies, Inc.). Library size and integrity were assessed using the Agilent Bioanalyzer (Santa Clara, CA) or Caliper GX (PerkinElmer, MA) apparatus. Sequencing was performed by Illumina HiSeq 2500 (Illumina, San Diego, CA). 20M paired-end reads (2×125nt) per biological replicate were generated (as elsewhere, biological replicates are independently cultured and engineered preparations, originating from a common cell pool); 3 *Foxg1*-OE and 4 Ctr replicate samples were profiled. Quality control of the sequenced reads was performed by a commercial operator (Sequentia, Barcelona, Spain) with the FASTQC v0.11.5 software, then low quality bases and adapters were removed with the software BBDuk version 35.85, setting a minimum base quality of 30 and a minimum read length of 35 bp. So-filtered high-quality reads were used in the following analyses.

### Ribosome engagement analysis

Transcripts whose ribosome engagement was affected by *Foxg1* overexpression were identified as follows. First, *Mus musculus* mRNA sequences (GRCm38.p6 reference genome version) were retrieved by Ensembl Biomart (Kinsella et al., 2011), selecting the principal isoform of each gene according to APPRIS annotations (Rodriguez et al., 2013) (if more transcripts were indexed at the highest level, then the longest one was selected). The resulting reference transcriptome included 22,442 transcripts. On this transcriptome, total RNA-seq FASTQ reads (Tigani et al., 2020) as well as TRAP-seq FASTQ reads, originating from sister primary neural cultures, were mapped using Bowtie2 (Langmead and Salzberg, 2012) (in “very-sensitive-local” configuration). Finally, the number of reads mapped to each transcript was computed by means of featureCounts (Liao et al., 2014) (with “primaryOnly=TRUE” and “minMQS=10” settings).

Next, for both total RNA-seq and TRAP-seq assays, differential gene expression analysis was performed using the R package DESeq2 software (Love et al., 2014). Then, for each gene, the difference between log_2_FC(trapRNAseq) and log_2_FC(totalRNAseq) (named Δlog_2_FC) was primarily calculated, as an index of *Foxg1*-OE impact on mRNA engagement to ribosomes. Moreover, statistical significance of Δlog_2_FC values was evaluated with Python package Ribodiff software (default parameters) (Zhong et al., 2017). Finally, genes were filtered out if not satisfying “p_adj_<0.1” conditions, as well as if not reaching the “baseMean” DESeq2 value of 200 in case of both RNA-seq and TRAP-seq profiling (Artimagnella and Mallamaci, doi: temporarily restricted).

### Ribosome progression analysis

This analysis was performed, taking advantage of the reference transcriptome generated for ribosome engagement analysis. TRAP-seq FASTQ reads were mapped on it using Bowtie2 (in “very-sensitive-local” configuration) and those falling within the cds’ further taken into account. Hence, for each transcript the cds was divided in 125-nt bins and the number of reads mapping to each bin was computed by featureCounts (with “allowMultiOverlap=TRUE”, “primaryOnly=TRUE”, and “minMQS=10” settings). Transcripts with <4 reads/bin in at least one sample out of 7 were filtered out. Then, for each bin/bin boundary the Ribosomal Progression Index (RPI) was calculated, as the ratio between the numbers of reads mapping downstream and upstream of it (to avoid potential infinites, numerator and denominator were increased by 1). The RPI of the last bin (3’ end) of all transcripts was discarded. Transcripts including a single cds-bin were not considered. Finally, for each bin of the 5040 transcripts analyzed which passed all the filters, the fold change (FC), i.e. the ratio among average RPI values peculiar to *Foxg1*-OE and *Ctr* groups, was calculated, and its statistical significance evaluated by t-test.

Next, for each transcript, boundaries with log_2_FC(RPI)≥1 and p<0.05 were annotated as “boundaries of interest, up” (boi_up_’s), and those with log_2_FC(rpi)≤-1 and p<0.05 as “boundaries of interest, down” (boi_down_’s). Then, boi_up_’s and boi_down_’s frequencies were evaluated over the full cds (f_boi.up_ and f_boi.down_, respectively). Finally, 34, (4+3)-type permutations of samples-set were built and the above analysis was performed for each of them in order to filter out potential false positive gene. Therefore, for each gene, f_boi.up_ and the f_boi.down_ z-scores were calculated and genes with z-scores<3 were filtered out (Artimagnella and Mallamaci, doi: temporarily restricted).

### Gene Ontology analysis

Gene Ontology analysis (GO) was performed with R package gProfiler2 software (Kolberg et al., 2020) (with “exclude_iea=T, user_threshold=0.1, sources=GO and correction_method= fdr” settings).

In the case of “Ribosome engagement analysis” genes, the input was the set of 358 “differentially engaged transcripts” (with p_adj_<0.1), while the background (custom_bg argument) included the 5122 transcripts with DESeq2 “baseMean”≥ 200 (referring to both RNA-seq and TRAP-seq data). FDR was set at 0.1.

In case of “Ribosome progression analysis” genes, the input was the set of 328 genes with f_boi.up_-z-score≥3 or f_boi.down_-z-score≥3, while the background (custom_bg argument) included the 5040 genes which passed all the “counting” filters listed above (Artimagnella and Mallamaci, doi: temporarily restricted). FDR was set at 0.2.

### Splicing and polyadenylation analyses

Both analyses were executed on totRNA samples from *Foxg1*-OE and control neocortical cultures (Artimagnella and Mallamaci, 2019), by a commercial operator (Sequentia, Barcelona, Spain).

In the case of splicing analysis, CASH software (Wu et al., 2018) was used. Genes with −0.1≥1′psi≥0.1 and fdr<0.05 were considered significant [here, 1′psi is the difference in percentage of “spliced-in transcripts” between *Foxg1*-OE and control samples].

In the case of polyadenylation analysis, ROAR software (Grassi et al., 2016) was used. Genes with 1/1.2≥r≥1.2 and padj<0.05 were considered significant [here, being the m/M the ratio between the shortest and the longest polyA isoform, r is the ratio between *Foxg1*-OE and control m/M parameters].

### RIP-seq profiling

Produced as described in “RNA immunoprecipitation (RIP) assay: RNA preparation” section, RIP samples were sequenced by IGA Technology Services Srl. Libraries were produced using retrotranscribed cDNA previously amplified by Ovation Ultralow Library System V2 (NuGEN Technologies, Inc.). Library size and integrity were assessed using the Agilent Bioanalyzer (Santa Clara, CA) or Caliper GX (PerkinElmer, MA) apparatus. Sequencing was performed by Illumina HiSeq 2500 (Illumina, San Diego, CA). 10M paired-end reads (2×125nt) per replicate were generated; 3 anti-Foxg1 and 3 IgG-Ctr paired samples were profiled. Quality control of the sequenced reads were performed by a commercial operator (Sequentia, Barcelona, Spain). Reads were processed with the FASTQC v0.11.5 software, then low quality bases and adapters were removed with the software BBDuk version 35.85, setting a minimum base quality of 30 and a minimum read length of 35 bp. So-filtered high-quality reads were used in the following analyses.

### RIP-seq analysis and identification of Foxg1-protein-interacting transcripts with Foxg1-sensitive ribosomal engagement and progression rates

Foxg1 protein-bound transcripts were identified as follows (steps 1-3 executed by Sequentia, Barcelona, Spain) First, a reference transcriptome was generated. The web-based tool Biomart (Kinsella et al., 2011) was used to extract GeneIDs, TranscriptIDs, cDNA sequences, APPRIS annotations (Rodriguez et al., 2013) and Transcript support levels (TSLs) of mouse genome GRCm38.p6. To select a unique representative transcript per gene, these rules were sequentially implemented: (1) the transcript with the highest APPRIS annotation level was chosen; (2) if multiple transcripts with the same annotation level were available, the transcript with the highest TSL was chosen; (3) if more than one transcript had the same TSL, one of them was randomly selected; (4) if a gene had no transcripts with either an APPRIS annotation or a TSL, one transcript was also randomly chosen. The final reference transcriptome consisted of 55647 unique transcripts.

Second, prior to mapping the reads to transcripts, the reference transcriptome was indexed with STAR (version 2.7.9a), using the genomeGenerate function. The parameter “genomeChrBinNbits” was set according to the formula:

Min(18, log2(max(GenomeLength/AmountOfReferences, ReadLength)))

RIP-seq reads were mapped in local alignment mode, with maximum intron size set to 1, so that the resulting BAM files did not actually include reads mapped on introns.

Third, Foxg1 protein/mRNA interaction peaks were identified by SICER2 (version 1.0.2) (Xu et al., 2014). The SICER2/sicer/lib/GenomeData.py file was manually edited in the SICER2 repository, to include a list of our reference transcriptome transcripts IDs, and a Python dictionary that maps these IDs to their lengths. Moreover, SICER was ran setting its parameters as follows: “fragment_size = median read length”, “redundancy_threshold = 1”, “window_size = 200”, “gap_size = 200”, and “effective_genome_fraction = 1”. In this way, 8352, 8851 and 7120 peak islands with fdr<0.1 were identified in samples 1, 2 and 3 respectively.

Fourth, peak islands were filtered out if not satisfying “aFoxg1/IgG_enrichment ≥2” and “fdr <0.05”. Next, transcripts sharing ≥1 peak island in ≥2 out of 3 biological replicates were considered as interacting with the Foxg1 protein. A total of 2857 transcripts satisfied this requirement.

Fifth, to estimate the magnitude of the geneset undergoing direct Foxg1 regulation of translation, these 2857 transcripts were intersected with the 358 and 328 ones resulting from our “Ribosome engagement analysis” and “Ribosome progression analysis” pipelines, respectively.

### Co-Immunoprecipitation (co-IP) assay

HEK293T cell lines were cultured and transfected as described in “HEK293T cell cultures” section, and as detailed in **Figure 5B**. After three days, cells were washed in 1X PBS and lysed with 500μl of CHAPS buffer, supplemented with 1X protease inhibitors (Roche). Next, lysates were processed for co-IP analysis by the FLAG Immunoprecipitation Kit (Sigma), according to manufacturer’s instructions. Specifically, total cell lysates were centrifuged at 12,000g for 10 min at 4°C, to remove debris. For each sample, the 4% of supernatant was saved as Input (IN). The remaining part was incubated with anti-Flag-conjugated resin for 3h at 4 °C, on a rotating wheel. Next, the immuno-precipitated resin (IP) was resuspended and washed 4 times in 1X wash buffer. Finally, IP and IN samples were denatured at 95 °C for 5 min in 1X sample buffer (supplemented with 0.5% β-Mercaptoethanol), prior to subsequent western blot analysis.

### Protein degradation assay

Cortico-cerebral cells were set up as described above (see “Primary cortico-cerebral cell cultures”) and as detailed in **Figure 3C**. At DIV8, cells were treated with 50μg/ml cycloheximide (CHX). Cells were analyzed at four different time points, 0, 6, 10, and 14 hours after CHX administration. For each point, samples were lysed in CHAPS buffer, supplemented with 1X protease inhibitors (Roche), and stored at −80°C. Upon thawing, samples were centrifuged at 12,000g for 10 min at 4°C, to remove debris, and then processed for western blot analysis.

### Western blot analysis

Western blot analysis was performed according to standard methods. Total cell lysates in CHAPS buffer were quantified by BCA protein assay kit (Fisher Scientific #10678484) (except for co-IP samples), and denatured at 95°C for 5 min, prior to loading. 20-30μg of proteins were loaded per each lane on a 10% acrylamide - 0.1% SDS gel. Afterwards, proteins were transferred to nitrocellulose membrane. Membranes were incubated 1h in 1X TBS-Tween containing 5% non-fat dry milk, before to be exposed to primary antibodies at 4°C overnight. Then, membranes were washed 3 times in 1X TBS-Tween, incubated 1h with HRP-conjugated secondary antibodies (DAKO, 1:2000) in 1X TBS-Tween containing 5% non-fat dry milk, at room temperature, washed again 3 times, and finally revealed by an ECL kit (GE Healthcare #GERPN2109). The following primary antibodies were used: anti-Foxg1, rabbit polyclonal (gift from G.Corte), 1:2000); anti-Flag, mouse monoclonal, clone M2 (Sigma #F1804), 1:1000; anti-Grin1-COOH-term, rabbit monoclonal, clone EPR2481(2) (Abcam #ab109182), 1:5000; anti-β-Actin HRP-conjugated, mouse monoclonal (Sigma #A3854), 1:20000. Images were acquired by an Alliance LD2–77.WL apparatus (Uvitec, Cambridge), and analyzed by Uvitec NineAlliance software. Finally, protein levels were normalized against Δ-actin.

### Numerical and statistical analysis

Full details of numerical and statistical analysis of data (including normalization criteria, number and definition of biological replicates, statistical tests employed for result evaluation) are provided in Figures and their legends.

Full primary data referred to in Figures and Supplementary Figures are reported in Supplementary Table S4.

## Supporting information

Supplementary Figures 1-2, with legends; Supplementary Tables S1-S4

## ACKNOWLEDGMENTS

We thank Simone Mortal for helping us with hippocampal cell culture and live-imaging experiments (**Figure 1G,H**).

We thank Cristina Fimiani for cloning “LVrc_TREt-pl-BGHpA”, employed as the backbone of some lentiviruses used in this study, as well as Vittoria Avaro for helping with immunofluorescence controls.

We thank Sequentia (Barcelona, Spain) for TRAPSeq service.

