## Supplementary Figures 1-2, with legends; Supplementary Tables S1-S4 for "Foxg1 regulates translation of neocortical neuronal genes, including the main NMDA receptor subunit gene, *Grin1*"

*by: Osvaldo Artimagnella et al*

**SUPPLEMENTARY INFORMATION**

*including:*

**- Supplementary Figures S1 and S2, with legends**

**- Tables S1, S2, S3 and S4 (the last reporting full primary data)**

**Figure S1**

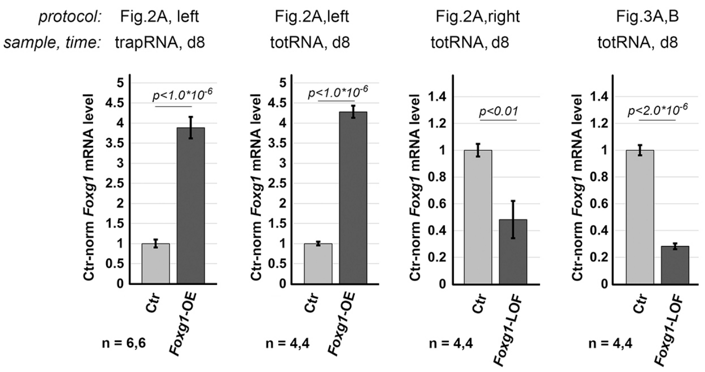

**Legend to Supplementary Figure S1. Evaluation of cumulative *Foxg1*-mRNA levels upon lentiviral delivery of a *Foxg1*-encoding or an *αFoxg1*-shRNA-encoding transgene to primary neocortical cultures.** Data refer to samples described as in Figures 2A and 3A,B. Normalization against *Rpl10a*. Throughout the Figure, *n* is the number of biological replicates, i.e. independently cultured and engineered preparations, originating from a common cell pool. Statistical evaluation of results performed by one-way ANOVA, one-tailed and unpaired. Errors bars indicate s.e.m.

**Figure S2**

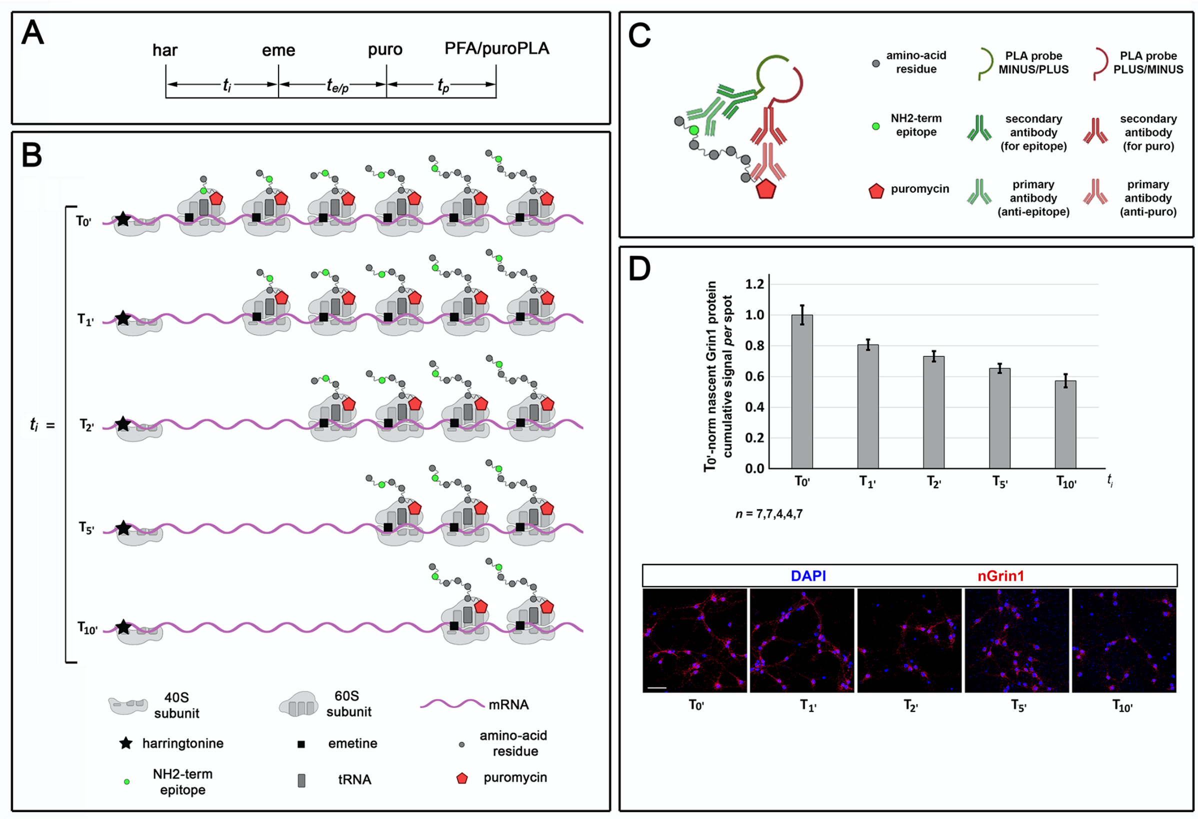

**Legend to Supplementary Figure S2. Puro-PLA run off assay.** (**A**) Temporal articulation of the puro-PLA run off assay. **(B)** Idealized representation of ribosomes progression along mRNA, upon harringtonine blockade of translation initiation. **(C**) Revealing puro-labeled polypeptides by PLA. (B) and (C) pictures created in “BioRender.com”. (**D**) Graph representing the cumulative signal per spot of nascent Grin1 (nGrin1) polypeptide at different time points after harringtonine administration, with examples of primary data. *n* is the number of images analyzed, evenly taken from pairs of independently cultured preparations, in turn originating from a common cell pool. Errors bars indicate s.e.m.

### Table S1. Translation factor genes in Foxg1-GOF neurons

*based on:*
*Artimagnella, Osvaldo, & Mallamaci, Antonello. (2020). Zenodo.*
*RNASeq profiling of Foxg1-GOF neocortical neurons.v2 (Version v2) [Data set].*
*https://doi.org/10.5281/zenodo.3739467*

| *Locus* | *log2FC* | *logCPM* | *P-value* | *FDR* | *Gene Name* |
|---|---|---|---|---|---|
| ENSMUSG00000035530 | -0.46 | 8.58 | 0.01 | 0.02 | Eif1 |
| ENSMUSG00000057561 | 0.15 | 6.06 | 0.07 | 0.12 | Eif1a |
| ENSMUSG00000024841 | -0.06 | 4.64 | 0.62 | 0.72 | Eif1ad |
| ENSMUSG00000067194 | 0.30 | 7.98 | 0.00 | 0.00 | Eif1ax |
| ENSMUSG00000006941 | -0.22 | 6.38 | 0.16 | 0.25 | Eif1b |
| ENSMUSG00000027810 | 0.17 | 5.44 | 0.05 | 0.09 | Eif2a |
| ENSMUSG00000029613 | -0.26 | 6.43 | 0.00 | 0.00 | Eif2ak1 |
| ENSMUSG00000031668 | 0.11 | 5.12 | 0.27 | 0.37 | Eif2ak3 |
| ENSMUSG00000005102 | -0.17 | 5.15 | 0.06 | 0.11 | Eif2ak4 |
| ENSMUSG00000029388 | -0.12 | 5.33 | 0.25 | 0.35 | Eif2b1 |
| ENSMUSG00000004788 | -0.12 | 4.77 | 0.42 | 0.54 | Eif2b2 |
| ENSMUSG00000028683 | -0.05 | 4.41 | 0.67 | 0.76 | Eif2b3 |
| ENSMUSG00000029145 | 0.01 | 5.39 | 0.95 | 0.97 | Eif2b4 |
| ENSMUSG00000003235 | -0.23 | 6.63 | 0.02 | 0.05 | Eif2b5 |
| ENSMUSG00000026427 | -0.25 | 5.19 | 0.00 | 0.01 | Eif2d |
| ENSMUSG00000021116 | 0.05 | 6.12 | 0.50 | 0.61 | Eif2s1 |
| ENSMUSG00000074656 | -0.49 | 7.26 | 0.00 | 0.00 | Eif2s2 |
| ENSMUSG00000035150 | 0.16 | 6.21 | 0.36 | 0.47 | Eif2s3x |
| ENSMUSG00000069049 | -0.89 | 5.39 | 0.00 | 0.00 | Eif2s3y |
| ENSMUSG00000024991 | 0.10 | 8.34 | 0.29 | 0.40 | Eif3a |
| ENSMUSG00000056076 | -0.09 | 7.31 | 0.44 | 0.56 | Eif3b |
| ENSMUSG00000030738 | -0.35 | 8.41 | 0.00 | 0.00 | Eif3c |
| ENSMUSG00000016554 | -0.56 | 6.91 | 0.00 | 0.00 | Eif3d |
| ENSMUSG00000022336 | -0.01 | 6.89 | 0.93 | 0.95 | Eif3e |
| ENSMUSG00000031029 | -0.13 | 7.15 | 0.47 | 0.58 | Eif3f |
| ENSMUSG00000070319 | -0.18 | 5.84 | 0.19 | 0.28 | Eif3g |
| ENSMUSG00000022312 | -0.07 | 7.14 | 0.62 | 0.71 | Eif3h |
| ENSMUSG00000028798 | -0.24 | 6.63 | 0.04 | 0.08 | Eif3i |
| ENSMUSG00000027236 | 0.12 | 4.40 | 0.23 | 0.33 | Eif3j1 |
| ENSMUSG00000043424 | -0.10 | 4.26 | 0.47 | 0.58 | Eif3j2 |
| ENSMUSG00000053565 | 0.02 | 6.22 | 0.92 | 0.95 | Eif3k |
| ENSMUSG00000033047 | 0.06 | 6.77 | 0.43 | 0.55 | Eif3l |
| ENSMUSG00000027170 | 0.01 | 6.47 | 0.93 | 0.95 | Eif3m |
| ENSMUSG00000059796 | -0.23 | 9.21 | 0.02 | 0.05 | Eif4a1 |
| ENSMUSG00000022884 | 0.35 | 9.54 | 0.00 | 0.00 | Eif4a2 |

| | | | | | |
|---|---|---|---|---|---|
| ENSMUSG00000025580 | -0.03 | 5.86 | 0.83 | 0.88 | Eif4a3 |
| ENSMUSG00000058655 | 0.28 | 8.23 | 0.00 | 0.00 | Eif4b |
| ENSMUSG00000028156 | -0.09 | 7.00 | 0.24 | 0.35 | Eif4e |
| ENSMUSG00000026254 | 0.12 | 6.62 | 0.13 | 0.20 | Eif4e2 |
| ENSMUSG00000093661 | 0.10 | 3.52 | 0.48 | 0.59 | Eif4e3 |
| ENSMUSG00000031490 | -2.16 | 3.15 | 0.00 | 0.00 | Eif4ebp1 |
| ENSMUSG00000020091 | -0.08 | 6.91 | 0.29 | 0.39 | Eif4ebp2 |
| ENSMUSG00000020454 | -0.02 | 6.08 | 0.85 | 0.90 | Eif4enif1 |
| ENSMUSG00000045983 | 0.08 | 8.77 | 0.38 | 0.49 | Eif4g1 |
| ENSMUSG00000005610 | 0.22 | 10.58 | 0.01 | 0.03 | Eif4g2 |
| ENSMUSG00000028760 | -0.22 | 8.12 | 0.08 | 0.14 | Eif4g3 |
| ENSMUSG00000040731 | 0.19 | 8.45 | 0.03 | 0.06 | Eif4h |
| ENSMUSG00000021282 | 0.03 | 7.98 | 0.74 | 0.81 | Eif5 |
| ENSMUSG00000078812 | -0.11 | 8.37 | 0.46 | 0.57 | Eif5a |
| ENSMUSG00000050192 | 0.48 | 4.60 | 0.01 | 0.01 | Eif5a2 |
| ENSMUSG00000026083 | -0.05 | 6.50 | 0.63 | 0.72 | Eif5b |
| ENSMUSG00000027613 | -0.01 | 5.38 | 0.93 | 0.95 | Eif6 |
| ENSMUSG00000037742 | 0.07 | 12.24 | 0.37 | 0.49 | Eef1a1 |
| ENSMUSG00000016349 | 1.08 | 9.06 | 0.00 | 0.00 | Eef1a2 |
| ENSMUSG00000025967 | 0.46 | 7.98 | 0.00 | 0.01 | Eef1b2 |
| ENSMUSG00000055762 | 0.01 | 6.21 | 0.97 | 0.98 | Eef1d |
| ENSMUSG00000001707 | -0.08 | 5.67 | 0.38 | 0.50 | Eef1e1 |
| ENSMUSG00000071644 | 0.16 | 8.29 | 0.25 | 0.35 | Eef1g |
| ENSMUSG00000034994 | 0.21 | 10.76 | 0.01 | 0.03 | Eef2 |
| ENSMUSG00000035064 | 0.19 | 6.25 | 0.03 | 0.05 | Eef2k |
| ENSMUSG00000033216 | 0.01 | 3.93 | 0.93 | 0.95 | Eefsec |
| ENSMUSG00000022283 | 0.35 | 8.38 | 0.00 | 0.00 | Pabpc1 |
| ENSMUSG00000011257 | 0.40 | 5.99 | 0.00 | 0.00 | Pabpc4 |
| ENSMUSG00000034732 | 0.87 | 1.94 | 0.28 | 0.38 | Pabpc5 |
| ENSMUSG00000022194 | 0.00 | 4.95 | 0.98 | 0.99 | Pabpn1 |
| ENSMUSG00000031960 | -0.55 | 8.62 | 0.00 | 0.00 | Aars |
| ENSMUSG00000023938 | 0.06 | 4.16 | 0.72 | 0.80 | Aars2 |
| ENSMUSG00000010755 | -0.79 | 7.36 | 0.00 | 0.00 | Cars |
| ENSMUSG00000056228 | -0.22 | 4.72 | 0.02 | 0.04 | Cars2 |
| ENSMUSG00000026356 | 0.20 | 6.16 | 0.01 | 0.03 | Dars |
| ENSMUSG00000026709 | -0.22 | 4.59 | 0.01 | 0.02 | Dars2 |
| ENSMUSG00000021420 | 0.04 | 3.38 | 0.76 | 0.83 | Fars2 |
| ENSMUSG00000003808 | -0.24 | 6.12 | 0.05 | 0.09 | Farsa |
| ENSMUSG00000026245 | -0.19 | 6.60 | 0.03 | 0.07 | Farsb |
| ENSMUSG00000029777 | -0.62 | 8.29 | 0.00 | 0.00 | Gars |

| | | | | | |
|---|---|---|---|---|---|
| ENSMUSG00000001380 | -0.36 | 7.07 | 0.00 | 0.00 | Hars |
| ENSMUSG00000019143 | -0.17 | 5.21 | 0.19 | 0.28 | Hars2 |
| ENSMUSG00000037851 | -0.67 | 7.37 | 0.00 | 0.00 | Iars |
| ENSMUSG00000026618 | -0.37 | 5.80 | 0.00 | 0.00 | Iars2 |
| ENSMUSG00000031948 | -0.03 | 6.69 | 0.82 | 0.88 | Kars |
| ENSMUSG00000024493 | -0.71 | 7.52 | 0.00 | 0.00 | Lars |
| ENSMUSG00000035202 | 0.57 | 9.09 | 0.03 | 0.06 | Lars2 |
| ENSMUSG00000007029 | -0.29 | 6.24 | 0.00 | 0.00 | Vars |
| ENSMUSG00000038838 | -0.06 | 4.64 | 0.51 | 0.62 | Vars2 |
| ENSMUSG00000028029 | -0.17 | 5.45 | 0.30 | 0.40 | Aimp1 |
| ENSMUSG00000029610 | -0.25 | 5.22 | 0.06 | 0.11 | Aimp2 |
| ENSMUSG00000028580 | 0.01 | 6.96 | 0.94 | 0.96 | Pum1 |
| ENSMUSG00000020594 | 0.27 | 7.75 | 0.08 | 0.14 | Pum2 |
| ENSMUSG00000041360 | -0.34 | 6.55 | 0.00 | 0.00 | Pum3 |
| ENSMUSG00000037331 | 0.01 | 7.35 | 0.94 | 0.96 | Larp1 |
| ENSMUSG00000025762 | -0.87 | 4.13 | 0.00 | 0.00 | Larp1b |
| ENSMUSG00000023025 | 0.37 | 5.96 | 0.05 | 0.09 | Larp4 |
| ENSMUSG00000033499 | -0.04 | 6.80 | 0.71 | 0.79 | Larp4b |
| ENSMUSG00000034839 | 0.63 | 4.97 | 0.00 | 0.00 | Larp6 |
| ENSMUSG00000027968 | 0.05 | 4.36 | 0.68 | 0.76 | Larp7 |
| ENSMUSG00000042426 | -0.02 | 5.42 | 0.81 | 0.87 | Dhx29 |

### Table S2A. Ribosomal-engagement: up-genes and down-genes (p-adj<0.1; see Figure 9): GO analysis (fdr<0.1)

***biological process***

| *fdr* | *term_ size* | *query_ size* | *inter-section _size* | *preci-sion* | *recall* | *term_id* | *term_name* |
|---|---|---|---|---|---|---|---|
| 0.009 | 336 | 358 | 46 | 0.128 | 0.137 | GO:0098916 | anterograde trans-synaptic signaling |
| 0.009 | 336 | 358 | 46 | 0.128 | 0.137 | GO:0007268 | chemical synaptic transmission |
| 0.009 | 352 | 358 | 47 | 0.131 | 0.134 | GO:0099536 | synaptic signaling |
| 0.009 | 341 | 358 | 46 | 0.128 | 0.135 | GO:0099537 | trans-synaptic signaling |
| 0.017 | 280 | 358 | 39 | 0.109 | 0.139 | GO:0007610 | behavior |
| 0.017 | 254 | 358 | 36 | 0.101 | 0.142 | GO:0099177 | regulation of trans-synaptic signaling |
| 0.017 | 253 | 358 | 36 | 0.101 | 0.142 | GO:0050804 | modulation of chemical synaptic transmission |
| 0.017 | 73 | 358 | 16 | 0.045 | 0.219 | GO:0007613 | memory |
| 0.038 | 8 | 358 | 5 | 0.014 | 0.625 | GO:0070542 | response to fatty acid |
| 0.083 | 512 | 358 | 57 | 0.159 | 0.111 | GO:0003008 | system process |
| 0.086 | 346 | 358 | 42 | 0.117 | 0.121 | GO:0050877 | nervous system process |
| 0.086 | 198 | 358 | 28 | 0.078 | 0.141 | GO:0042391 | regulation of membrane potential |

***cell compartment***

| *fdr* | *term _size* | *query_ size* | *inter-section _size* | *preci-sion* | *recall* | *term_id* | *term_name* |
|---|---|---|---|---|---|---|---|
| 0.000 | 340 | 358 | 49 | 0.137 | 0.144 | GO:0031226 | intrinsic component of plasma membrane |
| 0.000 | 607 | 358 | 73 | 0.204 | 0.120 | GO:0016021 | integral component of membrane |
| 0.000 | 649 | 358 | 76 | 0.212 | 0.117 | GO:0031224 | intrinsic component of membrane |
| 0.000 | 319 | 358 | 45 | 0.126 | 0.141 | GO:0005887 | integral component of plasma membrane |

| | | | | | | | |
|---|---|---|---|---|---|---|---|
| 0.009 | 374 | 358 | 46 | 0.128 | 0.123 | GO:0097447 | dendritic tree |
| 0.009 | 372 | 358 | 46 | 0.128 | 0.124 | GO:0030425 | dendrite |
| 0.011 | 414 | 358 | 49 | 0.137 | 0.118 | GO:0098794 | postsynapse |
| 0.012 | 83 | 358 | 16 | 0.045 | 0.193 | GO:0043235 | receptor complex |
| 0.035 | 1259 | 358 | 115 | 0.321 | 0.091 | GO:0071944 | cell periphery |
| 0.038 | 489 | 358 | 53 | 0.148 | 0.108 | GO:0036477 | somatodendritic compartment |
| 0.038 | 1176 | 358 | 108 | 0.302 | 0.092 | GO:0005886 | plasma membrane |
| 0.041 | 300 | 358 | 36 | 0.101 | 0.120 | GO:0098978 | glutamatergic synapse |
| 0.041 | 12 | 358 | 5 | 0.014 | 0.417 | GO:1902710 | GABA receptor complex |
| 0.047 | 89 | 358 | 15 | 0.042 | 0.169 | GO:0034702 | ion channel complex |
| 0.047 | 844 | 358 | 81 | 0.226 | 0.096 | GO:0030054 | cell junction |
| 0.052 | 727 | 358 | 71 | 0.198 | 0.098 | GO:0045202 | synapse |
| 0.052 | 119 | 358 | 18 | 0.050 | 0.151 | GO:0043197 | dendritic spine |
| 0.052 | 664 | 358 | 66 | 0.184 | 0.099 | GO:0043005 | neuron projection |
| 0.059 | 66 | 358 | 12 | 0.034 | 0.182 | GO:0098982 | GABA-ergic synapse |
| 0.061 | 123 | 358 | 18 | 0.050 | 0.146 | GO:0044309 | neuron spine |
| 0.061 | 27 | 358 | 7 | 0.020 | 0.259 | GO:0032590 | dendrite membrane |
| 0.066 | 43 | 358 | 9 | 0.025 | 0.209 | GO:0043198 | dendritic shaft |
| 0.066 | 35 | 358 | 8 | 0.022 | 0.229 | GO:0034705 | potassium channel complex |
| 0.067 | 156 | 358 | 21 | 0.059 | 0.135 | GO:0098797 | plasma membrane protein complex |
| 0.068 | 15 | 358 | 5 | 0.014 | 0.333 | GO:0016529 | sarcoplasmic reticulum |

| | | | | | | | |
|---|---|---|---|---|---|---|---|
| 0.071 | 79 | 358 | 13 | 0.036 | 0.165 | GO:0034703 | cation channel complex |
| 0.079 | 37 | 358 | 8 | 0.022 | 0.216 | GO:0032589 | neuron projection membrane |
| 0.079 | 10 | 358 | 4 | 0.011 | 0.400 | GO:1902711 | GABA-A receptor complex |
| 0.087 | 38 | 358 | 8 | 0.022 | 0.211 | GO:0098802 | plasma membrane signaling receptor complex |
| 0.087 | 833 | 358 | 77 | 0.215 | 0.092 | GO:0042995 | cell projection |
| 0.087 | 2 | 358 | 2 | 0.006 | 1.000 | GO:0099544 | perisynaptic space |
| 0.087 | 824 | 358 | 76 | 0.212 | 0.092 | GO:0120025 | plasma membrane bounded cell projection |
| 0.087 | 153 | 358 | 20 | 0.056 | 0.131 | GO:1990351 | transporter complex |
| 0.087 | 298 | 358 | 33 | 0.092 | 0.111 | GO:0098793 | presynapse |
| 0.087 | 321 | 358 | 35 | 0.098 | 0.109 | GO:0043025 | neuronal cell body |
| 0.087 | 2 | 358 | 2 | 0.006 | 1.000 | GO:0032426 | stereocilium tip |
| 0.093 | 66 | 358 | 11 | 0.031 | 0.167 | GO:0031256 | leading edge membrane |
| 0.096 | 95 | 358 | 14 | 0.039 | 0.147 | GO:0044306 | neuron projection terminus |

***molecular function***

| *fdr* | *term _size* | *query_ size* | *inter-section _size* | *preci-sion* | *recall* | *term_id* | *term_name* |
|---|---|---|---|---|---|---|---|
| 0.005 | 34 | 358 | 11 | 0.031 | 0.324 | GO:0022834 | ligand-gated channel activity |
| 0.005 | 32 | 358 | 11 | 0.031 | 0.344 | GO:0015276 | ligand-gated ion channel activity |
| 0.019 | 22 | 358 | 8 | 0.022 | 0.364 | GO:0005230 | extracellular ligand-gated ion channel activity |
| 0.033 | 83 | 358 | 16 | 0.045 | 0.193 | GO:0022836 | gated channel activity |
| 0.045 | 21 | 358 | 7 | 0.020 | 0.333 | GO:0022835 | transmitter-gated channel activity |
| 0.045 | 21 | 358 | 7 | 0.020 | 0.333 | GO:1904315 | transmitter-gated ion channel activity involved in regulation of postsynaptic membrane potential |

| | | | | | | | |
|---|---|---|---|---|---|---|---|
| 0.045 | 21 | 358 | 7 | 0.020 | 0.333 | GO:0022824 | transmitter-gated ion channel activity |
| 0.054 | 22 | 358 | 7 | 0.020 | 0.318 | GO:0099529 | neurotransmitter receptor activity involved in regulation of postsynaptic membrane potential |
| 0.064 | 24 | 358 | 7 | 0.020 | 0.292 | GO:0015108 | chloride transmembrane transporter activity |
| 0.064 | 23 | 358 | 7 | 0.020 | 0.304 | GO:0098960 | postsynaptic neurotransmitter receptor activity |
| 0.064 | 209 | 358 | 27 | 0.075 | 0.129 | GO:0015075 | ion transmembrane transporter activity |
| 0.064 | 12 | 358 | 5 | 0.014 | 0.417 | GO:0099095 | ligand-gated anion channel activity |
| 0.064 | 117 | 358 | 18 | 0.050 | 0.154 | GO:0005216 | ion channel activity |
| 0.064 | 12 | 358 | 5 | 0.014 | 0.417 | GO:0016917 | GABA receptor activity |
| 0.064 | 25 | 358 | 7 | 0.020 | 0.280 | GO:0030594 | neurotransmitter receptor activity |
| 0.064 | 108 | 358 | 17 | 0.047 | 0.157 | GO:0046873 | metal ion transmembrane transporter activity |
| 0.072 | 163 | 358 | 22 | 0.061 | 0.135 | GO:0008324 | cation transmembrane transporter activity |
| 0.072 | 184 | 358 | 24 | 0.067 | 0.130 | GO:0015318 | inorganic molecular entity transmembrane transporter activity |
| 0.072 | 122 | 358 | 18 | 0.050 | 0.148 | GO:0022803 | passive transmembrane transporter activity |
| 0.072 | 41 | 358 | 9 | 0.025 | 0.220 | GO:0015081 | sodium ion transmembrane transporter activity |
| 0.072 | 122 | 358 | 18 | 0.050 | 0.148 | GO:0015267 | channel activity |
| 0.072 | 20 | 358 | 6 | 0.017 | 0.300 | GO:0099094 | ligand-gated cation channel activity |
| 0.072 | 27 | 358 | 7 | 0.020 | 0.259 | GO:0015171 | amino acid transmembrane transporter activity |
| 0.077 | 9 | 358 | 4 | 0.011 | 0.444 | GO:0005416 | amino acid:cation symporter activity |
| 0.095 | 100 | 358 | 15 | 0.042 | 0.150 | GO:0001216 | DNA-binding transcription activator activity |
| 0.095 | 22 | 358 | 6 | 0.017 | 0.273 | GO:0005267 | potassium channel activity |
| 0.095 | 10 | 358 | 4 | 0.011 | 0.400 | GO:0022851 | GABA-gated chloride ion channel activity |

| | | | | | | | |
|---|---|---|---|---|---|---|---|
| 0.095 | 150 | 358 | 20 | 0.056 | 0.133 | GO:0022890 | inorganic cation transmembrane transporter activity |
| 0.095 | 10 | 358 | 4 | 0.011 | 0.400 | GO:0004890 | GABA-A receptor activity |
| 0.095 | 100 | 358 | 15 | 0.042 | 0.150 | GO:0001228 | DNA-binding transcription activator activity, RNA polymerase II-specific |
| 0.095 | 5 | 358 | 3 | 0.008 | 0.600 | GO:0005047 | signal recognition particle binding |
| 0.095 | 5 | 358 | 3 | 0.008 | 0.600 | GO:0005313 | L-glutamate transmembrane transporter activity |

### Table S2B. Ribosomal-progression: genes with f(boi.up)-z-score≥3 or f(boi.down)-z-score≥3 (see Figure 10): GO analysis (fdr<0.2)

***biological process***

| *fdr* | *term_ size* | *query_ size* | *inter-section _size* | *preci-sion* | *recall* | *term_id* | *term_name* |
|---|---|---|---|---|---|---|---|
| --- | --- | --- | --- | --- | --- | --- | --- |

***cell compartment***

| *fdr* | *term_ size* | *query_ size* | *inter-section _size* | *preci-sion* | *recall* | *term_id* | *term_name* |
|---|---|---|---|---|---|---|---|
| --- | --- | --- | --- | --- | --- | --- | --- |

***molecular function***

| *fdr* | *term_ size* | *query_ size* | *interse ction_si ze* | *preci-sion* | *recall* | *term_id* | *term_name* |
|---|---|---|---|---|---|---|---|
| 0.169 | 8 | 327 | 4 | 0.012 | 0.500 | GO:0000062 | fatty-acyl-CoA binding |
| 0.169 | 180 | 327 | 23 | 0.070 | 0.128 | GO:0004672 | protein kinase activity |
| 0.169 | 146 | 327 | 21 | 0.064 | 0.144 | GO:0004674 | protein serine/threonine kinase activity |
| 0.169 | 160 | 327 | 21 | 0.064 | 0.131 | GO:0030554 | adenyl nucleotide binding |
| 0.169 | 155 | 327 | 21 | 0.064 | 0.135 | GO:0032559 | adenyl ribonucleotide binding |

| | | | | | | | |
|---|---|---|---|---|---|---|---|
| 0.169 | 8 | 327 | 4 | 0.012 | 0.500 | GO:0120227 | acyl-CoA binding |
| 0.175 | 9 | 327 | 4 | 0.012 | 0.444 | GO:1901567 | fatty acid derivative binding |
| 0.198 | 5 | 327 | 3 | 0.009 | 0.600 | GO:0031402 | sodium ion binding |
| 0.198 | 5 | 327 | 3 | 0.009 | 0.600 | GO:0048185 | activin binding |

### Table S3. Plasmids and DNA fragments employed in this study: selected sequences.

**- “SrfI_Foxg1(cds-3’term)-EGFP_ApaI” fragment**

> SrfI-cut_mmuFoxg1-cds (nt1282-to-endSTOP-less)

GGGCCGCGTCCTCCTCTACGTCGCCGCAGGCCCCCTCGACCCTGCCCTGTGAGTCTTTAAGACCCTCT
TTGCCAAGTTTTACGACAGGACTGTCCGGGGGACTGTCTGATTATTTCACACATCAAAATCAGGGGTC
TTCTTCCAACCCTTTAATACAT

>5xGS-repeats

CTCGAGGGCTCCGGAAGCGGGTCTGGAAGCGGCTCCCTCGAG

> EGFP-2xSTOP_ApaI-cut

ATGGTGAGCAAGGGCGAGGAGCTGTTCACCGGGGTGGTGCCCATCCTGGTCGAGCTGGACGGCGACGT
AAACGGCCACAAGTTCAGCGTGTCCGGCGAGGGCGAGGGCGATGCCACCTACGGCAAGCTGACCCTGA
AGTTCATCTGCACCACCGGCAAGCTGCCCGTGCCCTGGCCCACCCTCGTGACCACCCTGACCTACGGC
GTGCAGTGCTTCAGCCGCTACCCCGACCACATGAAGCAGCACGACTTCTTCAAGTCCGCCATGCCCGA
AGGCTACGTCCAGGAGCGCACCATCTTCTTCAAGGACGACGGCAACTACAAGACCCGCGCCGAGGTGA
AGTTCGAGGGCGACACCCTGGTGAACCGCATCGAGCTGAAGGGCATCGACTTCAAGGAGGACGGCAAC
ATCCTGGGGCACAAGCTGGAGTACAACTACAACAGCCACAACGTCTATATCATGGCCGACAAGCAGAA
GAACGGCATCAAGGTGAACTTCAAGATCCGCCACAACATCGAGGACGGCAGCGTGCAGCTCGCCGACC
ACTACCAGCAGAACACCCCCATCGGCGACGGCCCCGTGCTGCTGCCCGACAACCACTACCTGAGCACC
CAGTCCGCCCTGAGCAAAGACCCCAACGAGAAGCGCGATCACATGGTCCTGCTGGAGTTCGTGACCGC
CGCCGGGATCACTCTCGGCATGGACGAGCTGTACAAGTAAGTCGACGGATCCTAAGGGCC

**- “Flag-rnoGrin1-203-HA” fragment**

>SalI-cut_ rnoGrin1-203-5’utr

TCGACCGCGCACTCGACTCAGCGTCAGGAAGCGGGGGCGGTGGGAGGGGTAGAACGCGTAGGTCCCGC
TCATGACTCCGCAGCTGCTGCAGTCGCCGCAGCATCGGGACCAGTCGCGCAGTCCGCGCTGCTGTCCT
TTCCGCCTTTTCCGCGCGGGTGTTCGAGCAGCGCCAAACACGCTTCAGCACCTCGGACAGCATCCGCC
GCGCTCGCCCGGGGCTCCTAGAGAACCCGGGGGCGCTTGACCGCGCGCGGGCGGCCCGCGGGTCGTAC
ATCGCGAGGTCGTCGCACTCGCGCAACCCAGAGCCAGGCCCGCTGTGCCCGGAGCTC

> rnoGrin1-203-cds(signal-peptide)

ATGAGCACCATGCACCTGCTGACATTCGCCCTGCTTTTTTCCTGCTCCTTCGCCCGC

>3xFlag

GATTACAAGGACCACGACGGAGACTACAAAGATCATGATATCGATTATAAGGACGATGACGATAAG

> rnoGrin1-203-cds(aa20-end)

GCCGCCTGCGACCCCAAGATCGTCAACATCGGCGCGGTGCTGAGCACGCGCAAGCATGAACAGATGTT
CCGCGAGGCAGTAAACCAGGCCAATAAGCGACACGGCTCTTGGAAGATACAGCTCAACGCCACTTCTG
TCACCCACAAGCCCAACGCCATACAGATGGCCCTGTCAGTGTGTGAGGACCTCATCTCTAGCCAGGTC
TACGCTATCCTAGTTAGCCACCCGCCTACTCCCAACGACCACTTCACTCCCACCCCTGTCTCCTACAC
AGCTGGCTTCTACAGAATCCCTGTCCTGGGACTGACTACCCGAATGTCCATCTACTCTGACAAGAGTA
TCCACCTGAGTTTCCTTCGCACGGTGCCGCCCTACTCCCACCAGTCCAGCGTCTGGTTTGAGATGATG
CGAGTCTACAACTGGAACCACATCATCCTGCTGGTCAGCGACGACCACGAGGGACGGGCAGCGCAGAA

GCGCTTGGAGACGTTGCTGGAGGAACGGGAGTCCAAGGCAGAGAAGGTGCTGCAGTTTGACCCAGGAA
CCAAGAATGTGACGGCTCTGCTGATGGAGGCCCGGGAACTGGAGGCCCGGGTCATCATCCTTTCTGCA
AGCGAGGACGACGCTGCCACAGTGTACCGCGCAGCCGCAATGCTGAACATGACGGGCTCTGGGTACGT
GTGGCTGGTCGGGGAACGCGAGATCTCTGGGAACGCCCTGCGCTACGCTCCTGATGGCATCATCGGAC
TTCAGCTCATCAATGGCAAGAATGAGTCAGCCCACATCAGTGACGCCGTGGGCGTGGTGGCACAGGCA
GTTCACGAACTCCTAGAGAAGGAGAATATCACTGACCCACCGCGGGGTTGCGTGGGCAACACCAACAT
CTGGAAGACAGGACCATTGTTCAAGAGGGTGCTGATGTCTTCTAAGTATGCGGACGGAGTGACTGGCC
GTGTGGAATTCAATGAGGATGGGGACCGGAAGTTTGCCAACTATAGTATCATGAACCTGCAGAACCGC
AAGCTGGTGCAAGTGGGCATCTACAATGGTACCCATGTCATCCCAAATGACAGGAAGATCATCTGGCC
AGGAGGAGAGACAGAGAAACCTCGAGGATACCAGATGTCCACCAGACTAAAGATAGTGACAATCCACC
AAGAGCCCTTCGTGTACGTCAAGCCCACAATGAGTGATGGGACATGCAAAGAGGAGTTCACAGTCAAT
GGTGACCCAGTGAAGAAGGTGATCTGTACGGGGCCTAATGACACGTCCCCAGGCAGCCCACGCCACAC
AGTGCCCCAGTGCTGCTATGGCTTCTGCATAGACCTGCTCATCAAGCTGGCGCGGACCATGAATTTTA
CCTATGAGGTGCACCTGGTGGCAGATGGCAAGTTTGGCACACAGGAGCGGGTAAACAACAGCAACAAA
AAGGAGTGGAACGGAATGATGGGCGAGCTACTCAGTGGCCAAGCGGACATGATTGTGGCACCACTGAC
CATCAACAATGAGCGTGCGCAGTACATAGAGTTCTCCAAGCCCTTCAAGTACCAGGGCCTGACCATTT
TGGTCAAGAAGGAGATTCCCAGGAGCACACTGGACTCATTTATGCAGCCTTTTCAGAGCACACTGTGG
TTGCTAGTAGGACTGTCAGTTCATGTGGTGGCTGTGATGCTGTACCTGCTGGACCGCTTCAGTCCCTT
TGGCCGATTCAAGGTGAACAGTGAGGAGGAGGAGGAAGATGCACTGACCCTGTCCTCTGCCATGTGGT
TTTCCTGGGGCGTCCTGCTCAACTCCGGCATTGGGGAAGGTGCCCCCCGGAGTTTCTCTGCACGTATC
CTAGGCATGGTGTGGGCTGGTTTCGCCATGATCATAGTGGCTTCCTACACTGCCAACTTGGCAGCTTT
CCTGGTGCTGGATCGGCCTGAGGAGCGCATCACGGGCATCAATGACCCCAGGCTCAGAAACCCCTCAG
ACAAGTTCATCTACGCAACTGTAAAGCAGAGCTCCGTGGACATCTACTTCCGGAGGCAGGTGGAGTTG
AGTACCATGTACCGGCACATGGAAAAACACAATTACGAGAGCGCAGCTGAGGCCATCCAGGCTGTGCG
GGACAACAAGCTGCACGCCTTTATCTGGGACTCGGCCGTGCTGGAGTTTGAGGCTTCACAGAAGTGCG
ATCTGGTGACCACGGGTGAGCTGTTCTTCCGCTCAGGCTTTGGCATCGGCATGCGCAAGGACAGCCCC
TGGAAGCAGAACGTTTCCCTGTCCATACTCAAGTCCCATGAGAATGGCTTCATGGAAGATCTGGATAA
GACATGGGTTCGGTATCAGGAATGCGACTCCCGCAGCAATGCTCCTGCAACCCTCACTTTTGAGAACA
TGGCAGGGGTCTTCATGCTGGTGGCTGGAGGCATCGTAGCTGGGATTTTCCTCATTTTCATTGAGATC
GCCTACAAGCGACACAAGGATGCCCGTAGGAAGCAGATGCAGCTGGCTTTTGCAGCCGTGAACGTGTG
GAGGAAGAACCTGCAGGATAGAAAGAGTGGTAGAGCAGAGCCCGACCCTAAAAAGAAAGCCACATTTA
GGGCTATCACCTCCACCCTGGCCTCCAGCTTCAAGAGACGTAGGTCCTCCAAAGACACGAGCACCGGG
GGTGGACGCGGCGCTTTGCAAAACCAAAAAGACACAGTGCTGCCGCGACGCGCTATTGAGAGGGAGGA
GGGCCAGCTGCAGCTGTGTTCCCGTCATAGGGAGAGC

>3xHA-STOP
TACCCGTACGATGTTCCAGATTACGCTGGCTATCCCTATGACGTCCCTGACTACGCAGGATCCTATCC
TTACGACGTGCCGGACTATGCTTGA

> rnoGrin1-203-3’utr(3’D4nt)
GACGCCCCGCCCGCCCTCCTCTGCCCCTCCCCCGCAGACAGACGCACGGGACAGCGGCCTGGCCCACG
CAGAGCCCCGGAGCACGACGGGGTCGGGGGAGGAGCACTCCCAGCCTCCCCCAGGCCGTGCCCGCCTG
CCCACCGGTCGGCCGGCTGGCCGGTCCACCCTGTCCCGGCCCCGCGCGTGCCCCCGACGTCGGAGCTA
ACGGGCCGCCTTGTCTGTGTATTTCTATTTTACAGCAGTACCATCCCACTGATATCACGGGCCCGCTC
AACCTCTCAGATCCCTCGGTCAGCACCGTGGTGTGAGGCCCCCCGGAGGCGCCCACCTGCCCAGTTAG
CCCGGCCAAGGACACTGATGAGTCCTGCTGCTCGGGAAGGCCTGAGGGAAGCCCACCCGCCCCAGAGA
CTGCCCACCCTGGGCCTCCCGTCCGCCTGCTCTGCTGCCTGGCGGGCAGCCCCTGCAGGACCAAGGTG
CGGACCAGAGCGGCTGAGGATGGGCCAGAGCTGAGCCGGCTGGGCAGGGCCACAGGGCGCTCCGGCAG
AGGCAGGGCCCTGAGGTCTCTGAGCAGTGGGGTGAGGGGCCTAAGTGGCCCCGGTCGGAGGAGTCTGG
AGCAGAAATGGCAGCCCCATCCTTCCTCCAGCCACTACCCCAAGCTACAGTGGGGGCCTATGGCCCCA
GCTTGCTAGGTCACCCCCGACCCTTCCTCCAGCGCCTGCTCTCTGCAACTTGATTTCCACCTCTCTCC

TGCTGCACCACCCTCCCACGACATTTCCCCACCCCATTCACTGGGTTGTCTCTGACCTTTCCCAGGGC
TAGCCTTCACTGCCCTAGTGGCAGTGCTTCAGGGGTGCTTTCTGGCTCCCAGACATCTAGGGCTCCAG
ACTCCAAGAGGGCTGAGCCTTCTCTTCTGTCCGCAGCCACAATAGGCTTCCTCAGACGCTGGCTCGTG
ATGAGTCCCGCACCTTGGGCACCAGGGAGCGCCATCTGCCTCCCAGTCCGGTGTCACTCACCCCACTA
CCTTGTACATGACCAGCTCTCCCAGTGTCCCAGTGTCTGCCCCAGGGACACCGGGCGCGCACAGCCAC
CCCTAATCCCGGTATTCAGTGGTGATGCCTAAAGGAATGTCAGAAAA

>polyA-synth_XbaI-cut
AAAAAAAAAAAGCGGCCGCAATAAAAGATCTTTATTTTCATTAGATCTGTGTGTTGGTTTTTTGTGTG
T

**- “rnoGrin1-203*.full” fragment**

> SalI-cut_ polylinker(sx)-rnoGrin1-203-5’utr
TCGACTTCGAACGCGCACTCGACTCAGCGTCAGGAAGCGGGGGCGGTGGGAGGGGTAGAACGCGTAGG
TCCCGCTCATGACTCCGCAGCTGCTGCAGTCGCCGCAGCATCGGGACCAGTCGCGCAGTCCGCGCTGC
TGTCCTTTCCGCCTTTTCCGCGCGGGTGTTCGAGCAGCGCCAAACACGCTTCAGCACCTCGGACAGCA
TCCGCCGCGCTCGCCCGGGGCTCCTAGAGAACCCGGGGGCGCTTGACCGCGCGCGGGCGGCCCGCGGG
TCGTACATCGCGAGGTCGTCGCACTCGCGCAACCCAGAGCCAGGCCCGCTGTGCCCGGAGCTC

> rnoGrin1-203-cds(aa1-30)-STOP
ATGAGCACCATGCACCTGCTGACATTCGCCCTGCTTTTTTCCTGCTCCTTCGCCCGCGCCGCCTGCGA
CCCCAAGATCGTCAACATCGGCTGA

>polylinker-rnoGrin1-203-cds(aa31-end)
GTTTAAACTCCGGACTCGAGGCGGTGCTGAGCACGCGCAAGCATGAACAGATGTTCCGCGAGGCAGTA
AACCAGGCCAATAAGCGACACGGCTCTTGGAAGATACAGCTCAACGCCACTTCTGTCACCCACAAGCC
CAACGCCATACAGATGGCCCTGTCAGTGTGTGAGGACCTCATCTCTAGCCAGGTCTACGCTATCCTAG
TTAGCCACCCGCCTACTCCCAACGACCACTTCACTCCCACCCCTGTCTCCTACACAGCTGGCTTCTAC
AGAATCCCTGTCCTGGGACTGACTACCCGAATGTCCATCTACTCTGACAAGAGTATCCACCTGAGTTT
CCTTCGCACGGTGCCGCCCTACTCCCACCAGTCCAGCGTCTGGTTTGAGATGATGCGAGTCTACAACT
GGAACCACATCATCCTGCTGGTCAGCGACGACCACGAGGGACGGGCAGCGCAGAAGCGCTTGGAGACG
TTGCTGGAGGAACGGGAGTCCAAGGCAGAGAAGGTGCTGCAGTTTGACCCAGGAACCAAGAATGTGAC
GGCTCTGCTGATGGAGGCCCGGGAACTGGAGGCCCGGGTCATCATCCTTTCTGCAAGCGAGGACGACG
CTGCCACAGTGTACCGCGCAGCCGCAATGCTGAACATGACGGGCTCTGGGTACGTGTGGCTGGTCGGG
GAACGCGAGATCTCTGGGAACGCCCTGCGCTACGCTCCTGATGGCATCATCGGACTTCAGCTCATCAA
TGGCAAGAATGAGTCAGCCCACATCAGTGACGCCGTGGGCGTGGTGGCACAGGCAGTTCACGAACTCC
TAGAGAAGGAGAATATCACTGACCCACCGCGGGGTTGCGTGGGCAACACCAACATCTGGAAGACAGGA
CCATTGTTCAAGAGGGTGCTGATGTCTTCTAAGTATGCGGACGGAGTGACTGGCCGTGTGGAATTCAA
TGAGGATGGGGACCGGAAGTTTGCCAACTATAGTATCATGAACCTGCAGAACCGCAAGCTGGTGCAAG
TGGGCATCTACAATGGTACCCATGTCATCCCAAATGACAGGAAGATCATCTGGCCAGGAGGAGAGACA
GAGAAACCTCGAGGATACCAGATGTCCACCAGACTAAAGATAGTGACAATCCACCAAGAGCCCTTCGT
GTACGTCAAGCCCACAATGAGTGATGGGACATGCAAAGAGGAGTTCACAGTCAATGGTGACCCAGTGA
AGAAGGTGATCTGTACGGGGCCTAATGACACGTCCCCAGGCAGCCCACGCCACACAGTGCCCCAGTGC
TGCTATGGCTTCTGCATAGACCTGCTCATCAAGCTGGCGCGGACCATGAATTTTACCTATGAGGTGCA
CCTGGTGGCAGATGGCAAGTTTGGCACACAGGAGCGGGTAAACAACAGCAACAAAAAGGAGTGGAACG
GAATGATGGGCGAGCTACTCAGTGGCCAAGCGGACATGATTGTGGCACCACTGACCATCAACAATGAG
CGTGCGCAGTACATAGAGTTCTCCAAGCCCTTCAAGTACCAGGGCCTGACCATTTTGGTCAAGAAGGA
GATTCCCAGGAGCACACTGGACTCATTTATGCAGCCTTTTCAGAGCACACTGTGGTTGCTAGTAGGAC
TGTCAGTTCATGTGGTGGCTGTGATGCTGTACCTGCTGGACCGCTTCAGTCCCTTTGGCCGATTCAAG
GTGAACAGTGAGGAGGAGGAGGAAGATGCACTGACCCTGTCCTCTGCCATGTGGTTTTCCTGGGGCGT

CCTGCTCAACTCCGGCATTGGGGAAGGTGCCCCCCGGAGTTTCTCTGCACGTATCCTAGGCATGGTGT
GGGCTGGTTTCGCCATGATCATAGTGGCTTCCTACACTGCCAACTTGGCAGCTTTCCTGGTGCTGGAT
CGGCCTGAGGAGCGCATCACGGGCATCAATGACCCCAGGCTCAGAAACCCCTCAGACAAGTTCATCTA
CGCAACTGTAAAGCAGAGCTCCGTGGACATCTACTTCCGGAGGCAGGTGGAGTTGAGTACCATGTACC
GGCACATGGAAAAACACAATTACGAGAGCGCAGCTGAGGCCATCCAGGCTGTGCGGGACAACAAGCTG
CACGCCTTTATCTGGGACTCGGCCGTGCTGGAGTTTGAGGCTTCACAGAAGTGCGATCTGGTGACCAC
GGGTGAGCTGTTCTTCCGCTCAGGCTTTGGCATCGGCATGCGCAAGGACAGCCCCTGGAAGCAGAACG
TTTCCCTGTCCATACTCAAGTCCCATGAGAATGGCTTCATGGAAGATCTGGATAAGACATGGGTTCGG
TATCAGGAATGCGACTCCCGCAGCAATGCTCCTGCAACCCTCACTTTTGAGAACATGGCAGGGGTCTT
CATGCTGGTGGCTGGAGGCATCGTAGCTGGGATTTTCCTCATTTTCATTGAGATCGCCTACAAGCGAC
ACAAGGATGCCCGTAGGAAGCAGATGCAGCTGGCTTTTGCAGCCGTGAACGTGTGGAGGAAGAACCTG
CAGGATAGAAAGAGTGGTAGAGCAGAGCCCGACCCTAAAAAGAAAGCCACATTTAGGGCTATCACCTC
CACCCTGGCCTCCAGCTTCAAGAGACGTAGGTCCTCCAAAGACACGAGCACCGGGGGTGGACGCGGCG
CTTTGCAAAACCAAAAAGACACAGTGCTGCCGCGACGCGCTATTGAGAGGGAGGAGGGCCAGCTGCAG
CTGTGTTCCCGTCATAGGGAGAGCTGA

> rnoGrin1-203-3’utr(3’D4nt)
GACGCCCCGCCCGCCCTCCTCTGCCCCTCCCCCGCAGACAGACGCACGGGACAGCGGCCTGGCCCACG
CAGAGCCCCGGAGCACGACGGGGTCGGGGGAGGAGCACTCCCAGCCTCCCCCAGGCCGTGCCCGCCTG
CCCACCGGTCGGCCGGCTGGCCGGTCCACCCTGTCCCGGCCCCGCGCGTGCCCCCGACGTCGGAGCTA
ACGGGCCGCCTTGTCTGTGTATTTCTATTTTACAGCAGTACCATCCCACTGATATCACGGGCCCGCTC
AACCTCTCAGATCCCTCGGTCAGCACCGTGGTGTGAGGCCCCCCGGAGGCGCCCACCTGCCCAGTTAG
CCCGGCCAAGGACACTGATGAGTCCTGCTGCTCGGGAAGGCCTGAGGGAAGCCCACCCGCCCCAGAGA
CTGCCCACCCTGGGCCTCCCGTCCGCCTGCTCTGCTGCCTGGCGGGCAGCCCCTGCAGGACCAAGGTG
CGGACCAGAGCGGCTGAGGATGGGCCAGAGCTGAGCCGGCTGGGCAGGGCCACAGGGCGCTCCGGCAG
AGGCAGGGCCCTGAGGTCTCTGAGCAGTGGGGTGAGGGGCCTAAGTGGCCCCGGTCGGAGGAGTCTGG
AGCAGAAATGGCAGCCCCATCCTTCCTCCAGCCACTACCCCAAGCTACAGTGGGGGCCTATGGCCCCA
GCTTGCTAGGTCACCCCCGACCCTTCCTCCAGCGCCTGCTCTCTGCAACTTGATTTCCACCTCTCTCC
TGCTGCACCACCCTCCCACGACATTTCCCCACCCCATTCACTGGGTTGTCTCTGACCTTTCCCAGGGC
TAGCCTTCACTGCCCTAGTGGCAGTGCTTCAGGGGTGCTTTCTGGCTCCCAGACATCTAGGGCTCCAG
ACTCCAAGAGGGCTGAGCCTTCTCTTCTGTCCGCAGCCACAATAGGCTTCCTCAGACGCTGGCTCGTG
ATGAGTCCCGCACCTTGGGCACCAGGGAGCGCCATCTGCCTCCCAGTCCGGTGTCACTCACCCCACTA
CCTTGTACATGACCAGCTCTCCCAGTGTCCCAGTGTCTGCCCCAGGGACACCGGGCGCGCACAGCCAC
CCCTAATCCCGGTATTCAGTGGTGATGCCTAAAGGAATGTCAGAAAA

>synthetic-DNA-contruct_XbaI-cut
AAAAAAAAAAAGCGGCCGCGGATCCCCTT

**- AgeI/SalI wt.mmuFoxg1aa357-381-V5 module**

>AgeI-cut_3xF-wt.mmuFoxg1$^{aa357-381}$-V5_SalI-cut
GGTGCCACCATGGTGGATTACAAGGACCACGATGGAGATTATAAAGATCATGACATTGACTATAAGGA
TGATGACGACAAAGGGAGTGGCTCCGGAGACCGGCTGGTCAACGGGGAGATCCCGTACGCCACGCACC
ACCTCACGGCCGCTGCACTAGCCGCATCGGTGCCCGGAAGTGGGTCCGGCGGCAAGCCCATCCCTAAC
CCACTGCTCGGATTGGACAGCACATAATAGGTC

**- AgeI/XhoI scr.mmuFoxg1aa357-381-V5 module**

>AgeI-cut_3xF-scr.mmuFoxg1$^{aa357-381}$-V5_XhoI-cut

GGTGCCACCATGGTGGATTACAAGGACCACGATGGAGATTATAAAGATCATGACATTGACTATAAGGA
TGATGACGACAAAGGGAGTGGCTCCGGAGTCGGGCTGCACGAGGCCCCGCTCGCCTACCTACCCGCAT
CGGCAACGGCCCACGCTACGAACGACGTGCGGATCGGAAGTGGGTCCGGCGGCAAGCCCATCCCTAAC
CCACTGCTCGGATTGGACAGCACATAATAGGGATCCCTC

**- "LVrc_TREt-pl-BGHpA" lentivector (full sequence)**

>LVrc_TREt-pl-BGHpA
GGAAATTGTAAACGTTAATATTTTGTTAAAATTCGCGTTAAATTTTTGTTAAATCAGCTCATTTTTTA
ACCAATAGGCCGAAATCGGCAAAATCCCTTATAAATCAAAAGAATAGACCGAGATAGGGTTGAGTGTT
GTTCCAGTTTGGAACAAGAGTCCACTATTAAAGAACGTGGACTCCAACGTCAAAGGGCGAAAAACCGT
CTATCAGGGCGATGGCCCACTACGTGAACCATCACCCTAATCAAGTTTTTTGGGGTCGAGGTGCCGTA
AAGCACTAAATCGGAACCCTAAAGGGAGCCCCCGATTTAGAGCTTGACGGGGAAAGCCGGCGAACGTG
GCGAGAAAGGAAGGGAAGAAAGCGAAAGGAGCGGGCGCTAGGGCGCTGGCAAGTGTAGCGGTCACGCT
GCGCGTAACCACCACACCCGCCGCGCTTAATGCGCCGCTACAGGGCGCGTCGCGCCATTCGCCATTCA
GGCTGCGCAACTGTTGGGAAGGGCGATCGGTGCGGGCCTCTTCGCTATTACGCCAGCTGGCGAAAGGG
GGATGTGCTGCAAGGCGATTAAGTTGGGTAACGCCAGGGTTTTCCCAGTCACGACGTTGTAAAACGAC
GGCCAGTGAGCGCGCGTAATACGACTCACTATAGGGCGAATTGGGTACGTCTCGACGCAAAAGCCTAG
GCCTCCAAAAAAGCCTCCTCACTACTTCTGGAATAGCTCAGAGGCCGAGGCGGCCTCGGCCTCTGCAT
AAATAAAAAAAATTAGTCAGCCATGGGGCGGAGAATGGGCGGAACTGGGCGGAGTTAGGGGCGGGATA
GCTAGAGCCAGACATGATAAGATACATTGATGAGTTTGGACAAACCACAACTAGAATGCAGTGAAAAA
AATGCTTTATTTGTGAAATTTGTGATGCTATTGCTTTATTTGTAACCATTATAAGCTGCAATAAACAA
GTTCCTCTCACTCTCTGATATTCATTTCTTTGCAAGTTATAAATACTGAATAATAAGATGACATGAAC
TACTACTGCTAGAGATTTTCCACACTGACTAAAAGGGTCTGAGGGATCTCTAGTTACCAGAGTCACAC
AACAGACGGGCACACACTACTTGAAGCACTCAAGGCAAGCTTTATTGAGGCTTAAGCAGTGGGTTCCC
TAGTTAGCCAGAGAGCTCCCAGGCTCAGATCTGGTCTAACCAGAGAGACCCAGTACAAGCAAAAAGCA
GATCTTGTCTTCGTTGGGAGTGAATTAGCCCTTCCAGTCCCCCCTTTTCTTTTAAAAAGTGGCTAAGA
TCTACAGCTGCCTTGTAAGTCATTGGTCTTAAAGGTACAAAGGCTAGAGTACTTAATACGAGAGTTTA
CTCCCTATCAGTGATAGAGAACGTATGTCGAGTTTACTCCCTATCAGTGATAGAGAACGATGTCGAGT
TTACTCCCTATCAGTGATAGAGAACGTATGTCGAGTTTACTCCCTATCAGTGATAGAGAACGTATGTC
GAGTTTACTCCCTATCAGTGATAGAGAACGTATGTCGAGTTTATCCCTATCAGTGATAGAGAACGTAT
GTCGAGTTTACTCCCTATCAGTGATAGAGAACGTATGTCGAGGTAGGCGTGTACGGTGGGAGGCCTAT
ATAAGCAGAGCTCGTTTAGTGAACCGTCAGATCGCCCTCGAGTCACAATTGGGCCCTTATGGCCGATT
TGGCCTAACTATAACGGTCCTAAGGTAGCGATCAGGATCCAGTTTAAACTACCGGTCATCTAGACACC
CGGGTGTGGTACCTGTGCCTTCTAGTTGCCAGCCATCTGTTGTTTGCCCCTCCCCCGTGCCTTCCTTG
ACCCTGGAAGGTGCCACTCCCACTGTCCTTTCCTAATAAAATGAGGAAATTGCATCGCATTGTCTGAG
TAGGTGTCATTCTATTCTGGGGGGTGGGGTGGGGCAGGACAGCAAGGGGGAGGATTGGGAAGACAATA
GCAGGCATGCTGGGGATGCGGTGGGCTCTATGGCCCCTTTAGTGAGGGTTAATTGTCGAGGCTAGTCT
CGTGATCGATAAAATTTTGAATTTTTGTAATTTGTTTTTGTAATTCTTTAGTTTGTATGTCTGTTGCT
ATTATGTCTACTATTCTTTCCCCTGCACTGTACCCCCCAATCCCCCCTTTTCTTTTAAAAGTTAACCG
ATACCGTCGAGATCCGTTCACTAATCGAATGGATCTGTCTCTGTCTCTCTCTCCACCTTCTTCTTCTA
TTCCTTCGGGCCTGTCGGGTCCCCTCGGGGTTGGGAGGTGGGTCTGAAACGATAATGGTGAATATCCC
TGCCTAACTCTATTCACTATAGAAAGTACAGCAAAAACTATTCTTAAACCTACCAAGCCTCCTACTAT
CATTATGAATAATTTTATATACCACAGCCAATTTGTTATGTTAAACCAATTCCACAAACTTGCCCATT
TATCTAATTCCAATAATTCTTGTTCATTCTTTTCTTGCTGGTTTTGCGATTCTTCAATTAAGGAGTGT
ATTAAGCTTGTGTAATTGTTAATTTCTCTGTCCCACTCCATCCAGGTCGTGTGATTCCAAATCTGTTC
CAGAGATTTATTACTCCAACTAGCATTCCAAGGCACAGCAGTGGTGCAAATGAGTTTTCCAGAGCAAC
CCCAAATCCCCAGGAGCTGTTGATCCTTTAGGTATCTTTCCACAGCCAGGATTCTTGCCTGGAGCTGC
TTGATGCCCCAGACTGTGAGTTGCAACAGATGCTGTTGCGCCTCAATAGCCCTCAGCAAATTGTTCTG
CTGCTGCACTATACCAGACAATAATTGTCTGGCCTGTACCGTCAGCGTCATTGAGGCTGCGCCCATAG

TGCTTCCTGCTGCTCCCAAGAACCCAAGGAACAAAGCTCCTATTCCCACTGCTCTTTTTCTCTCTGC
ACCACTCTTCTCTTTGCCTTGGTGGGTGCTACTCCTAATGGTTCAATTTTTACTACTTTATATTTATA
TAATTCACTTCTCCAATTGTCCCTCATATCTCCTCCTCCAGGTCTGAAGATCAGCGGCCGCTTGCTGT
GCGGTGGTCTTACTTTTGTTTTGCTCTTCCTCTATCTTGTCTAAAGCTTCCTTGGTGTCTTTTATCTC
TATCCTTTGATGCACACAATAGAGGGTTGCTACTGTATTATATAATGATCTAAGTTCTTCTGATCCTG
TCTGAAGGGATGGTTGTAGCTGTCCCAGTATTTGTCTACAGCCTTCTGATGTTTCTAACAGGCCAGGA
TTAACTGCGAATCGTTCTAGCTCCCTGCTTGCCCATACTATATGTTTTAATTTATATTTTTTCTTTCC
CCCTGGCCTTAACCGAATTTTTTCCCATCGCGATCTAATTCTCCCCCGCTTAATACTGACGCTCTCGC
ACCCATCTCTCTCCTTCTAGCCTCCGCTAGTCAAAATTTTTGGCGTACTCACCAGTCGCCGCCCCTCG
CCTCTTGCCGTGCGCGCTTCAGCAAGCCGAGTCCTGCGTCGAGAGAGCTCTGGTTTCCCTTTCGCTTT
CAGGTCCCTGTTCGGGCGCCACTGCTAGAGATTTTCCACACTGACTAAAAGGGTCTGAGGGATCTCTA
GTTACCAGAGTCACACAACAGACGGGCACACACTACTTGAAGCACTCAAGGCAAGCTTTATTGAGGCT
TAAGCAGTGGGTTCCCTAGTTAGCCAGAGAGCTCCCAGGCTCAGATCTGGTCTAACCAGAGAGACCCC
GGTTCACTAAACGAGCTCTGCTTATATAGACCTCCCACCGTACACGCCTACCGCCCATTTGCGTCAAT
GGGGCGGAGTTGTTACGACATTTTGGAAAGTCCCGTTGATTTTGGTGCCAAAACAAACTCCCATTGAC
GTCAATGGGGTGGAGACTTGGAAATCCCCGTGAGTCAAACCGCTATCCACGCCCATTGATGTACTGCC
AAAACCGCATCACCATGGTAATAGCGATGACTAATACGTAGATGTACTGCCAAGTAGGAAAGTCCCAT
AAGGTCATGTACTGGGCATAATGCCAGGCGGGCCATTTACCGTCATTGACGTCAATAGGGGGCGTACT
TGGCATATGATACACTTGATGTACTGCCAAGTGGGCAGTTTACCGTAAATACTCCACCCATTGACGTC
AATGGAAAGTCCCTATTGGCGTTACTATGGGAACATACGTCATTATTGACGTCAATGGGCGGGGGTCG
TTGGGCGGTCAGCCAGGCGGGCCATTTACCGTAAGTTATGTAACGCGGAACTCCATATATGGGCTATG
AACTAATGACCCCGTAATTGATTACTATTAATAACTAGTCAATAATCAATGTCAACATGGCGGTAATG
TTGGACATGAGCCAATATAAATGTACATATTATGATATGGATACAACGTATGCAATGGCCAAGCTTGC
AGCTCCAGCTTTTGTTCCCTTTAGTGAGGGTTAATTGCGCGCTTGGCGTAATCATGGTCATAGCTGTT
TCCTGTGTGAAATTGTTATCCGCTCACAATTCCACACAACATACGAGCCGGAAGCATAAAGTGTAAAG
CCTGGGGTGCCTAATGAGTGAGCTAACTCACATTAATTGCGTTGCGCTCACTGCCCGCTTTCCAGTCG
GGAAACCTGTCGTGCCAGCTGCATTAATGAATCGGCCAACGCGCGGGGAGAGGCGGTTTGCGTATTGG
GCGCTCTTCCGCTTCCTCGCTCACTGACTCGCTGCGCTCGGTCGTTCGGCTGCGGCGAGCGGTATCAG
CTCACTCAAAGGCGGTAATACGGTTATCCACAGAATCAGGGGATAACGCAGGAAAGAACATGTGAGCA
AAAGGCCAGCAAAAGGCCAGGAACCGTAAAAAGGCCGCGTTGCTGGCGTTTTTCCATAGGCTCCGCCC
CCCTGACGAGCATCACAAAAATCGACGCTCAAGTCAGAGGTGGCGAAACCCGACAGGACTATAAAGAT
ACCAGGCGTTTCCCCCTGGAAGCTCCCTCGTGCGCTCTCCTGTTCCGACCCTGCCGCTTACCGGATAC
CTGTCCGCCTTTCTCCCTTCGGGAAGCGTGGCGCTTTCTCATAGCTCACGCTGTAGGTATCTCAGTTC
GGTGTAGGTCGTTCGCTCCAAGCTGGGCTGTGTGCACGAACCCCCCGTTCAGCCCGACCGCTGCGCCT
TATCCGGTAACTATCGTCTTGAGTCCAACCCGGTAAGACACGACTTATCGCCACTGGCAGCAGCCACT
GGTAACAGGATTAGCAGAGCGAGGTATGTAGGCGGTGCTACAGAGTTCTTGAAGTGGTGGCCTAACTA
CGGCTACACTAGAAGGACAGTATTTGGTATCTGCGCTCTGCTGAAGCCAGTTACCTTCGGAAAAAGAG
TTGGTAGCTCTTGATCCGGCAAACAAACCACCGCTGGTAGCGGTGGTTTTTTTGTTTGCAAGCAGCAG
ATTACGCGCAGAAAAAAAGGATCTCAAGAAGATCCTTTGATCTTTTCTACGGGGTCTGACGCTCAGTG
GAACGAAAACTCACGTTAAGGGATTTTGGTCATGAGATTATCAAAAAGGATCTTCACCTAGATCCTTT
TAAATTAAAAATGAAGTTTTAAATCAATCTAAAGTATATATGAGTAAACTTGGTCTGACAGTTACCAA
TGCTTAATCAGTGAGGCACCTATCTCAGCGATCTGTCTATTTCGTTCATCCATAGTTGCCTGACTCCC
CGTCGTGTAGATAACTACGATACGGGAGGGCTTACCATCTGGCCCCAGTGCTGCAATGATACCGCGAG
ACCCACGCTCACCGGCTCCAGATTTATCAGCAATAAACCAGCCAGCCGGAAGGGCCGAGCGCAGAAGT
GGTCCTGCAACTTTATCCGCCTCCATCCAGTCTATTAATTGTTGCCGGGAAGCTAGAGTAAGTAGTTC
GCCAGTTAATAGTTTGCGCAACGTTGTTGCCATTGCTACAGGCATCGTGGTGTCACGCTCGTCGTTTG
GTATGGCTTCATTCAGCTCCGGTTCCCAACGATCAAGGCGAGTTACATGATCCCCCATGTTGTGCAAA
AAAGCGGTTAGCTCCTTCGGTCCTCCGATCGTTGTCAGAAGTAAGTTGGCCGCAGTGTTATCACTCAT
GGTTATGGCAGCACTGCATAATTCTCTTACTGTCATGCCATCCGTAAGATGCTTTTCTGTGACTGGTG
AGTACTCAACCAAGTCATTCTGAGAATAGTGTATGCGGCGACCGAGTTGCTCTTGCCCGGCGTCAATA
CGGGATAATACCGCGCCACATAGCAGAACTTTAAAAGTGCTCATCATTGGAAAACGTTCTTCGGGGCG

AAAACTCTCAAGGATCTTACCGCTGTTGAGATCCAGTTCGATGTAACCCACTCGTGCACCCAACTGAT
CTTCAGCATCTTTTACTTTCACCAGCGTTTCTGGGTGAGCAAAAACAGGAAGGCAAAATGCCGCAAAA
AAGGGAATAAGGGCGACACGGAAATGTTGAATACTCATACTCTTCCTTTTTCAATATTATTGAAGCAT
TTATCAGGGTTATTGTCTCATGAGCGGATACATATTTGAATGTATTTAGAAAAATAAACAAATAGGGG
TTCCGCGCACATTTCCCCGAAAAGTGCCACCTG

**Table S4. Full primary data.** Includes full primary data referred to by Figures 2, 3, 4, 5, 6, 7, 8, 9, 11, S1 and S2

### FIGURE 2

### B

| | TRAP/SN | *Foxg1 (5'utr)* | *Slc17a6* | *Gria1* | *Gabra1* | *Grid1* | *Grin1* | *Bdnf-2c* | *Bdnf-4* | *Psd95* |
|---|---|---|---|---|---|---|---|---|---|---|
| *sample values* | Ctr.1 | 1.19 | 1.13 | 1.05 | 1.62 | 1.23 | 1.21 | 1.01 | 1.28 | 1.44 |
| | Ctr.2 | 1.00 | 1.08 | 1.35 | 1.19 | 1.12 | 1.14 | 1.06 | 0.88 | 0.92 |
| | Ctr.3 | 0.64 | 0.89 | 0.87 | 0.77 | 0.99 | 0.87 | 1.31 | 0.92 | 0.71 |
| | Ctr.4 | 0.78 | 0.99 | 0.91 | 0.82 | 0.86 | 0.82 | 0.62 | 0.93 | 0.87 |
| | Ctr.5 | 0.79 | 1.25 | 1.30 | 0.98 | 1.07 | 1.23 | | | 1.51 |
| | Ctr.6 | 1.59 | 0.66 | 0.52 | 0.62 | 0.72 | 0.72 | | | 0.55 |
| | *Foxg1* -OE.1 | 1.54 | 0.66 | 1.28 | 0.82 | 2.07 | 1.31 | 0.70 | 0.54 | 1.64 |
| | *Foxg1* -OE.2 | 0.81 | 0.77 | 0.97 | 0.56 | 1.12 | 1.28 | 0.92 | 0.75 | 1.26 |
| | *Foxg1* -OE.3 | 0.64 | 1.21 | 1.01 | 0.82 | 1.00 | 1.10 | 1.61 | 0.74 | 0.82 |
| | *Foxg1* -OE.4 | 1.23 | 1.85 | 1.13 | 0.99 | 1.54 | 1.61 | 0.41 | 1.06 | 1.46 |
| | *Foxg1* -OE.5 | 0.67 | 1.06 | 1.13 | 1.24 | 1.56 | 1.33 | | | 1.01 |
| | *Foxg1* -OE.6 | 0.72 | 1.21 | 1.58 | 1.01 | 1.74 | 1.45 | | | 1.39 |
| *average* | Ctr | 1.00 | 1.00 | 1.00 | 1.00 | 1.00 | 1.00 | 1.00 | 1.00 | 1.00 |
| | *Foxg1* -OE | 0.93 | 1.13 | 1.19 | 0.91 | 1.51 | 1.35 | 0.91 | 0.77 | 1.26 |
| *sem* | Ctr | 0.14 | 0.08 | 0.13 | 0.15 | 0.08 | 0.09 | 0.14 | 0.09 | 0.16 |
| | *Foxg1* -OE | 0.15 | 0.17 | 0.09 | 0.09 | 0.16 | 0.07 | 0.26 | 0.11 | 0.12 |
| *p* | | 0.38 | 0.26 | 0.13 | 0.30 | 0.01 | 0.01 | 0.39 | 0.08 | 0.11 |

### D

| | protein | *ex20-Grin1* |
|---|---|---|
| *sample values* | Ctr.1 | 1.01 |
| | Ctr.2 | 1.20 |
| | Ctr.3 | 1.00 |
| | Ctr.4 | 0.78 |
| | *Foxg1* -LOF.1 | 0.67 |
| | *Foxg1* -LOF.2 | 0.50 |
| | *Foxg1* -LOF.3 | 0.64 |
| | *Foxg1* -LOF.4 | 0.72 |
| *average* | Ctr | 1.00 |
| | *Foxg1* -LOF | 0.63 |
| *sem* | Ctr | 0.14 |
| | *Foxg1* -LOF | 0.07 |
| *p* | | 0.005 |

| | mRNA | *pan-Grin1* | *ex20-Grin1* |
|---|---|---|---|
| *sample values* | Ctr.1 | 1.04 | 0.88 |
| | Ctr.2 | 0.97 | 1.15 |
| | Ctr.3 | 1.01 | 1.01 |
| | Ctr.4 | 0.97 | 0.95 |
| | *Foxg1* -LOF.1 | 1.03 | 1.08 |
| | *Foxg1* -LOF.2 | 1.12 | 1.68 |
| | *Foxg1* -LOF.3 | 1.05 | 1.28 |
| | *Foxg1* -LOF.4 | 1.07 | 1.22 |
| *average* | Ctr | 1.00 | 1.00 |
| | *Foxg1* -LOF | 1.07 | 1.31 |
| *sem* | Ctr | 0.02 | 0.06 |
| | *Foxg1* -LOF | 0.02 | 0.13 |
| *p* | | 0.02 | 0.03 |

### E

| | protein | *ex20-Grin1* |
|---|---|---|
| *sample values* | Ctr.1 | 0.97 |
| | Ctr.2 | 0.97 |
| | Ctr.3 | 1.06 |
| | Ctr.4 | 1.00 |
| | *Foxg1* -OE.1 | 1.16 |
| | *Foxg1* -OE.2 | 1.22 |
| | *Foxg1* -OE.3 | 1.07 |
| | *Foxg1* -OE.4 | 1.06 |
| *average* | Ctr | 1.00 |
| | *Foxg1* -LOF | 1.13 |
| *sem* | Ctr | 0.02 |
| | *Foxg1* -LOF | 0.04 |
| *p* | | 0.01 |

| | mRNA | *pan-Grin1* | *ex20-Grin1* |
|---|---|---|---|
| *sample values* | Ctr.1 | 0.97 | 1.10 |
| | Ctr.2 | 0.97 | 0.99 |
| | Ctr.3 | 0.94 | 0.87 |
| | Ctr.4 | 1.12 | 1.04 |
| | *Foxg1* -OE.1 | 0.75 | 0.77 |
| | *Foxg1* -OE.2 | 0.94 | 1.03 |
| | *Foxg1* -OE.3 | 0.82 | 0.75 |
| | *Foxg1* -OE.4 | 0.94 | 0.74 |
| *average* | Ctr | 1.00 | 1.00 |
| | *Foxg1* -OE | 0.86 | 0.82 |
| *sem* | Ctr | 0.04 | 0.05 |
| | *Foxg1* -OE | 0.05 | 0.07 |
| *p* | | 0.03 | 0.04 |

### E

| Grin1 protein/ Grin1-mRNA | *pan-Grin1 norm* |
|---|---|
| *Foxg1* -LOF | 0.59 |
| Ctr | 1.00 |
| *Foxg1* -OE | 1.31 |

| Grin1 protein/ Grin1-mRNA | *ex20-Grin1 norm* |
|---|---|
| *Foxg1* -LOF | 0.48 |
| Ctr | 1.00 |
| *Foxg1* -OE | 1.37 |

### FIGURE 3

### A

(a)

| | nascent Grin1 protein (anti-NH2-term) | *whole neuron* *cumulative signal per cell* | *whole neuron* *cumulative signal per spot* | *neurites* *cumulative signal per spot* |
|---|---|---|---|---|
| *sample values* | Ctr.1 | 1.04 | 1.03 | 1.01 |
| | Ctr.2 | 1.08 | 1.15 | 1.00 |
| | Ctr.3 | 1.06 | 1.06 | 1.04 |
| | Ctr.4 | 0.91 | 0.83 | 0.80 |
| | Ctr.5 | 1.08 | 0.92 | 0.88 |
| | Ctr.6 | 0.93 | 1.04 | 1.20 |
| | Ctr.7 | 0.90 | 0.98 | 1.08 |
| | *Foxg1* -LOF.1 | 0.93 | 0.81 | 0.84 |
| | *Foxg1* -LOF.2 | 0.97 | 0.88 | 0.92 |
| | *Foxg1* -LOF.3 | 0.90 | 0.84 | 0.92 |
| | *Foxg1* -LOF.4 | 1.00 | 0.93 | 0.97 |
| | *Foxg1* -LOF.5 | 0.86 | 0.75 | 0.81 |
| | *Foxg1* -LOF.6 | 0.96 | 0.98 | 1.04 |
| | *Foxg1* -LOF.7 | 0.87 | 0.76 | 0.73 |
| | *Foxg1* -LOF.8 | 0.97 | 0.79 | 0.91 |
| *average* | Ctr | 1.00 | 1.00 | 1.00 |
| | *Foxg1* -LOF | 0.93 | 0.84 | 0.89 |
| *sem* | Ctr | 0.03 | 0.04 | 0.05 |
| | *Foxg1* -LOF | 0.02 | 0.03 | 0.03 |
| *p* | | 0.04 | 0.003 | 0.04 |

(b)

| | nascent Grin1 protein (anti-COOH-term) | whole neuron *cumulative signal per cell* | whole neuron *cumulative signal per spot* | neurites *cumulative signal per spot* |
|---|---|---|---|---|
| *sample values* | Ctr.1 | 0.72 | 0.87 | 0.91 |
| | Ctr.2 | 1.18 | 1.09 | 1.01 |
| | Ctr.3 | 1.16 | 1.04 | 1.09 |
| | Ctr.4 | 0.96 | 0.96 | 1.01 |
| | Ctr.5 | 0.92 | 0.94 | 0.91 |
| | Ctr.6 | 1.14 | 1.02 | 1.11 |
| | Ctr.7 | 0.87 | 1.06 | 1.05 |
| | Ctr.8 | 1.05 | 1.02 | 0.91 |
| | *Foxg1* -LOF.1 | 0.90 | 0.95 | 0.93 |
| | *Foxg1* -LOF.2 | 0.78 | 0.80 | 0.99 |
| | *Foxg1* -LOF.3 | 0.75 | 0.85 | 0.88 |
| | *Foxg1* -LOF.4 | 0.71 | 0.87 | 0.87 |
| | *Foxg1* -LOF.5 | 0.87 | 0.91 | 1.01 |
| | *Foxg1* -LOF.6 | 0.85 | 0.89 | 0.98 |
| | *Foxg1* -LOF.7 | 1.02 | 0.94 | 0.97 |
| | *Foxg1* -LOF.8 | 0.44 | 0.63 | 0.74 |
| *average* | Ctr | 1.00 | 1.00 | 1.00 |
| | *Foxg1* -LOF | 0.79 | 0.86 | 0.92 |
| *sem* | Ctr | 0.06 | 0.03 | 0.03 |
| | *Foxg1* -LOF | 0.06 | 0.04 | 0.03 |
| *p* | | 0.01 | 0.003 | 0.05 |

### B

| | anti-puromycin | *average signal level* |
|---|---|---|
| sample values | Ctr.1 | 1.00 |
| | Ctr.2 | 0.98 |
| | Ctr.3 | 1.02 |
| | *Foxg1* -LOF.1 | 0.98 |
| | *Foxg1* -LOF.2 | 0.99 |
| | *Foxg1* -LOF.3 | 0.98 |
| *average* | Ctr | 1.00 |
| | *Foxg1* -LOF | 0.98 |
| *sem* | Ctr | 0.012 |
| | *Foxg1* -LOF | 0.004 |
| p | | 0.09 |

### C

*sample values*

| | | *t0* | | *t6* | | *t10* | | *t14* | |
|---|---|---|---|---|---|---|---|---|---|
| | | Ctr | *Foxg1* -LOF | Ctr | *Foxg1* -LOF | Ctr | *Foxg1* -LOF | Ctr | *Foxg1* -LOF |
| **Grin1 protein level** | *#1* | 1.00 | 0.69 | 0.73 | 0.76 | 0.67 | 1.03 | 0.31 | 0.55 |
| | *#2* | 0.96 | 0.99 | 1.11 | 0.89 | 0.33 | 0.85 | 0.68 | 0.45 |
| | *#3* | 1.03 | 1.32 | 1.13 | 0.95 | 0.52 | 0.68 | 0.28 | 0.41 |

*p (ANCOVA)*
0.0468

### FIGURE 4

### A

| | | *sample values* | | | | | | |
|---|---|---|---|---|---|---|---|---|
| | | *t0* | | *t11* | | *average t11/t0* | | |
| | | Ctr | *Foxg1 -* LOF | Ctr | *Foxg1 -* LOF | Ctr | *Foxg1 -* LOF | *p* |
| **nascent Grin1 protein (cumulative signal per spot)** | *#1* | 1.06 | 0.91 | 0.63 | 0.63 | 1.00 | 0.96 | 0.15 |
| | *#2* | 1.00 | 0.95 | 0.65 | 0.65 | | | |
| | *#3* | 0.88 | 0.95 | 0.65 | 0.62 | | | |
| | *#4* | 1.06 | 1.15 | 0.70 | | | | |
| | *#5* | | 1.04 | | | | | |

### B

| | | **mRNA** | | **IF level** | | | | | | | |
|---|---|---|---|---|---|---|---|---|---|---|---|
| | | **(a)** | | **(b)** | | **(c)** | | **(e)** | | **(d)** | |
| | | *Flag-rnoGrin1-HA* | | *Flag-rnoGrin1-HA, N-term* | | *Flag-rnoGrin1-HA, C-term* | | *Flag-rnoGrin1-HA, C-term/N-term* | | *Flag-rnoGrin1-HA,N-term/mRNA* | |
| | | *Foxg1 -* LOF.1 | Ctr.1 | *Foxg1 -* LOF | Ctr | *Foxg1 -* LOF | Ctr | *Foxg1 -* LOF | Ctr | *Foxg1 -* LOF | Ctr |
| | *#1* | 0.97 | 0.45 | 0.77 | 0.44 | 0.59 | 0.50 | 0.74 | 1.12 | | |
| | *#2* | 0.98 | 0.32 | 3.02 | 0.38 | 3.48 | 1.02 | 1.12 | 2.61 | | |
| | *#3* | 1.23 | 0.49 | 0.74 | 0.14 | 0.72 | 0.16 | 0.95 | 1.06 | | |
| | *#4* | 0.82 | 0.90 | 0.29 | 1.90 | 0.30 | 3.85 | 0.99 | 1.97 | | |
| | *#5* | | 0.86 | 0.62 | 0.16 | 0.63 | 0.21 | 1.00 | 1.28 | | |
| | *#6* | | | 0.43 | 0.46 | 0.53 | 0.49 | 1.18 | 1.03 | | |
| | *#7* | | | 0.09 | 0.16 | 0.11 | 0.32 | 1.23 | 1.93 | | |
| | *#8* | | | 2.74 | 0.77 | 2.86 | 1.01 | 1.01 | 1.27 | | |
| | *#9* | | | 0.56 | 1.06 | 0.75 | 1.38 | 1.30 | 1.26 | | |
| | *#10* | | | 0.42 | 0.24 | 0.50 | 0.27 | 1.18 | 1.12 | | |
| | *#11* | | | 1.14 | 0.27 | 1.16 | 0.25 | 0.99 | 0.93 | | |
| | *#12* | | | 1.49 | 2.18 | 1.47 | 2.08 | 0.95 | 0.93 | | |
| | *#13* | | | 1.40 | 0.66 | 1.46 | 0.67 | 1.01 | 0.99 | | |
| | *#14* | | | 0.80 | 2.53 | 0.79 | 2.36 | 0.97 | 0.90 | | |
| | *#15* | | | 1.64 | 0.74 | 2.31 | 0.97 | 1.37 | 1.27 | | |
| | *#16* | | | 1.51 | 1.48 | 1.40 | 1.50 | 0.90 | 0.98 | | |
| | *#17* | | | 0.28 | 0.74 | 0.25 | 0.85 | 0.86 | 1.10 | | |
| | *#18* | | | 1.71 | 0.27 | 1.60 | 0.39 | 0.91 | 1.42 | | |
| | *#19* | | | 1.80 | 0.29 | 1.60 | 0.29 | 0.86 | 0.98 | | |
| | *#20* | | | 0.15 | 0.27 | 0.19 | 0.51 | 1.21 | 1.84 | | |
| | *#21* | | | 1.87 | 0.13 | 1.70 | 0.14 | 0.88 | 1.00 | | |
| | *#22* | | | 0.08 | 0.41 | 0.12 | 0.47 | 1.41 | 1.11 | | |
| | *#23* | | | 2.50 | 0.70 | 2.57 | 0.80 | 0.99 | 1.10 | | |
| | *#24* | | | 1.87 | 3.45 | 2.10 | 3.49 | 1.09 | 0.98 | | |
| | *#25* | | | 0.28 | 0.15 | 0.30 | 0.20 | 1.02 | 1.32 | | |
| | *#26* | | | 0.33 | 0.83 | 0.33 | 0.88 | 0.96 | 1.02 | | |
| | *#27* | | | 1.91 | 0.13 | 1.92 | 0.14 | 0.98 | 1.05 | | |
| | *#28* | | | 0.27 | 1.63 | 0.28 | 2.08 | 1.00 | 1.24 | | |
| | *#29* | | | 0.15 | 0.25 | 0.14 | 0.26 | 0.95 | 1.03 | | |
| | *#30* | | | 0.49 | 0.37 | 0.47 | 0.45 | 0.93 | 1.20 | | |
| | *#31* | | | 0.46 | 2.52 | 0.79 | 2.98 | 1.68 | 1.15 | | |
| | *#32* | | | 0.51 | 0.60 | 0.48 | 0.52 | 0.91 | 0.83 | | |
| | *#33* | | | 0.24 | 2.06 | 0.21 | 2.19 | 0.83 | 1.03 | | |
| *sample values* | *#34* | | | 1.24 | 0.76 | 1.30 | 0.74 | 1.01 | 0.94 | | |
| | *#35* | | | 0.25 | 0.50 | 0.43 | 0.51 | 1.66 | 0.98 | | |
| | *#36* | | | 0.84 | 1.12 | 0.86 | 1.15 | 1.00 | 1.00 | | |
| | *#37* | | | 2.41 | 3.27 | 2.48 | 3.53 | 1.00 | 1.05 | | |
| | *#38* | | | 1.34 | 1.06 | 1.14 | 0.89 | 0.83 | 0.81 | | |
| | *#39* | | | 1.58 | 1.65 | 1.66 | 1.79 | 1.02 | 1.05 | | |
| | *#40* | | | 1.04 | 0.39 | 1.11 | 0.73 | 1.03 | 1.83 | | |
| | *#41* | | | 0.99 | 0.38 | 0.83 | 0.49 | 0.82 | 1.23 | | |
| | *#42* | | | 0.19 | 0.85 | 0.13 | 0.86 | 0.68 | 0.99 | | |
| | *#43* | | | 0.12 | 0.26 | 0.11 | 0.25 | 0.91 | 0.91 | | |
| | *#44* | | | 2.74 | 0.70 | 2.21 | 0.89 | 0.79 | 1.24 | | |
| | *#45* | | | 0.54 | 0.22 | 0.38 | 0.25 | 0.68 | 1.10 | | |
| | *#46* | | | 2.05 | 0.74 | 1.69 | 0.78 | 0.80 | 1.03 | | |
| | *#47* | | | 0.31 | 1.59 | 0.33 | 1.66 | 1.03 | 1.01 | | |
| | *#48* | | | 0.92 | 0.39 | 0.69 | 0.43 | 0.73 | 1.06 | | |
| | *#49* | | | 2.04 | 2.16 | 1.66 | 2.02 | 0.79 | 0.91 | | |
| | *#50* | | | 0.39 | 0.42 | 0.34 | 0.37 | 0.85 | 0.84 | | |
| | *#51* | | | 0.50 | 0.21 | 0.43 | 0.24 | 0.83 | 1.07 | | |
| | *#52* | | | 0.49 | 1.65 | 0.45 | 2.16 | 0.90 | 1.27 | | |
| | *#53* | | | 0.47 | 1.10 | 0.46 | 1.37 | 0.94 | 1.21 | | |
| | *#54* | | | 0.27 | 0.29 | 0.38 | 0.31 | 1.35 | 1.03 | | |
| | *#55* | | | 1.37 | 1.22 | 1.25 | 1.24 | 0.89 | 0.99 | | |
| | *#56* | | | 1.34 | 0.29 | 1.55 | 0.49 | 1.12 | 1.67 | | |
| | *#57* | | | | 0.96 | | 1.41 | | 1.44 | | |
| | *#58* | | | | 0.61 | | 0.72 | | 1.14 | | |
| | *#59* | | | | 1.40 | | 1.34 | | 0.93 | | |
| | *#60* | | | | 1.28 | | 1.22 | | 0.93 | | |
| | *#61* | | | | 1.47 | | 1.34 | | 0.88 | | |
| | *#62* | | | | 0.60 | | 0.57 | | 0.93 | | |
| | *#63* | | | | 2.00 | | 2.34 | | 1.13 | | |
| | *#64* | | | | 0.21 | | 0.29 | | 1.35 | | |
| | *#65* | | | | 0.93 | | 0.95 | | 0.99 | | |
| | *#66* | | | | 1.67 | | 2.47 | | 1.43 | | |
| | *#67* | | | | 1.71 | | 2.21 | | 1.26 | | |
| | *#68* | | | | 1.61 | | 2.04 | | 1.23 | | |
| *average* | | 1.00 | 0.60 | 1.00 | 0.94 | 1.00 | 1.08 | 1.00 | 1.16 | 1.00 | 1.56 |
| *sem* | | 0.08 | 0.12 | 0.11 | 0.09 | 0.11 | 0.11 | 0.03 | 0.04 | --- | --- |
| *p* | | 0.02 | | 0.34 | | 0.29 | | 0.0006 | | --- | |

### FIGURE 5

**A**

**cumulative PLA signal per mCherry+ cell**

| | | *sample values* | | | | | |
|---|---|---|---|---|---|---|---|
| | | *#1* | *#2* | *#3* | *average* | *sem* | *p* |
| | *plasmids: a,b1,c* | | | | | | |
| | aFoxg1,aV5 | 2.76 | 4.03 | | 3.40 | 0.64 | |
| *antibodies* | aFoxg1 | 1.05 | 1.14 | | 1.09 | 0.05 | 0.03 |
| | aV5 | 0.42 | 1.40 | | 0.91 | 0.49 | 0.046 |
| | *plasmids: a,b2,c* | | | | | | |
| | aFoxg1,aV5 | 0.78 | 3.00 | 1.08 | 1.62 | 0.69 | |
| *antibodies* | aFoxg1 | 0.55 | 1.34 | | 0.95 | 0.40 | 0.26 |
| | aV5 | 0.69 | 1.41 | | 1.05 | 0.36 | 0.29 |
| | *plasmids: a,b3,c* | | | | | | |
| | aFoxg1,aFlag | 4.56 | 2.92 | | 3.74 | 0.82 | |
| *antibodies* | aFoxg1 | 1.04 | 0.46 | | 0.75 | 0.29 | 0.04 |
| | aFlag | 1.12 | 1.38 | | 1.25 | 0.13 | 0.048 |
| | *plasmids: a,b4,c* | | | | | | |
| | aFoxg1,aV5 | 1.67 | 1.49 | | 1.58 | 0.09 | |
| *antibodies* | aFoxg1 | 0.93 | 0.60 | | 0.76 | 0.16 | 0.02 |
| | aV5 | 1.80 | 0.67 | | 1.24 | 0.56 | 0.30 |

**B**

| **co-IP.1** | | *IP-Foxg1/IN-Foxg1*IN-Flag-bait* | **co-IP.2** | | IP-Foxg1/IN-Foxg1*IN-Flag-bait |
|---|---|---|---|---|---|
| *sample values* | Flag-Gephyrin.1 | 0.88 | *sample values* | Flag-GFP.1 | 0.59 |
| | Flag-Gephyrin.2 | 1.12 | | Flag-GFP.2 | 1.41 |
| | Flag-eIF4E.1 | 99.25 | | Flag-eIF4E.1 | 35.59 |
| | Flag-eIF4E.2 | 99.02 | | Flag-eIF4E.2 | 34.76 |
| *average* | Flag-Gephyrin | 1.00 | *average* | Flag-GFP | 1.00 |
| | Flag-eIF4E | 99.13 | | Flag-eIF4E | 35.17 |
| *sem* | Flag-Gephyrin | 0.12 | *sem* | Flag-GFP | 0.41 |
| | Flag-eIF4E | 0.12 | | Flag-eIF4E | 0.41 |
| *p* | | 0.000001 | *p* | | 0.0001 |

### FIGURE 6

### A

**(a)**

| | PLA-spots per cell, whole cells | sample values #1 | #2 | #3 | #4 | average | sem | p |
|---|---|---|---|---|---|---|---|---|
| | a-eIF4E, aFoxg1 | 4.60 | 4.80 | 2.27 | 2.79 | 3.62 | 0.64 | |
| *antibodies* | a-eIF4E | 1.58 | 2.46 | 1.26 | 0.62 | 1.48 | 0.38 | 0.01 |
| | aFoxg1 | 0.60 | 0.57 | 0.49 | 0.41 | 0.52 | 0.04 | 0.01 |

**(b)**

| | PLA-spots per cell, neurites | sample values #1 | #2 | #3 | #4 | average | sem | p |
|---|---|---|---|---|---|---|---|---|
| | a-eIF4E, aFoxg1 | 4.20 | 3.18 | 2.06 | 3.21 | 3.17 | 0.44 | |
| *antibodies* | a-eIF4E | 2.22 | 2.22 | 1.44 | 0.76 | 1.66 | 0.35 | 0.02 |
| | aFoxg1 | 0.45 | 0.22 | 0.35 | 0.34 | 0.34 | 0.05 | 0.003 |

### B

**(a)**

| | PLA (aFoxg1, a-eIF4E) | *PLA-signal per cell* |
|---|---|---|
| *sample values* | LV_a.1 | 0.94 |
| | LV_a.2 | 1.12 |
| | LV_a.3 | 0.94 |
| | LV_b.1 | 0.90 |
| | LV_b.2 | 0.77 |
| | LV_b.3 | 0.84 |
| *average* | LV_a | 1.00 |
| | LV_b | 0.84 |
| *sem* | LV_a | 0.06 |
| | LV_b | 0.04 |
| *p* | | 0.04 |

**(b)**

| | nascent Grin1 protein | *PLA-signal per cell* |
|---|---|---|
| *sample values* | LV_a.1 | 0.90 |
| | LV_a.2 | 1.15 |
| | LV_a.3 | 0.95 |
| | LV_b.1 | 0.69 |
| | LV_b.2 | 0.75 |
| | LV_b.3 | 0.66 |
| *average* | LV_a | 1.00 |
| | LV_b | 0.70 |
| *sem* | LV_a | 0.08 |
| | LV_b | 0.03 |
| *p* | | 0.01 |

### FIGURE 7

### A

**(a)**

| | | IP/IN | *Grin1* |
|---|---|---|---|
| *sample values* | IgG | #1 | 1.00 |
| | | #2 | 1.00 |
| | | #3 | 0.98 |
| | | #4 | 1.02 |
| | aFoxg1 | #1 | 7.52 |
| | | #2 | 35.87 |
| | | #3 | 35.29 |
| | | #4 | 4.40 |
| | | #5 | 4.71 |
| *average* | IgG | | 1.00 |
| | aFoxg1 | | 17.56 |
| *sem* | IgG | | 0.01 |
| | aFoxg1 | | 7.38 |
| *p* | | | 0.04 |

**(b)**

| | Grin1 IP/IN | *IgG (Ctr)* | *aFoxg1 (Ctr)* | *aFoxg1 (Foxg1-LOF)* |
|---|---|---|---|---|
| *sample values* | #1 | 1.14 | 11.47 | 6.65 |
| | #2 | 0.86 | 28.81 | 8.86 |
| *average* | | 1.00 | 20.14 | 7.76 |
| *sem* | | 0.14 | 8.67 | 1.11 |
| *p* | IgG (Ctr) | | | |
| | aFoxg1 (Ctr) | 0.079 | | |
| | aFoxg1 (Foxg1-LOF) | 0.013 | 0.146 | |

**(c)**

| | | IP(aEGFP)/IN | *Grin1* |
|---|---|---|---|
| *sample values* | Ctr | #1 | 1.06 |
| | | #2 | 0.94 |
| | Foxg1-GFP | #1 | 6.94 |
| | | #2 | 5.32 |
| *average* | Ctr | | 1.00 |
| | *Foxg1-GFP* | | 6.13 |
| *sem* | Ctr | | 0.06 |
| | *Foxg1-GFP* | | 0.81 |
| *p* | | | 0.01 |

### B

| rnoGrin1-203* IP/IN | #1 | #2 | #3 | #4 | #5 | #6 | #7 | #8 | #9 | #10 | #11 | #12 | #13 | #14 | *aver* | *sem* | *p (vs-d0)* |
|---|---|---|---|---|---|---|---|---|---|---|---|---|---|---|---|---|---|
| | *sample values* | | | | | | | | | | | | | | | | |
| *d0* | 0.80 | 1.35 | 1.07 | 0.77 | 1.01 | 1.06 | 1.02 | 0.91 | 0.75 | 1.25 | 1.00 | 1.00 | 0.44 | 1.56 | 1.00 | 0.08 | |
| *d1* | 0.73 | 0.76 | 0.91 | 0.72 | | | | | | | | | | | 0.78 | 0.04 | 0.07 |
| *d2* | 1.06 | 1.00 | | | | | | | | | | | | | 1.03 | 0.03 | 0.44 |
| *d3* | 0.43 | 0.60 | 0.57 | 0.51 | | | | | | | | | | | 0.53 | 0.04 | 0.002 |
| *d4* | 2.27 | 2.52 | | | | | | | | | | | | | 2.39 | 0.13 | ######## |
| *d5* | 0.12 | 0.49 | | | | | | | | | | | | | 0.30 | 0.19 | 0.002 |

### FIGURE 8

| | nascent Grin1 protein | *wt_Ctr* | *wt_K5* | *wt_K10-noK25* | *Foxg1-LOF_Ctr* | *Foxg1.LOF_K5* | *Foxg1.LOF_K10-noK25* |
|---|---|---|---|---|---|---|---|
| *samples values* | *#1* | 0.79 | 0.06 | 0.53 | 0.57 | 0.07 | 0.10 |
| | *#2* | 1.14 | 0.04 | 0.57 | 0.44 | 0.11 | 0.21 |
| | *#3* | 1.07 | 0.03 | 0.67 | 0.49 | 0.09 | 0.36 |
| | *#4* | | | | | 0.09 | 0.42 |
| *average* | | 1.00 | 0.04 | 0.59 | 0.50 | 0.09 | 0.27 |
| *sem* | | 0.108 | 0.009 | 0.043 | 0.037 | 0.008 | 0.073 |
| *p* | wt_Ctr | | | | | | |
| | wt_K5 | 0.0005 | | | | | |
| | wt_K10-noK25 | 0.0123 | 0.0001 | | | | |
| | Foxg1-LOF_Ctr | 0.0060 | | | | | |
| | Foxg1.LOF_K5 | | 0.0064 | | 0.00003 | | |
| | Foxg1.LOF_K10-noK25 | | | 0.0096 | 0.0285 | 0.0236 | |

| Interaction test | |
|---|---|
| *2-ways ANOVA genotype (wt;foxg1-LOF)-vs-stimulation(Ctr;K5)* | 0.0019 |
| *2-ways ANOVA genotype (wt;foxg1-LOF)-vs-stimulation(K5;K10noK25)* | 0.0033 |
| *2-ways ANOVA genotype (wt;foxg1-LOF)-vs-stimulation(Ctr;K10noK25)* | 1 |

### FIGURE 9

**C**

| gene | *Zfp637* | *Ints9* | *Atp11c* | *Nr4a3* | *Nup93* | *Grik2* | *Kif26b* | *Phb2* | *Sgk1* | *Homer1* | *Cacna2d1* | *Nsa2* | *Ddx1* | *Syt4* | *Gabra2* | *Fez1* | *Sox4* |
|---|---|---|---|---|---|---|---|---|---|---|---|---|---|---|---|---|---|
| *Log2FC(totRNA)* | -0.27 | 0.01 | -0.42 | -3.88 | -0.56 | -2.07 | 0.21 | -0.18 | -0.43 | -0.53 | -1.59 | -0.16 | -0.01 | -1.72 | -1.59 | -0.36 | -1.33 |
| *ΔLog2FC* | 3.03 | 3.02 | 2.26 | 3.04 | 2.63 | 2.45 | 1.36 | 1.22 | 1.90 | 1.56 | 1.34 | 0.83 | 0.74 | 0.94 | 0.88 | 0.78 | 0.75 |

| gene | *Nmt1* | *Cltb* | *Cdc40* | *Ccnt2* | *Dbr1* |
|---|---|---|---|---|---|
| *Log2FC(totRNA)* | -0.06 | 0.00 | 0.00 | 0.00 | 0.01 |
| *ΔLog2FC* | 1.36 | 0.40 | -0.70 | 0.44 | -0.74 |

| gene | *Pan3* | *Elavl4* | *Slc1a1* | *Grk2* | *Chd8* | *Dop1a* | *Psen1* | *Kcnb1* | *Shank2* | *Cacna2d2* | *Cdkn1a* | *Ddx21* | *Slc17a7* | *Kcnq3* |
|---|---|---|---|---|---|---|---|---|---|---|---|---|---|---|
| *Log2FC(totRNA)* | 0.00 | 0.72 | 0.16 | 0.26 | 0.06 | -0.26 | -0.08 | 0.88 | 1.04 | 3.23 | 0.15 | 0.24 | 1.90 | 2.37 |
| *ΔLog2FC* | -0.99 | -0.69 | -0.94 | -0.96 | -1.11 | -1.27 | -1.59 | -1.11 | -1.15 | -1.72 | -2.03 | -2.20 | -2.37 | -2.50 |

### FIGURE 11

**A**

| | **nascent protein (cumulative signal per cell)** | *Nmt1* | *Sgk1* | *Homer1* |
|---|---|---|---|---|
| | Ctr.1 | 1.02 | 1.25 | 1.38 |
| | Ctr.2 | 0.98 | 1.14 | 0.69 |
| | Ctr.3 | | 0.38 | 0.88 |
| | Ctr.4 | | 1.22 | 0.91 |
| | Ctr.5 | | | 1.14 |
| *sample values* | | | | |
| | *Foxg1* -OE.1 | 0.94 | 4.24 | 1.26 |
| | *Foxg1* -OE.2 | 0.72 | 2.51 | 0.91 |
| | *Foxg1* -OE.3 | 1.09 | 3.73 | 0.68 |
| | *Foxg1* -OE.4 | | 1.89 | 1.65 |
| | *Foxg1* -OE.5 | | | 1.10 |
| average | Ctr | 1.00 | 1.00 | 1.00 |
| | *Foxg1* -OE | 0.92 | 3.09 | 1.12 |
| sem | Ctr | 0.02 | 0.21 | 0.11 |
| | *Foxg1* -OE | 0.11 | 0.54 | 0.15 |
| p | | 0.30 | 0.01 | 0.28 |

**B**

| | | *sample values* | | | | | | |
|---|---|---|---|---|---|---|---|---|
| | | Ctr | | *Foxg1* -OE | | *average t4.5/t0* | | |
| **nascent Camk2b protein (cumulative signal per spot)** | | *t0* | *t4.5* | *t0* | *t4.5* | Ctr | *Foxg1* -OE | *p* |
| | *#1* | 1.14 | 0.78 | 1.12 | 0.59 | 1.00 | 0.68 | 0.00104 |
| | *#2* | 0.97 | 0.78 | 1.02 | 0.52 | | | |
| | *#3* | 0.89 | 0.77 | 0.86 | 0.47 | | | |

| | | Ctr | | *Foxg1* -OE | | *average t6/t0* | | |
|---|---|---|---|---|---|---|---|---|
| **nascent Fmr1 protein (cumulative signal per spot)** | | *t0* | *t6* | *t0* | *t6* | Ctr | *Foxg1* -OE | *p* |
| | *#1* | 1.16 | 0.47 | 1.21 | 0.91 | 1.00 | 1.60 | 0.02600 |
| | *#2* | 0.66 | 0.36 | 1.46 | 0.86 | | | |
| | *#3* | 0.71 | 0.30 | 0.81 | 0.85 | | | |
| | *#4* | 0.88 | 0.61 | 0.59 | 0.66 | | | |
| | *#5* | 1.54 | 0.78 | 1.06 | 0.33 | | | |
| | *#6* | 1.06 | 0.20 | 0.88 | 0.75 | | | |

### FIGURE S1

| | Fig.2A left, trapRNA | *Foxg1* |
|---|---|---|
| *sample values* | Ctr.1 | 0.83 |
| | Ctr.2 | 0.89 |
| | Ctr.3 | 1.42 |
| | Ctr.4 | 1.09 |
| | Ctr.5 | 1.02 |
| | Ctr.6 | 0.75 |
| | *Foxg1* -OE.1 | 3.87 |
| | *Foxg1* -OE.2 | 4.47 |
| | *Foxg1* -OE.3 | 3.79 |
| | *Foxg1* -OE.4 | 3.18 |
| | *Foxg1* -OE.5 | 4.81 |
| | *Foxg1* -OE.6 | 3.19 |
| *average* | Ctr | 1.00 |
| | *Foxg1* -OE | 3.89 |
| *sem* | Ctr | 0.10 |
| | *Foxg1* -OE | 0.27 |
| *p* | | 0.000001 |

| | Fig.2A left, totRNA | *Foxg1* |
|---|---|---|
| *sample values* | Ctr.1 | 0.99 |
| | Ctr.2 | 0.88 |
| | Ctr.3 | 1.11 |
| | Ctr.4 | 1.02 |
| | *Foxg1* -OE.1 | 4.33 |
| | *Foxg1* -OE.2 | 4.34 |
| | *Foxg1* -OE.3 | 3.87 |
| | *Foxg1* -OE.4 | 4.58 |
| *average* | Ctr | 1.00 |
| | *Foxg1* -OE | 4.28 |
| *sem* | Ctr | 0.05 |
| | *Foxg1* -OE | 0.15 |
| *p* | | 0.0000004 |

| | Fig.2A right, totRNA | *Foxg1* |
|---|---|---|
| *sample values* | Ctr.1 | 0.86 |
| | Ctr.2 | 1.02 |
| | Ctr.3 | 1.08 |
| | Ctr.4 | 1.04 |
| | *Foxg1* -LOF.1 | 0.70 |
| | *Foxg1* -LOF.2 | 0.07 |
| | *Foxg1* -LOF.3 | 0.55 |
| | *Foxg1* -LOF.4 | 0.61 |
| *average* | Ctr | 1.00 |
| | *Foxg1* -LOF | 0.48 |
| *sem* | Ctr | 0.05 |
| | *Foxg1* -LOF | 0.14 |
| *p* | | 0.006 |

| | Fig.3A-B, totRNA | *Foxg1* |
|---|---|---|
| *sample values* | Ctr.1 | 0.99 |
| | Ctr.2 | 0.90 |
| | Ctr.3 | 1.08 |
| | Ctr.4 | 1.03 |
| | *Foxg1* -LOF.1 | 0.24 |
| | *Foxg1* -LOF.2 | 0.25 |
| | *Foxg1* -LOF.3 | 0.34 |
| | *Foxg1* -LOF.4 | 0.30 |
| *average* | Ctr | 1.00 |
| | *Foxg1* -LOF | 0.28 |
| *sem* | Ctr | 0.04 |
| | *Foxg1* -LOF | 0.02 |
| *p* | | 0.000002 |

### FIGURE S2

**D**

| | **nascent Grin1 protein (cumulative signal per spot)** | *T0* | *T1* | *T2* | *T5* | *T10* |
|---|---|---|---|---|---|---|
| *samples values* | *#1* | 1.18 | 0.71 | 0.76 | 0.64 | 0.67 |
| | *#2* | 1.11 | 0.75 | 0.63 | 0.63 | 0.59 |
| | *#3* | 0.71 | 0.79 | 0.78 | 0.63 | 0.66 |
| | *#4* | 1.11 | 0.83 | 0.76 | 0.72 | 0.68 |
| | *#5* | 1.05 | 0.73 | | | 0.37 |
| | *#6* | 0.95 | 0.93 | | | 0.49 |
| | *#7* | 0.89 | 0.91 | | | 0.54 |
| *average* | | 1.00 | 0.81 | 0.73 | 0.65 | 0.57 |
| *sem* | | 0.06 | 0.03 | 0.03 | 0.03 | 0.04 |
